# Longevity of Implantable Silicon-ICs for Emerging Neural Applications: Evaluation of Bare Die and PDMS-Coated ICs After Accelerated Aging and Implantation Studies

**DOI:** 10.1101/2024.03.06.583769

**Authors:** Kambiz Nanbakhsh, Ahmad Shah Idil, Callum Lamont, Csaba Dücső, Ömer Can Akgun, Domonkos Horváth, Kinga Tóth, István Ulbert, Federico Mazza, Timothy G. Constandinou, Wouter Serdijn, Anne Vanhoestenberghe, Nick Donaldson, Vasiliki Giagka

**Affiliations:** Department of Microelectronics, Faculty of Electrical Engineering, Mathematics and Computer Science, Delft University of Technology, Delft, The Netherlands; Department of Medical Physics and Biomedical Engineering, University College London, London, United Kingdom; Centre for Energy Research, Budapest, Hungary; Research Centre for Natural Sciences, Institute of Cognitive Neuroscience and Psychology, HUN-REN, Budapest, Hungary; Department of Electrical & Electronic Engineering, Imperial College London, United Kingdom; UK Dementia Research Institute, Care Research and Technology Centre, London, United Kingdom; Mint Neurotechnologies Ltd, London, United Kingdom; School of Biomedical Engineering & Imaging Sciences King’s College London; London Institute of Healthcare Engineering – LIHE, London, United Kingdom; Department of Neuroscience Erasmus Medical Center, Rotterdam, The Netherlands; Department of System Integration and Interconnection Technologies, Fraunhofer Institute for Reliability and Microintragration IZM, Berlin, Germany

**Keywords:** Implantable bioelectronics, active neural interface, integrated circuits, biofluid barriers, microelectronics packaging, PDMS

## Abstract

Silicon integrated circuits (ICs) are central to the next-generation miniature active neural implants, whether packaged in soft polymers for flexible bioelectronics or implanted as bare die for neural probes. These emerging applications bring the IC closer to the corrosive body environment, raising reliability concerns, particularly for long-term clinical use. Here, we evaluated the long-term electrical and material stability of silicon-ICs from two foundries, after one-year accelerated *in vitro* and *in vivo* animal studies. The ICs featured various custom-designed test structures and were partially PDMS coated, creating two regions on each chip, uncoated “bare die” and “PDMS-coated”. During the accelerated *in vitro* study, ICs were electrically biased and periodically monitored. Results demonstrated stable electrical performance for at least a year, suggesting that bare die ICs can function in the body for months. Despite electrical stability, material analysis revealed chemical and electrically driven degradation of the IC passivation in the bare die regions. In contrast, PDMS-coated regions revealed no such degradation, making PDMS a highly suitable encapsulant for ICs intended for years-long implantation. Based on the new insights, guidelines are proposed that may enhance the longevity of implantable ICs, significantly broadening their applications in the biomedical field.

## 1. Introduction

Since the first silicon integrated circuit (IC) was inserted in the brain for sensing neural activity^1,2^, tremendous efforts have been made in exploiting the high performance and power efficiency offered by the semiconductor industry in opening new applications in the field of healthcare: brain-computer interfaces^3–5^, flexible bioelectronics^4,6–8^, biosensors^9–12^, active silicon probes^6,13,14^, and optogenetics^15,16^.

Driven by the need for miniaturization, these emerging applications are moving away from the hermetic metal enclosures, traditionally utilized for IC protection in the body, and are opting for newly engineered thin organic and inorganic coatings^6,17–22^. This shift, however, introduces reliability risks by bringing the chip closer to the corrosive environment of the body. Silicon-ICs are liable to failure if penetrated by body fluid: field effect transistors are susceptible to mobile ions (e.g., Na^+^, K^+^) that might reach the gate oxide^23,24^, and water anywhere in the intricate sub-micron structures of the IC will facilitate corrosion and leakage currents^25,26^. Additionally, the body itself should be protected from the electrical bias voltages on the chip which could pose a hazard if the IC’s insulation is breached. With the growing interest in miniaturizing active neural implants, these reliability risks, and the uncertainty regarding the longevity of the IC have been identified as one of the main obstacles limiting the widespread clinical adoption of these emerging applications, particularly for chronic use^27–32^.

The long-term reliability of implantable ICs relies on the stability of their constituent materials. However, thus far, there has been no in-depth study evaluating the long-term stability of IC materials in the implant environment. Gaining such insight will be instrumental in guiding future IC design and material engineering for IC packaging, ultimately contributing to the development of next-generation, long-lasting active neural implants.

In this paper, we evaluated the electrical and material stability of ICs sourced from two complementary metal-oxide-semiconductor (CMOS) foundries, after long-term *in vitro* and *in vivo* studies. Our primary objectives were: 1) to determine the longevity of bare ICs when directly exposed to physiological media by identifying the degradation pathways that can lead to failure, and 2) evaluate the possibility of utilizing polydimethylsiloxane silicone rubber (PDMS) as a soft biocompatible coating, in preventing the degradations and extending the longevity of implantable ICs.

Silicon ICs are intricate multilayer structures, comprised of conducting metallization layers and insulating ceramics layers used as intermetallic dielectrics (IMD). Between the different metal layers, metal vias are used for making vertical interconnections. All layers of the IC stack are deposited onto a silicon substrate (**Figure 1**). The chemical composition of the materials used in the IC can vary depending on the technology node and the deposition processes utilized by the IC foundry^33^. For the metallization, aluminum or copper are generally used, for the vias tungsten (W), and for the insulating IMD layers silicon dioxide is commonly employed^34^. The topmost insulating layers, known as the “passivation” layers, are usually a dual layer of silicon nitride (SiNx) and silicon dioxide (SiO_2_). These layers are deposited by plasma-enhanced chemical vapor deposition (PECVD) techniques and cover the entire surface of the IC, except for the pad openings. The passivation layers have been specifically included in the CMOS process to protect the IC against ambient humidity and contamination. The sidewalls of the chip, however, are not protected by the passivation layers. For this reason, the CMOS industry has used a die seal ring (a stack of metallization encircling the die) to protect the inner circuitry from impurity ingress and to provide structural support during dicing^35^. From the bottom, the IC is protected by the ∼ 200 - 300 µm thick silicon substrate, which has high density and high barrier properties.

**Figure 1.**
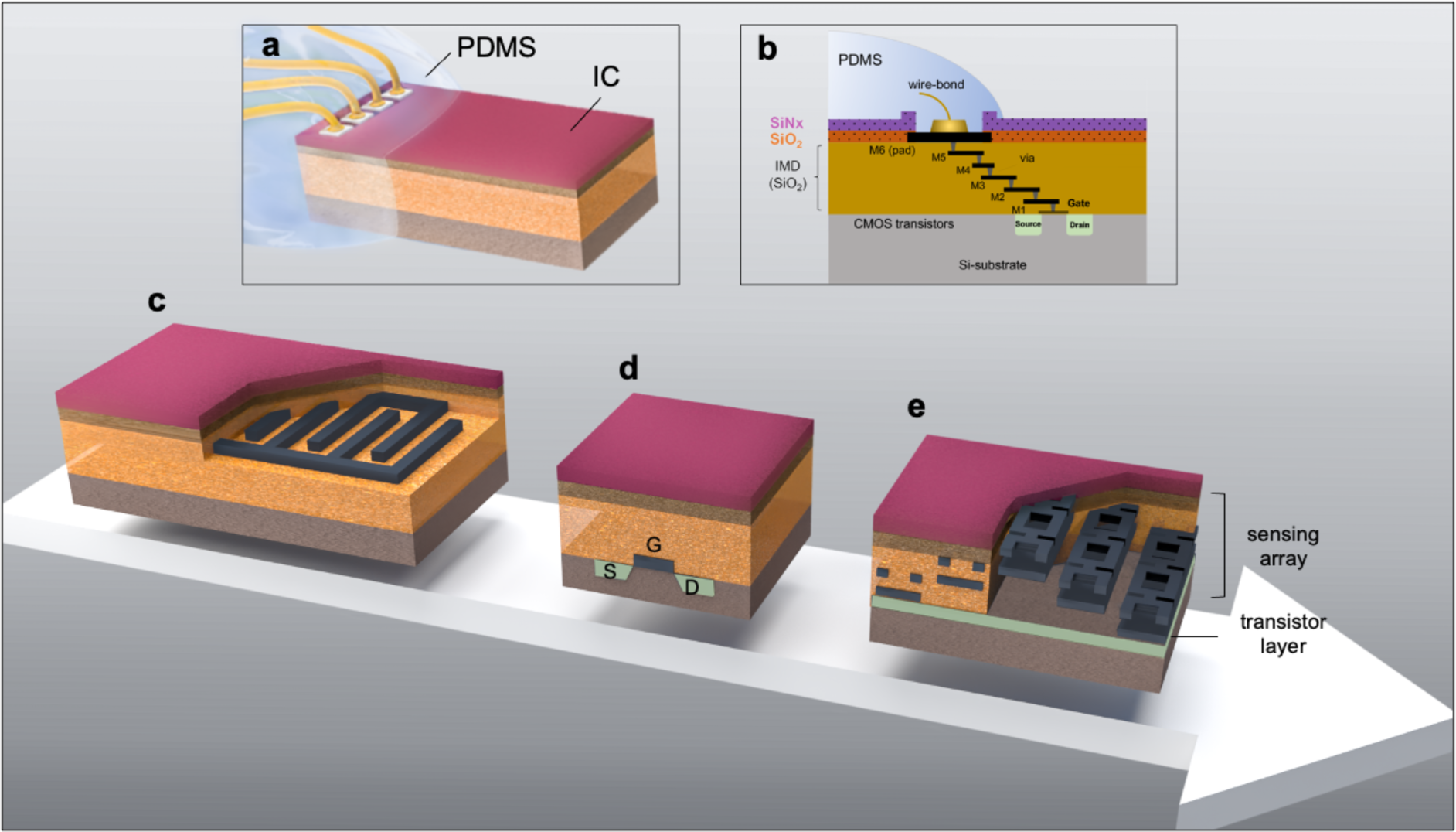
Schematic illustrations of silicon-IC test structures (dimensions not to scale). **a)** A wire-bonded IC partially coated with PDMS, covering the wire-bonds and regions of the outer IC surface, leaving most of the IC structure and sidewalls exposed. **b)** A cross-sectional schematic demonstrating the multilayer stack of a representative 6-metal CMOS process, from bottom to top: the source and drain diffusion regions implanted in the Si-substrate, metal layers 1 to 6 (M1 to M6) with M6 being the top-most metal, and where each metal layer is insulated with a SiO_2_ intermetallic dielectric (IMD), and the final passivation layers generally made of SiO_2_ and SiNx. **c) – e)** Schematic of implemented test structures in silicon-IC, from simple to more advanced with, **c)** an interdigitated capacitor (IDC) structure implemented using the top metal layers where the structure is closer to the outer surface, **d)** A planar metal oxide semiconductor (MOS) transistor with the drain, source and gate implemented in the Si-substrate boundary, making the MOS structures the deepest buried structure underneath the IMD stack and passivation layers. **e)** A dielectric sensor array with on-chip sensing circuitry for in-situ monitoring of insulation and dielectric changes. Dielectric changes between different metal layers (M6-M5, M5-M4 and M4-M3) are sensed using on-chip MOS transistor circuitry, digitized and sent off-chip in a digital bit stream.

When implanted as bare die, the chemical stability and barrier properties of the IC passivation layers play a significant role in the longevity of the IC. Both these properties are determined by the material type and the deposition processes utilized by the chip foundry. Pre-existing defects, pinholes, or nanopores have been reported for ceramic dielectric layers which could compromise their protection in wet ionic environments such as the human body^36^. PECVD silicon nitride and dioxide layers have also been reported to dissolve when exposed to wet ionic media^36–39^. When reaching interfaces, water molecules may also cause interlayer delamination in the IC stack, causing mechanical damage and failure^33,36^.

When coated in PDMS, the IC is protected from direct exposure to tissue and body fluids. Nevertheless, due to PDMS’s high moisture permeability, the IC material may still undergo degradation when implanted in the body. In our previous work^40^, we demonstrated that not all silicon-based dielectrics are equally suitable for long-term implantation as diffusing moisture through the PDMS may also penetrate the dielectric and gradually degrade its insulating and dielectric properties. Therefore, long-lasting moisture barrier properties of the IC materials are a key requirement for implantable chips intended for long-term use. In addition, for PDMS-coated ICs, strong adhesion between the IC material and PDMS is crucial for ensuring long-term electrical functionality. Strong interfacial adhesion is required to maintain electrical insulation between the wire-bonded pads^25,41^. Adhesion stability is influenced by the surface chemistry of the IC’s passivation layer and the PDMS type^25^.

In this investigation, ICs were partially coated with PDMS, intentionally leaving most of the IC passivation surface and sidewalls exposed as bare die (**Figures 1a-b**). Where bare, the intricate metallization and transistor structures within the chip are only protected by the IC’s own passivation and IMD layers. In the PDMS coated region, PDMS serves as additional protection but also insulates the wire bonds and the bonding pads through its adhesion to the IC passivation. ICs were then either subjected to an accelerated *in vitro* aging study in phosphate buffered saline (PBS) solution at 67 °C or implanted in rats.

Three test structures were implemented on the ICs: 1) interdigitated capacitors (IDC), 2) metal oxide semiconductor (MOS) transistors, and 3) a dielectric sensor array with on-chip measurement and processing circuitry (**Figures 1c-e**). Using these structures, the electrical performance of the ICs is periodically monitored throughout the accelerated *in vitro* study, contributing to the long-term evaluation of the IC materials and the applied PDMS interfaces.

Complementary to the electrical measurements, several material analysis techniques were used to evaluate and compare the degradation mechanisms in the bare die and PDMS-coated regions of the aged chips. For both regions, the impact of two aging environments (PBS solution at 67 °C and rat) was also evaluated and compared. For this purpose, for the first time, advanced material analysis techniques have been used on CMOS foundry ICs to explore the bio-chemical degradation mechanisms after their long-term exposure to the implant environment.

The study presented here advances our understanding of the electrical and material stability of silicon-ICs in the implant environment, both as bare die and when coated with PDMS. The newfound insights from this study will inform and enable the design of state-of-the-art chip-scale active neural implants for chronic implantation.

## 2. Results

### 2.1. IC design, fabrication and PDMS coating

Here, we present the custom-designed IC test structures fabricated using two commercial CMOS foundries (hereafter referred to as Chip-A and Chip-B). All ICs were partially PDMS-coated and subjected to either a year-long accelerated *in vitro* study in PBS solution at 67 °C or a year-long *in vivo* animal study in rats.

For the accelerated *in vitro* study, bare dice were glued on ceramic adaptors with pre-printed Pt/Au tracks, gold wire-bonded, and then partially coated with PDMS (**Figure 2a**, details in Experimental section). By doing so, two distinct regions are created on the same chip: 1) an exposed “bare die” region; and 2) a “PDMS-coated” region (**Figure 2b**). During the aging studies, both regions were simultaneously stressed and evaluated. On all prepared samples, the test structure under investigation was left exposed (as bare die). This approach enables the evaluation of the IC’s own passivation and dielectric layers in protecting the sensitive test structures underneath when the ICs are exposed to the *in vitro* and *in vivo* aging environment.

**Figure 2.**
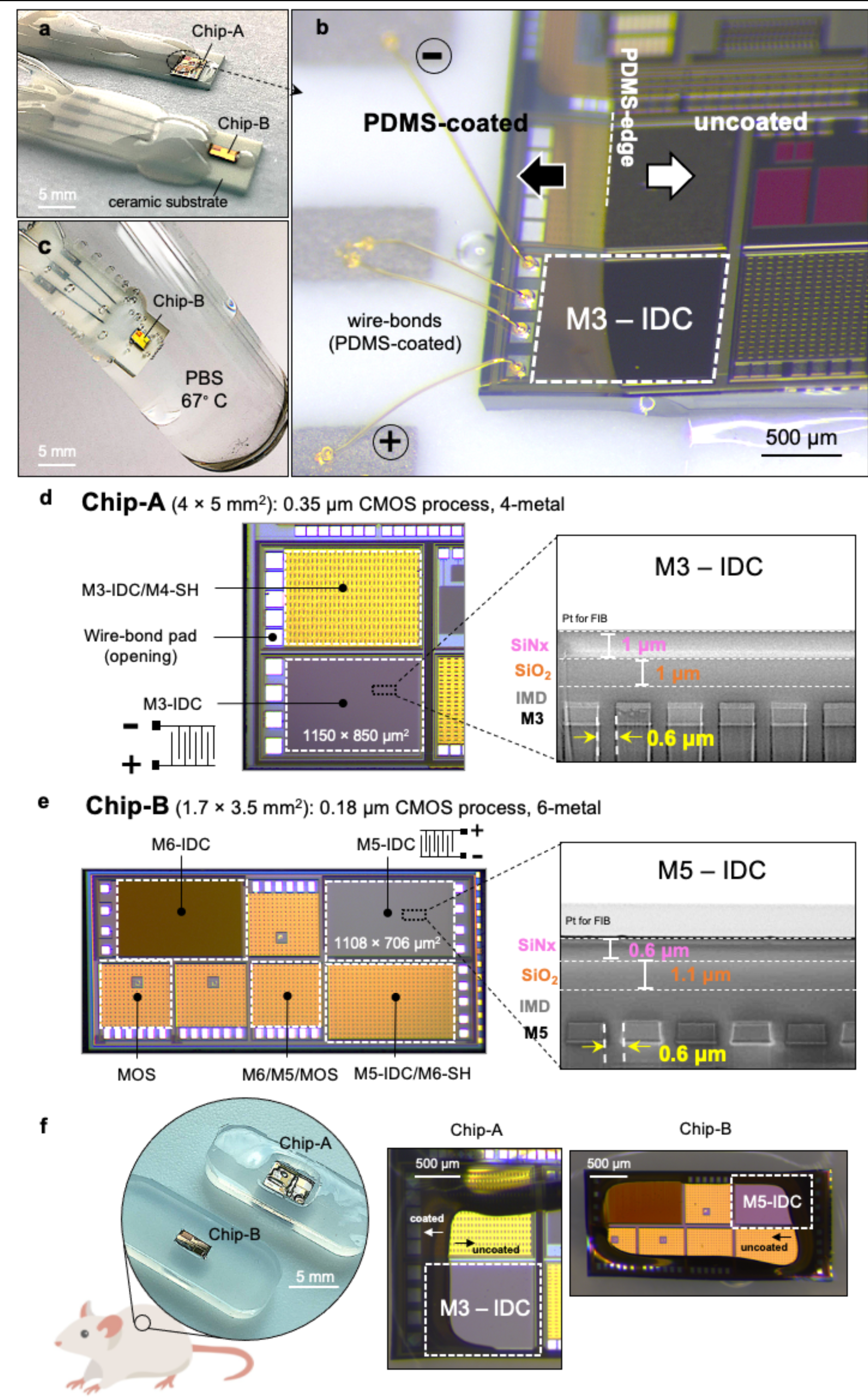
Silicon-IC test structures and preparation for long-term accelerated *in vitro* and *in vivo* aging. **a-c)** ICs wire-bonded and prepared for *in vitro* accelerated aging (voltage and temperature) in phosphate buffered saline (PBS, pH ∼ 7.4) at 67 °C, **a)** Optical image of a Chip-A and B IC placed on ceramic substrates, wire-bonded and partially PDMS-coated, **b)** Tilted optical micrograph of a representative Chip-A sample with the M3-IDC test structure wire-bonded and locally PDMS-coated, leaving most of the chip surface and sidewalls exposed, **c)** Optical image of a partially PDMS-coated Chip-B fully immersed in PBS solution at 67 °C. **d-e)** Top view optical micrographs and cross-sectional SEM images of Chip-A and B ICs with **d)** Optical micrograph of a Chip-A IC fabricated in a 0.35 µm 4-metal CMOS process with two IDC test structures (left). Cross-sectional SEM image of the M3-IDC test structure showing the interdigitated metal structures and top SiNx/SiO_2_ passivation layers. **e)** Optical micrograph of Chip-B showing various IDC and MOS transistor test structures fabricated in a 0.18 µm 6-metal CMOS process (left). Cross-sectional SEM image of M5-IDC test structure showing metal-5 and top SiNx/SiO_2_ passivation layers (right). **f)** ICs prepared for subcutaneous implantation in rat. Chip-A and Chip-B ICs positioned on soft PDMS substrates and locally coated with PDMS, covering the aluminum pads and regions of the top passivation while leaving most of the chip surface exposed to the body. Note that ICs used for the animal study are not wire-bonded.

In the PDMS-coated region, critical interfaces are created between the PDMS and the IC material (i.e., SiNx, aluminum pad, silicon substrate). Strong long-lasting adhesive bonds at these interfaces are key in preventing voids and shunt water leakage paths required to ensure the long-term electrical functionality of the IC. On our structures, the most critical PDMS interface is with the SiNx passivation at the PDMS-edge area (**Figure 2b** and **Figure S1**, Supporting Information). In this area, the PDMS-SiNx interface is directly subjected to the PBS solution. Interfacial debonding would allow lateral ingress of the solution, causing significant failure when the electrolyte reaches the electrically biased wire-bonds.

During the accelerated *in vitro* study, the IDC test structures were electrically biased either by applying a DC electrical voltage between the combs of the IDC or between the combs and the PBS solution. The main objectives for biasing the IDC structures were to: 1) electrically stress the dielectrics and passivation layers while being directly exposed to liquid PBS solution in the uncoated regions, or to diffusing moisture (gaseous water) in the PDMS-protected regions, 2) accelerate failure in the directly exposed region in case of microcracks or defects in the passivation layers, as previously reported^42–44^, and 3) in the PDMS-coated region, investigate if the combined effects of a continuous electric field and moisture would weaken the PDMS-SiN interface adhesion, potentially leading to debonding and lateral saline ingress.

A key factor in the design of the test structures is the presence or absence of a “shield (SH)”. The shield is a metal layer implemented using the top-metal of the process and designed to further protect the test structures underneath from any moisture/ion ingress (See Experimental Section). Additionally, around each test structure, a wall-of-via (WoV) is implemented which is a similar metal structure to the die seal ring but included around each individual test structure to reduce the possibility of moisture/ion penetration from the edges (sidewalls) of the test structure.

Chip-A test structures were fabricated in a 0.35 µm CMOS 4-metal process. On Chip-A two IDC test structures were implemented: 1) M3-IDC; and 2) M4-SH/M3-IDC (**Figure 2d** and **Table S1**, Supporting Information). M3-IDC (area: 1150 µm x 850 µm) contained 606 interdigitated fingers implemented in M3 (one layer below the top metal) and protected by the IMD and the top SiNx/SiO_2_ passivation layers, each being ∼1 µm thick, as presented in the cross-sectional scanning electron microscope (SEM) image in **Figure 2b**. The adjacent M3-IDC/M4-SH test structure used a similar M3-IDC interdigitated structure but included a top metal shield (∼ 2.8 μm) implemented in M4 (See cross-sectional SEM image in **Figure S2**).

Chip-B test structures were fabricated using a 0.18 µm 6-metal process and contained three types of structures: 1) IDCs; 2) MOS transistors; and 3) a dielectric sensor array (**Figure 2e** and **Table S2**, Supporting Information). **Figure S3** shows the WoV around the test structures on Chip-B.

The IDC structures on Chip-B were implemented in Metal 5 (M5-IDC) or Metal 6 (M6-IDC). One IDC structure was also implemented with a shield on top (M6-SH/M5-IDC). In the M5-IDC structure, the comb metallization is in M5 and is protected by the IMD and the top SiNx/SiO_2_ passivation layers, each having a thickness of ∼ 0.6 μm and ∼ 1.1 μm, respectively. M6-IDC is implemented in the top metal layer of the technology node and has the comb metallization protected only by the SiNx/SiO_2_ passivation layers (**Figure S4**). M5-IDC/M6-SH has the metallization on M5 and a shield layer on M6 (∼1 μm).

There are two MOS transistor structures; either with or without a shield barrier. All transistors have a W/L of 20 µm / 0.36 µm. The shield barrier for the MOS structure is a double-layer metal implemented in M5 and M6.

The dielectric sensor was diced and separated from the other test structures on Chip-B to ease wire bonding and sample preparation required for the long-term accelerated *in vitro* aging. The custom-designed sensor is capable of measuring IMD resistance values in the 10^14^ Ω range with the sensitivity to detect changes at a sub-MΩ resolution^45^. This sensor was designed and tested with two goals in mind: 1) to verify if a complex CMOS circuit could operate when fully submerged in PBS solution at 67 °C, and 2) acquire more sensitive measurements of the dielectric changes and capture a 3D mapping of the possible moisture/ion penetration pathways into the chip.

For the accelerated *in vitro* aging study, the IDC structures (n=22 from Chip-A and n=34 for Chip-B) were fully immersed in PBS solution at 67 °C while being continuously electrically biased. For biasing, three voltages were used: unbiased (n=12), 5 V DC (n=22), 15 V DC (n=22) (**Figures S5a-b**). These voltages were selected based on their relevance in neurostimulator ICs^29^. **Table S3**, Supporting Information, provides an overview of all the samples and the respective test conditions used in the study. The electrical performance of the IDCs was monitored monthly using electrochemical impedance spectroscopy (EIS) for at least 12 months. The NMOS test structures (n*=*5) and dielectric sensor (n=1) were immersed in PBS at 67 °C and monitored monthly while in soak for a 12 month duration (**Figures S5c-d**).

For the long-term *in vivo* study, a total of n=12 ICs (n=6 from each) were placed on PDMS substrates and implanted in 6 rats for a maximum duration of 12 months (**Figure 2f** and **Figure S6**, Supporting Information). Explantations were phased at 3 months (n=4), 7 months (n=4) and 12 months (n=4). While implanted, the ICs were neither powered nor electrically monitored.

The chemical composition of the IC passivation layers was characterized by depth profiling the layers using time-of-flight secondary ion mass spectrometry (ToF-SIMS) and X-ray photoelectron spectroscopy (XPS). The silicon nitride on both Chip-A and B was measured to be non-stoichiometric with a Si: 50% and N: 50% and a N/Si: ∼ 1, common for PECVD deposited SiNx layers^33^. The hydrogen content in the nitride layers was measured to be ∼ 20% and ∼15% for Chip-A and B, respectively (See **Table S4** and **Figure S7**).

The SiO_2_ passivation layer for Chip-A showed a stoichiometric layer (Si: ∼33% and O: ∼67%, O/Si of ∼ 2). For Chip-B, the silicon dioxide comprised of two oxide layers. XPS results showed both layers having a O/Si of ∼ 2 (Si: ∼33% and O: ∼67%). However, ToF-SIMS results, showed the top ∼ 100 nm oxide layer having a slightly higher [OH^-^] intensity (x2.5) than the bottom layer. The bottom SiO_2_ layer had similar [OH^-^] intensities to the Chip-A SiO_2_ passivation layer. The passivation layers contained very low carbon levels with no other detectable impurities (See ToF-SIMS depth profile results in **Figure S7,** Supporting Information).

Note that for both ICs, the chemistry of the passivation layers is uniform across the entire IC. However, the thicknesses and conformality of the passivation layers are affected when using the topmost metal layers: M4 for Chip-A and M6 for Chip-B (See **Figures S2** and **S4,** Supporting Information).

The IMD layers for both chips were analyzed using energy-dispersive X-ray (EDX) elemental mapping on the IC cross-sections. Silicon dioxide is used as the IMD material for both chips, with Chip-B IMDs being slightly fluorinated (SiOF) (**Figure S2,** Supporting Information).

Fluorination is done for more modern processes to reduce the dielectric constant of the IMD layers (from *k* ∼ 4.0 for SiO_2_ to 3.5 in the fluorinated case)^34^.

### 2.2. Electrical performance in PBS solution at 67 °C

Figure 3 gives a representative set of electrical results for ICs aged in the accelerated *in vitro* study in 67 °C PBS solution. Figure 3a presents the electrical results for M5-IDC and M6-SH/M5-IDC test structures (Chip-B), continuously biased while in soak (5 V DC). EIS results (shown as Bode plots) demonstrate stable capacitive characteristics (phase ∼ −90°) and high impedance values (|Z| @ 0.01 Hz > 10^10^ Ω) throughout the 12-month accelerated aging for n=4 samples from each test structure. Results for other tested IDC structures, from both Chip-A and Chip-B, and biased at 15V DC also showed similar stable capacitive behavior over the 12-month accelerated study (**Figure S8**, Supporting Information).

**Figure 3.**
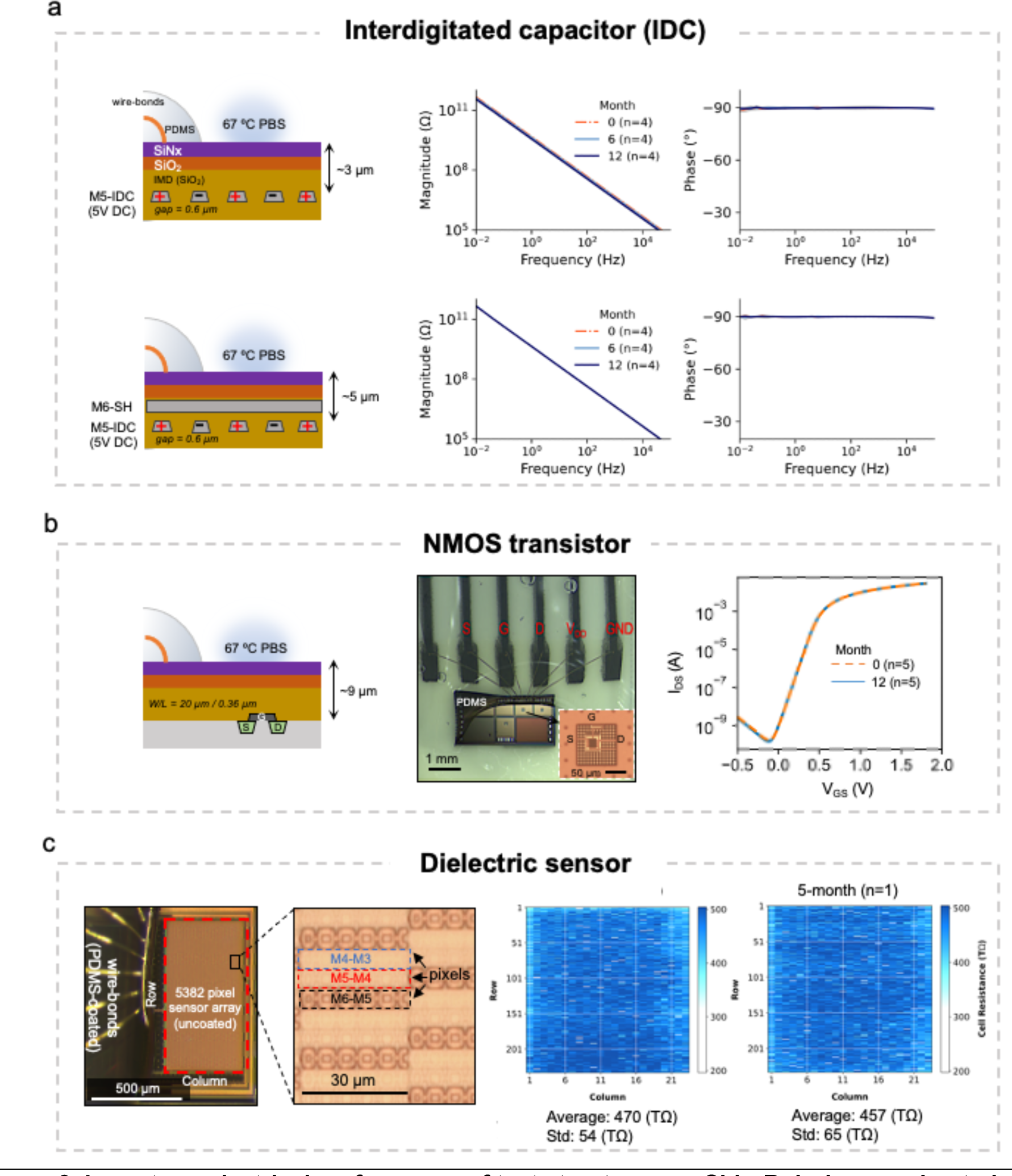
Long-term electrical performance of test structures on Chip-B during accelerated aging in PBS solution at 67 °C. **a)** Representative electrochemical impedance spectroscopy (EIS) results of M5-IDC test structures without (top, n=4), and with top-metal (M6) barrier (bottom, n=4), showing stable capacitive characteristics after one year aging. **b)** Cross sectional schematic, top view optical micrograph and V_GS_-I_DS_ transfer characteristics of NMOS transistor before and after 12-month aging (n=5, with one shielded and four unshielded MOS test structures). **c)** Top view optical micrograph of dielectric sensor with a high magnification image of the pixels (left). Measurements results of the dielectric sensor array (5382 pixels) demonstrating full functionality of the bare CMOS IC with no change in dielectric characteristics after 5 months of aging in PBS solution at 67 °C (right). All electrical measurements are performed while the ICs and its PDMS coated wire-bonds are fully submerged in PBS solution.

These results indicate three important characteristics of the applied materials and interface. Firstly, they show the stability of the dielectric and insulation properties of the passivation and IMD layers surrounding the IDC metals when fully submerged in PBS solution for at least a year while being continuously subjected to a maximum electric field of 0.25 MV/cm (at 15V DC). This is irrespective of the fluorinated silicon dioxide used in the IMD stack for Chip-B which has been reported to be unstable in the presence of moisture ^33,46^. Secondly, the stable capacitive behavior (phase ∼ - 90°) indicates that the passivation and IMD ceramic layers contain no nano-defects or pinholes which might have otherwise created parallel resistive leakage paths through the PBS solution. Thirdly, regarding the PDMS coating, these results demonstrate that PDMS effectively prevents water leakage paths by forming long-lasting adhesion to the IC’s nitride passivation. In addition, they attest to the stable electrical insulation properties of PDMS while submerged in PBS solution.

A few of our samples (14 out of 56) showed EIS irregularities (See **Table S5**, Supporting Information) which were attributed to either, 1) wire-bond corrosion (See **Figures S9 and S10**, Supporting Information), 2) stress-induced cracking in the passivation layers, or 3) poor conformality of the passivation layers. Notably, the latter two failure scenarios on both IC foundries were influenced by the use of the top metallization of the chip (**Figures S11** and **S12**, Supporting Information). The IDC structures implemented in the lower metallization (M3 in Chip-A and M5 in Chip-B) did not show any EIS irregularities.

NMOS transistors on Chip-B, both with (n=1) and without (n=4) the double shielded metal barrier, showed stable V_GS_-I_DS_ characteristics over the 1-year accelerated aging in 67 °C PBS solution (Figure 3b). For the NMOS structures without the shield metal barriers, the passivation and IMD stack are the barriers between the ionic solution and the MOS structure.

Figure 3c presents the results for the exposed dielectric sensor (Chip-B) after 5-months of aging in PBS solution at 67 °C. The top surface and three side-walls of the chip were intentionally left exposed to the PBS solution. Comparing the results at 0-month and 5-month shows minimal change when measuring between M6-M5, M5-M4 and M4-M3. Testing on this chip stopped due to an intermittent wire-bond connection failure. The results, however, demonstrate the stability of the dielectric materials on Chip-B (fluorinated silicon dioxide). More significantly, these results demonstrate the stable electrical functionality of a complex CMOS IC designed in a 0.18 µm CMOS process when submerged as a bare chip in 67 °C PBS solution.

### 2.3. Material performance: exposed vs PDMS-coated

In the previous section, the electrical stability of various IC test structures was demonstrated through a 1-year accelerated *in vitro* aging study in PBS solution at 67 °C. Despite the electrical stability, exposure to ionic fluids may still degrade the IC materials without causing discernible changes in the electrical characteristics. As identification of degradation pathways can aid in the longevity estimation of the IC, as a next step we analyzed the materials on both the exposed (bare die) and PDMS-coated regions of the aged ICs. Our investigation began by using cross sectional SEM imaging to evaluate the stability of the entire IC material stack. We were particularly evaluating the stack for any instances of interlayer delamination or intralayer degradation. Next, we examined the chemical stability and barrier properties of the top SiNx/SiO_2_ passivation layers using ToF-SIMS and XPS depth profiling. Throughout the material investigations, a comparative analysis was also conducted to understand the impact of the two aging environments on degradation: PBS solution at 67 °C and the *in vivo* animal environment at 37 °C. Finally, we investigated the impact of long-term exposure to electrical fields by focusing on a subset of IDC structures which were continuously subjected to electrical biasing (5 V and 15 V) in PBS solution at 67 °C for 12 months.

#### 2.3.1. Effect of environment: PBS solution (67 °C) and rat (37 °C)

##### 2.3.1.1. Stability of the entire IC multilayer stack

The stability of the IC’s multilayer stack was first evaluated using optical microscopy and later by SEM, examining the surface and various cross-sections across the chip. For both Chip-A and B samples, cross-sections were created using focused ion beam (FIB) at three different locations on the IDC test structures (corner, center, and PDMS-edge). Figure 4a-c shows representative cross-sections SEM images taken at the PDMS-edge of two M3-IDC test structures (from Chip-A) at two different time points, 3-month, and 10-month. Results show intact metallization and no sign of interlayer delamination in the IC stack. Higher magnification imaging, however, reveals a thinning of the top SiNx passivation in the exposed region, while no such degradation is observed for the PDMS-coated region (**Figure b-c**). At 3-month, ∼ 640 nm of the entire 1 µm SiNx passivation is remaining on the sample, corresponding to a SiNx dissolution rate of ∼ 120 nm/month for Chip-A. Dissolution was found to be uniform across the surface of the IC. Based on this dissolution rate, at 10-month, a complete loss of the entire SiNx is expected. Figure 4c shows a cross-sectional SEM image of a similar test structure aged for 10 months where only the SiO_2_ passivation is present in the exposed region, corroborating the previously calculated SiNx dissolution rate. Note that despite the complete loss of the SiNx layer at month 10, EIS results for M3-IDC structures on Chip-A remained stable up to at least 12 months of aging in PBS. After month 10, the SiO_2_ passivation is exposed and together with the intermetallic dielectric layer serve as the protective barriers on the M3-IDC structure. Similar cross-sectional SEM investigations for a Chip-B sample showed a chemical dissolution of the exposed nitride layer, but at a different rate (**Figure S13**).

**Figure 4.**
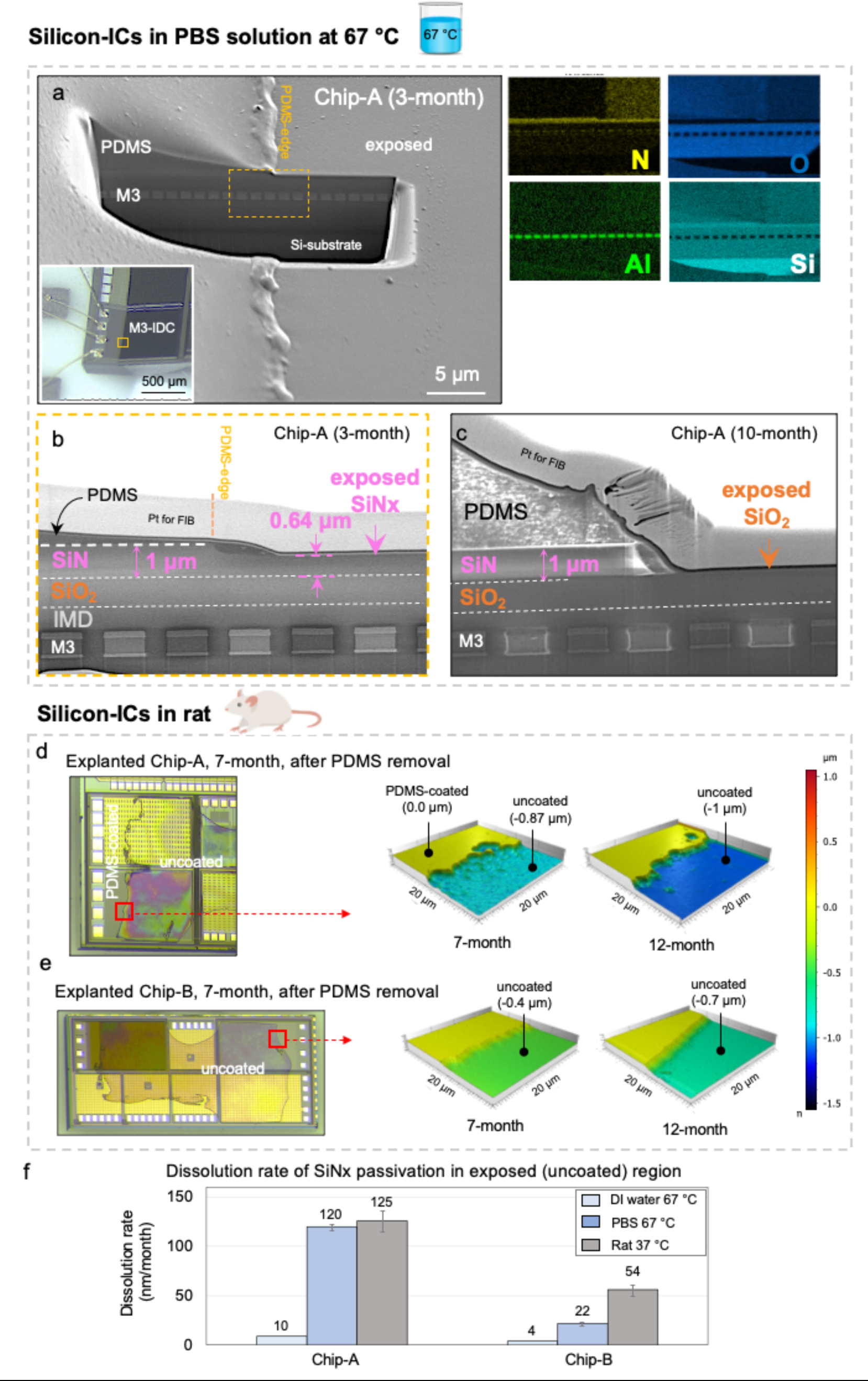
Stability of IC multilayer stack and top passivation layers after accelerated *in vitro* and *in vivo* aging. **a-c)** Representative cross-sectional SEM images of Chip-A samples after accelerated *in vitro* aging in PBS solution at 67 °C, **a)** Cross-sectional SEM image on the PDMS edge of a M3-IDC (Chip-A) structure after 6-months of aging in 67 °C PBS showing the IMD material stack from surface to silicon substrate. Inset: optical micrograph of the area used for cross-section analysis. Right, EDX elemental analysis showing intact aluminum IDC metallization. **b)** Magnified cross-sectional SEM image at the PDMS-edge showing a thinning of exposed SiNx passivation (∼350 nm) after 6-months of aging in 67 °C PBS. **c)** Cross-sectional SEM image of a Chip-A sample after 10 months with complete loss of SiNx passivation in the uncoated regions, leaving the SiO_2_ passivation exposed to the PBS solution. **d-e)** Representative optical micrograph and 3D AFM surface profiles of explanted Chip-A and B samples after PDMS decapsulation. All AFM profiles were taken within a 20 μm × 20 μm window around the former PDMS-edge (red square). AFM profiles taken after 7 and 12 months implantation in rat. **f)** Dissolution rates of uncoated SiNx passivation from Chip-A and B in different aging media (n≥2).

A group of Chip-A (n=10) and Chip-B (n=8) ICs were aged for longer than 12 months in PBS at 67 °C. After 12 months, optical microscopy on all Chip-A ICs showed gradual signs of delamination near the sidewalls of the chip. SEM investigations revealed the delamination to be between the silicon substrate and the entire IC stack (See **Figure S14**, Supporting Information). The delamination, however, was only observed on the side walls directly exposed to the PBS solution and was irrespective of electrical biasing. PDMS-protected regions remained intact. Chip-B samples, on the contrary, did not show any signs of delamination (**Figure S15,** Supporting Information).

Next, the explanted ICs was investigated using SEM cross sectional imaging and atomic force microscopy (AFM). The 3-month explanted ICs were examined using optical microscopy and cross-sectional SEM imaging (**Figures S16** and **S17,** Supporting Information). Besides SiNx dissolution on the directly exposed regions, no other degradation in the IC material stack was observed for the 3-month explanted chips. The PDMS-coated regions showed no observable SiNx dissolution. More notably, it was found that even thin PDMS layers (< 1 µm) can protect the SiNx from degradation in the body.

The 7-month and 12-month explanted samples were analyzed using optical microscopy and AFM. Samples were cleaned from tissue residues, PDMS decapsulated and later analyzed using optical microscopy and AFM on a 20 µm x 20 µm area at the PDMS-edge (Figure 4d-e and **Figure S18,** Supporting Information). Optical inspections revealed a non-uniform color on the exposed regions of the chip surface, indicating non-uniform dissolution of the passivation. The non-uniform dissolution could be due to the non-homogeneous coverage of various enzymes and tissue on the chip surface. After decapsulation, PDMS-coated regions were similarly inspected, showing no observable signs of degradation (See **Figure S18**).

AFM analysis on 7-month and 12-month explanted ICs showed a loss of the SiNx passivation (Figure 4d-e and **Figure S19**, Supporting Information). Based on the calculated dissolution rates (**Figure S20**, Supporting Information) on the 12-month explanted Chip-A samples, the 1 µm SiNx passivation layer had completely dissolved after ∼8 months, leaving the SiO_2_ passivation directly exposed to the body for ∼4 months. Despite the 4-month direct exposure, AFM results show the depth to be no less than 1 µm, indicating that the SiO_2_ passivation on Chip-A did not dissolve after 4 months of direct exposure to body environment. Similarly, for the 12-month explanted Chip-B samples, based on the dissolution rate of SiNx, after the complete dissolution of the SiNx at month ∼10, the SiO_2_ passivation was directly exposed for ∼2 months to body environment (**Figure S20,** Supporting Information). The AFM profile at 12-month, however, showed a loss of ∼700 nm indicating that in addition to the ∼ 600 nm SiNx layer, the top SiO_2_ passivation on Chip-B has experienced a ∼100 nm loss within the two months of exposure to body. As revealed in section 2.1, the SiO_2_ passivation on Chip-B contains two oxide layers with the topmost layer (∼100 nm) having a x2.5 higher [OH^-^] intensity (**Figure S7,** Supporting Information). Higher hydroxyl ions incorporated in SiO_2_ PECVD ceramics have been reported to reduce their density and chemical stability in moist environments^47^.

In terms of biocompatibility of the implanted ICs, no adverse tissue reaction or inflammation was observed for all the 11 animal models, despite the gradual dissolution of the SiNx passivation on both ICs (See **Figures S21** and **S22,** Supporting Information).

In Figure 4f, we compare the dissolution rates of SiNx passivation after exposure to various aging environments. A comparison of the dissolution rates in PBS solution at 67 °C (120 nm/month for Chip-A and 22 nm/month for Chip-B) and *in vivo* at 37 °C (∼125 nm/month for Chip-A and ∼54 nm/month for Chip-B) clearly illustrates the body’s more aggressive environment for silicon nitride dissolution compared to PBS at 67 °C.

Supporting experiments were conducted by aging the IC in de-ionized (DI) water. The lower dissolution rate of SiNx in DI water at 67 °C compared to PBS solution at the same temperature indicates that the dissolution process for SiNx is not entirely hydrolytic (water driven) but that the ions present in the solution (Na^+^, Cl^-^, PO_4-_) greatly affect the dissolution kinetics. Dissolution of thin-film ceramic material has been previously reported in literature, showing the chemical instability of these PECVD deposited ceramics when exposed to wet ionic environments.

When comparing the SiNx dissolution rates on Chip-A and B, a higher dissolution rate for Chip-A material is observed in all ageing environments. This is despite the similar N/Si for both chips (Chip-A and B both having a N: 50% and Si: 50%) as given in **Figure S7**, Supporting Information. Besides other factors such as the morphology which could impact chemical stability, the higher dissolution rate could be due to the slightly higher hydrogen (H) content (∼20% for Chip-A and ∼15% for Chip-B).

##### 2.3.1.2. Stability and barrier properties of IC passivation layers

Next, we investigated the chemical stability and ionic barrier properties of the SiNx/SiO_2_ passivation after their long-term exposure to physiological media.

The ingress of various cations (Na^+^, K^+^, Ca^+^ and Mg^+^) and anions (Cl^-^, PO_4-_, S^-^) in the exposed regions of the ICs was measured by positive and negative mode ToF-SIMS depth profiling, respectively. Depth profiles were taken from surface to ∼200 nm within the SiO_2_ passivation layer. Figure 5 presents the positive mode depth profiles of Chip-A and B ICs, explanted after 7 and 12 months *in vivo*. In the positive mode, [Si_2_N^+^] and [SiO^+^] cluster ions were used to identify the SiNx and SiO_2_ passivation layers^48^. For the 7-month explanted ICs, ∼ 100 nm and ∼ 200 nm of the SiNx passivation remained on Chip-A and Chip-B, respectively. These results support the AFM measurements showing the dissolution and thinning of the SiNx passivation (Figure 4d-e). Despite the dissolution and long-term exposure, no ingress of alkali metals is present in the remaining SiNx layer, indicating that the SiNx passivation from both IC foundries maintained its ionic barrier properties over the long-term exposure *in vivo*.

**Figure 5.**
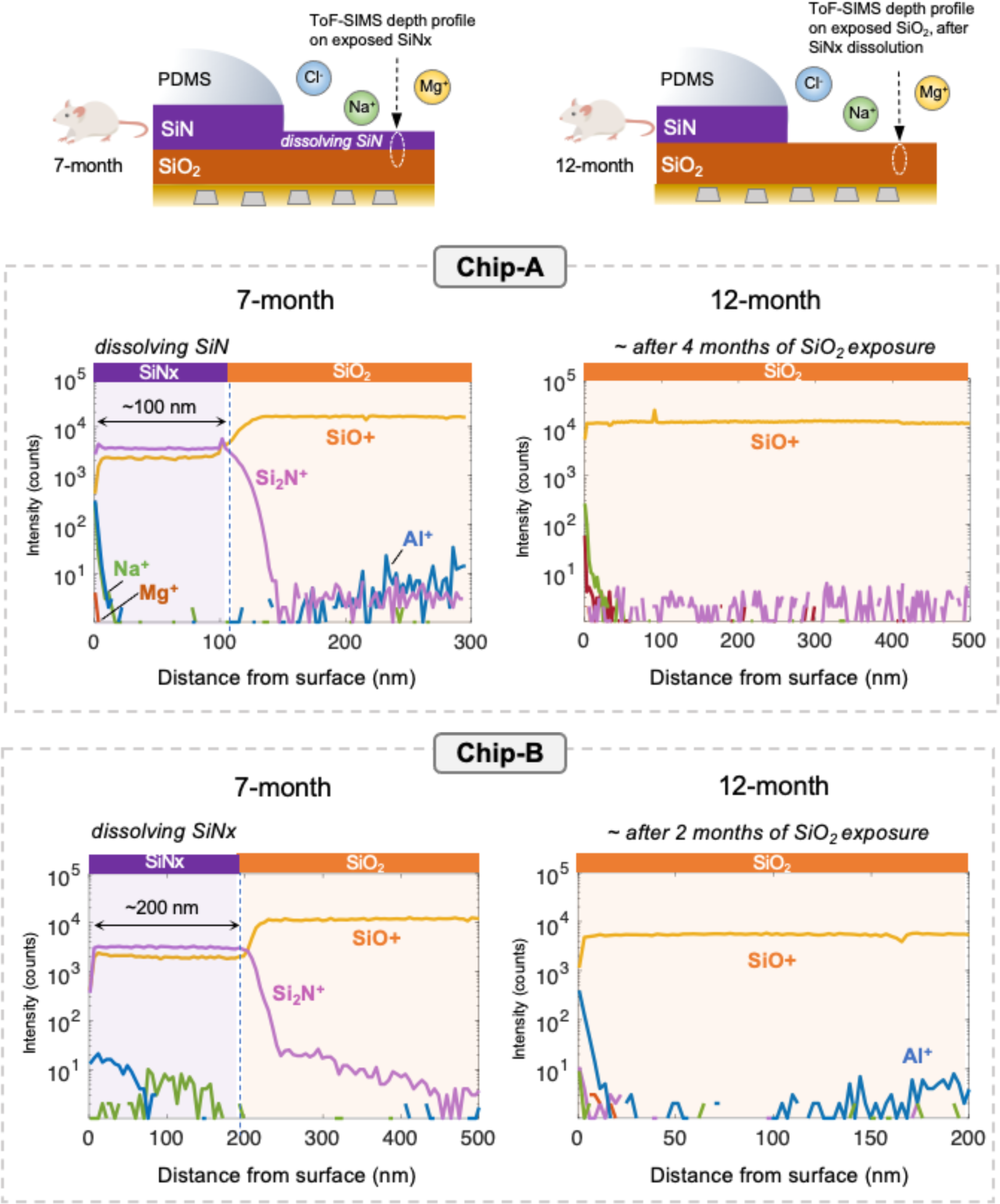
Positive mode ToF-SIMS depth profiling analyzing the ionic barrier properties of exposed passivation layers after 7 and 12 months implantation in rat. After 7 months, approximately 100 nm and 200 nm of the dissolving SiNx remained on Chip-A and B samples, respectively. Despite the dissolution, no detectable ionic penetration is found within the remaining SiNx layers. After 12 months, with the complete dissolution of SiNx, the SiO_2_ passivation is directly exposed for approximately 4 and 2 months for Chip-A and B, respectively, but no ionic species are detected within the SiO_2_ passivation layers.

On the 12-month explanted ICs, no SiNx was detected on either chip, leaving the SiO_2_ passivation directly exposed to the body environment. Based on the estimated SiNx dissolution rates as given in Figure 4f, after the complete SiNx dissolution, the SiO_2_ passivation is exposed to tissue for approximately 4 and 2 months for Chip-A and Chip-B samples, respectively. Interestingly, no ionic ingress was detected for the exposed SiO_2_ layers from either IC foundry. Negative mode depth profiles did not show any ingress of anions and contain any extra information. Therefore, they are not shown.

These results indicate that both SiNx and SiO_2_ passivation layers from the two selected silicon-IC foundries are effective barriers against ionic ingress from the body. Note that in some profiles, a slight increase in the aluminum (Al) intensity was found in the deeper depths of the SiO_2_ as the profile was getting closer the IDC metallization. Closer to the surface or in the SiNx, however, no aluminum was detected, indicating that the passivation layers act as a two-sided barrier, preventing ionic ingress into the chip, but also inhibiting the out-diffusion of metals (Al) into the body.

Next, we investigated the chemical stability of the SiNx and SiO_2_ passivation layers on the directly exposed and PDMS-coated regions of the aged ICs. It has been reported that moisture can dissociate the Si-N, Si-O and Si-Si bonds, creating silanol groups (Si-OH) in the lattice^47,49,50^. These silanol groups can have detrimental effects on ceramic dielectric layers such as increase their dielectric constant and lower their ionic barrier properties, facilitating ion penetration within the layer.

For this purpose, using negative mode ToF-SIMS depth profiling, the [SiN^-^] and [SiO_2-_] cluster ions were used to evaluate the chemical stability of the SiNx and SiO_2_ passivation layers, respectively^48^. The moisture barrier properties of these layers were also analyzed by monitoring the [OH^-^] cluster ion intensity within the depth profiles^51^. In all cases, results on aged ICs were compared to depth profile results from reference ICs (as is from foundry).

Profiling was done at three different depths: SiNx-surface (analyzed from 0 - 15 nm); SiNx-bulk (analyzed from 50 - 100 nm within the SiNx layer), and SiO_2_-bulk ( analyzed from 100 to 200 nm within the SiO_2_ passivation layer). Surface analysis (0 - 15 nm) was done with sub-nanometre (0.139 nm) step sizes using a low energy ion gun (See Experimental). All bulk profiles were collected using ∼1 nm step sizes.

The schematic in Figure 6 summarizes the findings from the depth profiling elemental analysis. In the exposed region, a thickness increase in the surface oxidation of the SiNx passivation is observed with a slight increase in ionic impurities in the oxidized layer. In the PDMS-coated region, on the other hand, only an increase in oxidation thickness was found with no increase in ionic impurities.

**Figure 6.**
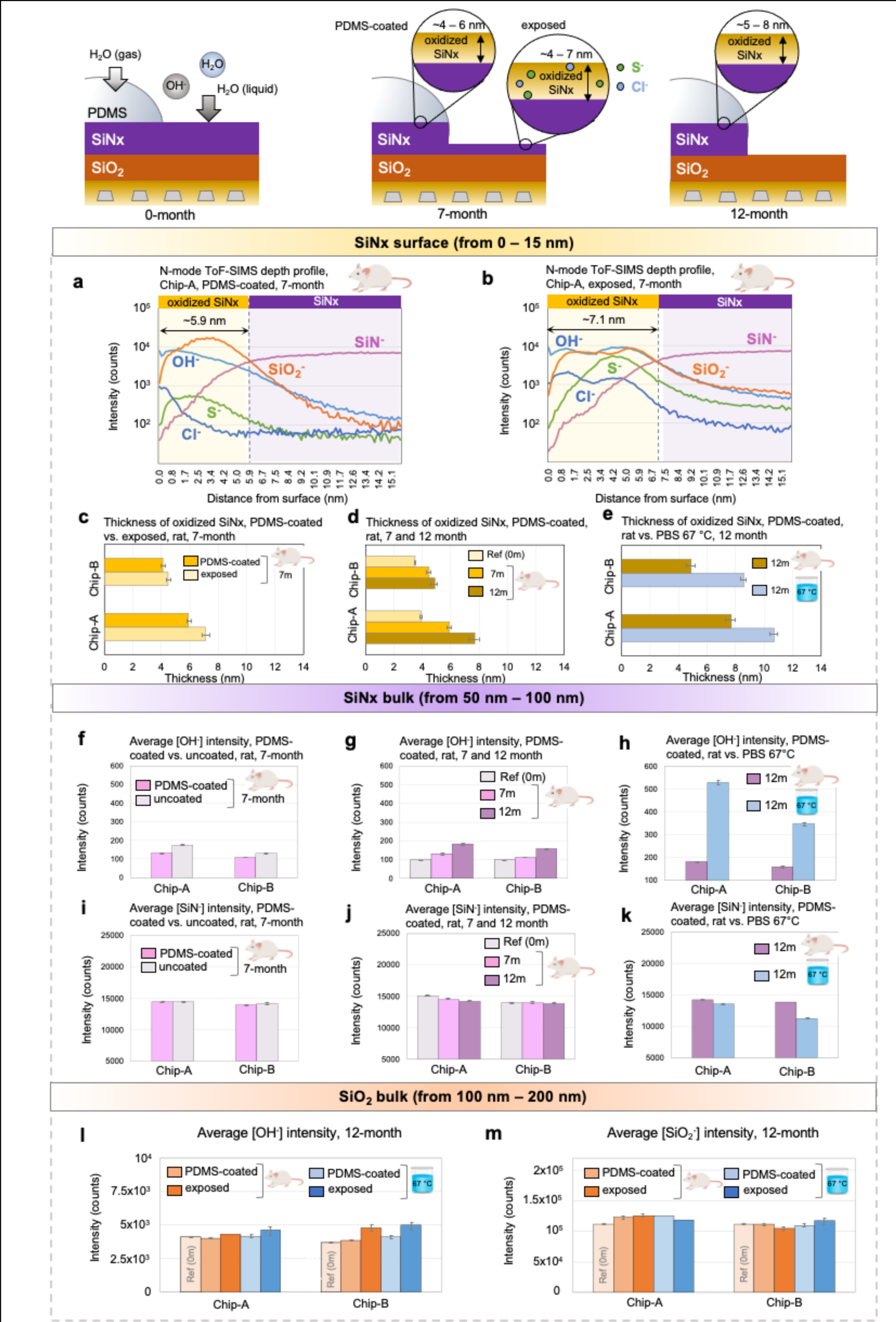
Chemical stability and moisture barrier properties of PDMS-coated and exposed passivation layers after accelerated *in vitro* and *in vivo* aging analyzed using negative mode ToF-SIMS depth profiling. **a-b)** Representative ToF-SIMS surface depth profiles from 0 - 15 nm of PDMS-coated and uncoated regions of a 7-month explanted Chip-A sample acquired with 0.1 nm step-size. Surface depth profiles show an oxidation of the SiNx passivation with higher chlorine (Cl) and sulfur (S) impurity ions in the oxidized layer for the exposed region. **c - e)** Comparing the thicknesses of the oxidized SiNx at different time points and different aging media (in 7m or 12m, m denotes month). **c)** Comparing the thicknesses in the PDMS-coated and uncoated regions after 7 months *in vivo*, **d)** Comparing the thicknesses in the PDMS-coated regions after 7 and 12 months *in vivo* with that of a dry reference sample and **e)** Comparing the thicknesses in the PDMS-coated regions after 12 months accelerated *in vitro* and *in vivo* aging. **f-n)** average [SiN^-^] and [OH^-^] intensities within the SiNx passivation bulk (averaged ToF-SIMS depth profile data from 50 - 100 nm). n-o) average [SiO_2_^-^] and [OH^-^] intensities within the SiO_2_ passivation bulk (averaged ToF-SIMS depth profile data from 100 - 200 nm). Note that in the uncoated region, at 12-month, the SiO_2_ passivation is directly exposed to body environment.

Figures 6a-b give the negative mode depth profiles of a representative Chip-A sample after 7 months of implantation in rat, comparing the first 15 nm of the SiNx passivation in the PDMS-coated and uncoated regions. In both regions, the first 5 nm demonstrated low intensities of [SiN^-^] with high intensities of [OH^-^] and [SiO_2-_]. This pattern indicates a surface oxidation of the SiNx, revealing an additional degradation mechanism affecting the SiNx passivation on the IC. In the exposed regions, a slightly higher chlorine [Cl^-^] and sulphur [S^-^] intensity was also detected in the oxidised layer. These ions could be a result of biofluids and amino acids touching the oxidized layer and diffusing into its pores^52^. In the PDMS-coated region, on the other hand, the oxidized SiNx layer showed lower [Cl^-^] and [S^-^] intensities, similar to reference level ICs. To compare the thickness of the oxidized layers in the two regions, the depth in which the [SiO_2-_] intensity is higher than the [SiN^-^] intensity is used to identify the oxidized layer in the SiNx passivation. When comparing thicknesses, a slightly thicker oxidized layer is measured in the exposed region of the IC. Surface ToF-SIMS results for a 7-month explanted Chip-B can be found in **Figure S23**, Supporting Information.

Figure 6c compares the thicknesses of the oxidized SiNx layer in the uncoated and coated regions for Chip-A and B after 7 months *in vivo*. The oxidized layer on Chip-B, for both PDMS-coated and uncoated regions, shows a thinner layer compared to Chip-A, indicating the higher density and stability of the nitride passivation on Chip-B. Figure 6d shows the thickness of the oxidized SiNx in the PDMS-coated regions over time. After 12 months of implantation in rat, ∼ 7.8 nm (0.78% of total SiNx thickness) and ∼ 4.9 nm (0.75% of total SiNx thickness) of the nitride passivation has been oxidized for the Chip-A and B ICs, respectively. Note that for reference chips (0-month) an oxide layer is already present on the SiNx surface.

Figure 6e compares the effect of the two aging environments (PBS solution at 67 °C and rat at 37 °C) on the oxidation of the nitride passivation in the PDMS-coated regions. After 12 months, ICs soaked in 67 °C PBS solution showed a thicker oxidised layer on the SiNx. These results indicate that the 67 °C accelerates the moisture diffusion and oxidation process within the SiNx layer.

The [OH^-^] and [SiN^-^] intensities within the SiNx-bulk were evaluated by averaging the intensities within the 50 - 100 nm depth of the SiN layer. Figure 6f presents the [OH^-^] intensities within the SiN-bulk for uncoated and PDMS-coated regions after 7 months of implantation in rat. Slightly higher [OH^-^] is seen in the uncoated regions that were directly exposed to tissue environment. The [OH^-^] intensity in the PDMS-coated regions was also evaluated over time in Figure 6g, indicating a gradual increase in the intensity with time. When comparing the two aging environments at month 12, a marked increase in [OH^-^] intensity is observed for the ICs soaked in PBS solution at 67 °C, resulting from the higher diffusion rate of moisture with temperature.

Figures 6i-k depict the [SiN^-^] intensity in the bulk region, showing stable intensities for all samples except those soaked in PBS solution at 67 °C. The decreased [SiN^-^] intensity observed in samples soaked in PBS solution suggests that elevated temperatures accelerate moisture diffusion and promote the breakage of Si-N bonds within the SiNx passivation layer.

Next, the SiO_2_-bulk passivation was examined by evaluating the [OH^-^] and [SiO_2-_] intensities in the layer (averaging the intensities between 100 - 200 nm). Figures 6l-m show the average intensities for the uncoated and PDMS-coated regions after 12 months exposure to both aging environments. On both Chip-A and B, a relatively stable [OH^-^] intensity is seen in the SiO_2_ layer for the PDMS-coated regions. For Chip-B a slight increase in [OH^-^] intensity is recorded for the uncoated regions. Note that for the uncoated regions, the SiO_2_ can be directly exposed to the aging environments (either PBS or tissue) after the complete dissolution of the SiNx. In the PDMS-coated regions, the SiO_2_ is always protected by both PDMS and top the SiNx passivation layer.

The [SiO_2-_] intensity within the bulk SiO_2_ layer also demonstrated stable intensities for both uncoated and PDMS-coated regions after 12 months of aging, attesting to the chemical stability of the SiO_2_ passivation layer.

#### 2.3.2. Effect of electrical field: DC electrical bias in PBS solution at 67 °C

As the final investigation, we evaluated the long-term effect of continuous electric bias on the IC materials while being immersed in PBS solution at 67 °C.

Figure 7a gives a schematic representation of a partially PDMS-coated IC applied to electrical bias (between the IDC metallization and solution) in PBS at 67 °C. During biasing, both combs of the IDC were connected to the negative (-) potential and +5 V or +15 V DC was connected to a stainless-steel rod in the PBS solution. The polarity was chosen to drive the alkaline ions (Na^+^ and K^+^) towards the IC passivation. The right schematic in Figure 7a gives an overview of the findings demonstrating the effect of the continuous biasing on the exposed and PDMS-coated regions of the chip.

**Figure 7.**
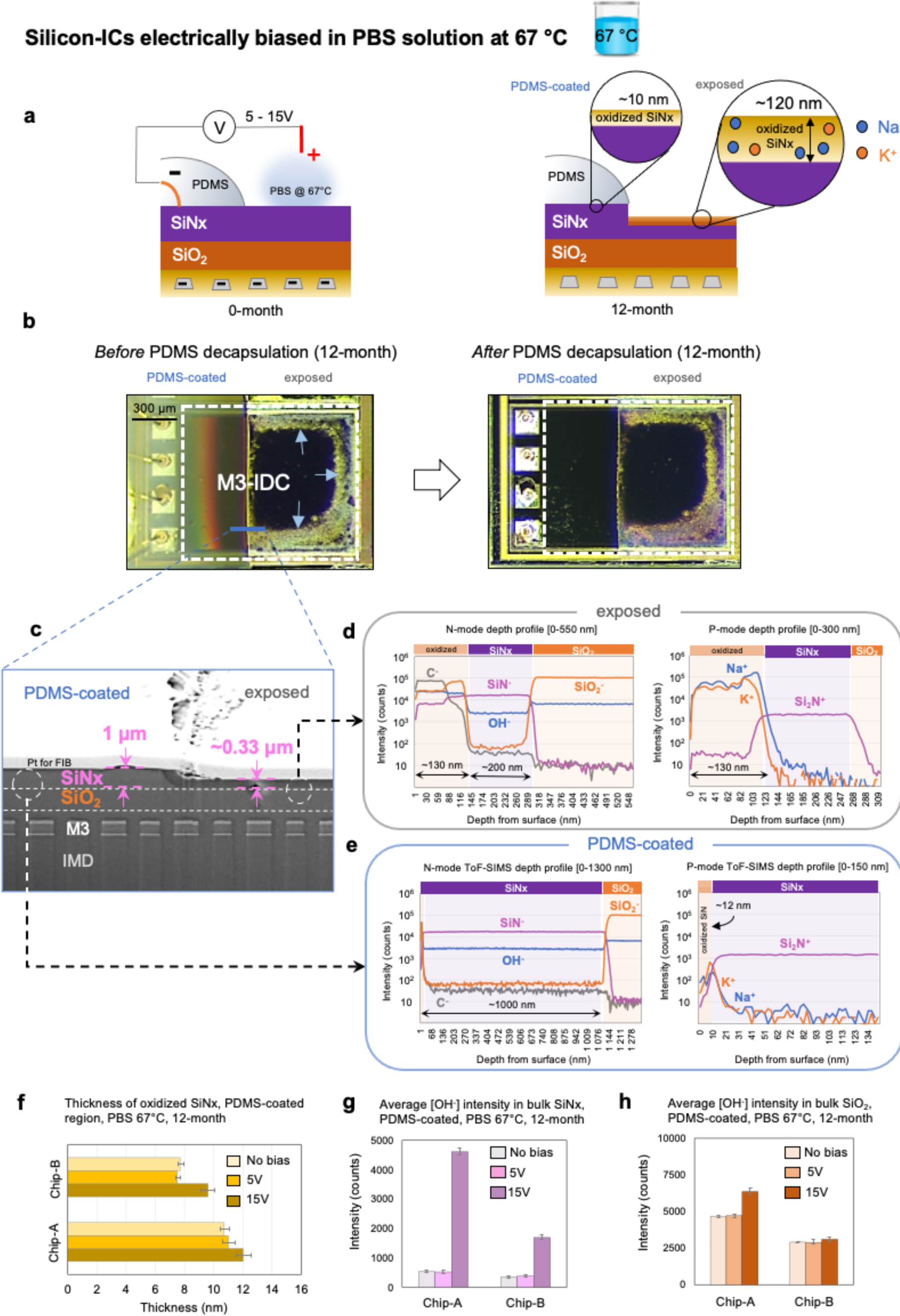
Chemical stability and ionic barrier properties of PDMS-coated and uncoated passivation layers after continuous DC biasing in PBS solution at 67 °C. **a)** Left schematic: biasing configuration with DC voltage being applied between the IDC metallization (both combs) and an electrode in PBS solution. Right schematic: the effect of the continuous biasing on passivation. **b)** Top view optical micrographs of a representative M3-IDC structure (Chip-A) sample after 12 months of continuous 15 V biasing between M3-IDC and PBS showing color change in the exposed region. Right image after PDMS decapsulation. **c)** Cross-sectional SEM image of the Chip-A sample at the PDMS-edge. **d-e)** Positive and negative mode ToF-SIMS depth profiles of the exposed and PDMS-coated regions. **f)** Thickness of the oxidized SiNx surface for Chip-A and Chip-B ICs in the PDMS-coated region after continuous biasing at 5 and 15V between IDC and PBS. For Chip-B samples, electrical biasing was applied between M5-IDC structure (negative polarity) and PBS (positive polarity). **g-h)** Average [OH^-^] intensity within the SiNx and SiO_2_ bulk (averaged from 100 - 200 nm within the layer) in the PDMS-coated region after continuous electrical biasing with 5 or 15 V DC.

Microscopically, structures continuously biased with 15 V DC voltage (n=6 from Chip-A and n=12 from Chip-B) showed signs of discoloration across their surface in the exposed region, as demonstrated in a representative M3-IDC structure from Chip-A (arrows in Figure 7b). In contrast, PDMS-coated areas of the M3-IDC structure showed no discoloration. Note that despite the observed discoloration in the exposed regions, stable capacitive behavior was recorded over the 12-month aging duration when measured between the comb structures and PBS (**Figure S8,** Supporting Information).

Figure 7c presents a cross-sectional SEM image taken at the PDMS-edge. In both the PDMS-coated and exposed regions, intact aluminum metallization was observed with no noticeable difference in the buried multilayer IC stack. However, the exposed region showed the presence of a thinned SiNx passivation layer (∼0.33 µm). This contrasts with the unbiased Chip-A sample where complete dissolution of the SiNx was observed after ∼10 months of direct exposure to PBS solution at 67 °C (Figure 4c). Figure 7c also reveals a different contrast between the thinned SiNx layer in the exposed region and the PDMS-protected region. The PDMS-protected region showed the expected thickness for the SiNx layer (1 µm).

For two biased M3-IDC (Chip-A) and two M5-IDC (Chip-B) test structures the chemical stability and barrier properties of the exposed (uncoated) and PDMS-coated regions were evaluated using negative and positive mode ToF-SIMS depth profiling. Figure 7d presents the depth profiling results for the uncoated regions on the representative M3-IDC structure (Chip-A). Negative mode depth profile results revealed a ∼120 nm oxidized layer on top of a 100 nm SiNx layer. Within the oxide layer, high intensities of [SiO_2-_], [OH^-^] and carbon [C^-^] were found. Positive mode depth profiles also showed high intensities of [Na^+^] and [K^+^] ions in the top ∼120 nm oxidized layer. Beyond the 120 nm depth, once the profile enters the remaining ∼100 nm SiNx layer, a marked drop in the impurity ions was observed demonstrating that the remaining SiNx still acts as an ion barrier. Other analyzed chips also showed similar findings.

XPS depth profiling was also used to quantify the chemical composition and ionic impurities within the oxidized layer (See **Figure S24,** Supporting Information). The chemical composition of the oxidized SiNx indicated an off-stoichiometric oxide layer for Chip-A (Si:28%, O: 58%) and Chip-B (Si:28%, O: 62%) samples. XPS depth profiling also revealed a 2 - 3% sodium (Na) incorporation in the oxidized nitride for both Chip-A and B.

In comparison to the uncoated region, depth profiling on the PDMS-coated region of the biased IDC structures showed a stable [SiN^-^] profile within the entire ∼ 1 µm SiNx passivation layer (Figure 7e). A thin (∼12 nm) oxide layer was observed on top of the SiNx layer with no ionic penetration. The low ionic penetration in the oxidized layer is due to the ionic barrier properties of the PDMS coating^53^, in this case, even in the presence of electrical fields. Nevertheless, as explained the previous section, due to PDMS’s inherent moisture permeability, a gradual oxidation of the SiNx passivation can be expected even in the PDMS-coated regions of the IC.

To examine if the applied electrical fields affected the oxidation rate of SiNx, the thicknesses of the oxidized layer in the PDMS-coated regions were measured for a group of samples which were biased with a 5 V (n=2) or 15 V (n=2) DC voltages (between IDC and PBS) during the 12-month aging study in PBS solution at 67 °C (Figure 7f). At 5 V DC, no significant change in oxide thickness was measured compared to unbiased IDC structures. However, at 15 V DC, both Chip-A and Chip-B samples showed a slightly thicker oxidized layer, suggesting that higher electrical fields may enhance the oxidation rate of SiNx in the PDMS-coated region.

Figures 7g-h gives the average [OH^-^] intensity in the bulk (100 - 200 nm) of the SiNx and SiO_2_ passivation layers. In the SiNx layer, higher [OH^-^] intensities were observed on samples subjected to continuous 15 V DC voltage as compared to 5 V and unbiased chips (Figure 7g). The [OH^-^] incorporation may result from the higher applied voltage, as higher electrical fields may introduce strain in the molecular structure of the dielectric, making the structure more susceptible to moisture attack. It should be noted that the increased [OH^-^] incorporation in the SiNx layer was exclusively detected using ToF-SIMS due to its enhanced sensitivity.

XPS depth profile results in the SiNx passivation revealed no oxygen and a N/Si ratio of ∼ 1 (similar to reference, as is from foundry ICs) for both Chip-A and Chip-B samples (**Figure S23**). The SiO_2_ passivation was similarly investigated (Figure 7i), where a slightly higher [OH^-^] intensity was observed for devices under 15 V continuous DC biasing.

## 3. Discussion and guidelines for longevity

In this work, we intended to determine the functional lifetime of bare and PDMS-coated ICs in physiological media by testing devices until failure. However, through the 1-year accelerated *in vitro* study, very few electrical IC failures were recorded on the n=62 tested samples. Therefore, complementary material evaluations techniques were performed on the *in vitro* and *in vivo* aged ICs to detect alternative degradation pathways that could help in estimating the longevity of ICs in the body.

In the exposed bare die regions, we demonstrated the long-term moisture and ionic barrier properties of the SiNx and SiO_2_ passivation layers from both CMOS foundries. Nevertheless, as bare die, the dissolution of the SiNx passivation was observed to be the main degradation pathway *in vivo* (dissolution rate of ∼125 and ∼54 nm/month for Chip-A and B, respectively). Despite the nitride dissolution, the SiO_2_ passivation underneath exhibited excellent ionic and moisture barrier properties suggesting that uncoated/unprotected ICs can maintain their functionality in the body for at least 12 months. Note that this is only the case when the top metallization is not used. When the top metal is used, due to the poor conformality of the PECVD passivation layers, the top metal will be exposed to tissue within 6-7 months of implantation causing metal corrosion (see cross-sectional SEM images in **Figure S25,** Supporting Information). Besides device malfunction, this would pose toxicity and safety risk as it would expose the body to electrical DC voltages. Therefore, for applications requiring the use of top metal (nominal or thick top metal), the use of a PDMS coating or a thin conformal coating is advised^17^.

In the PDMS-coated region, gradual surface oxidation of the SiNx passivation was observed after 1-year *in vivo* (oxidation rate of ∼3.9 and 1.4 nm/year for Chip-A and B, respectively). With this rate, it would take decades for the entire SiNx passivation to oxidize. Nevertheless, in case of total oxidation of the entire nitride passivation, the SiO_2_ passivation underneath would still serve as the second barrier protecting the electronics. Taking a conservative estimate and considering the oxidation of the SiNx to be the main degradation path for PDMS protected ICs, we believe PDMS encapsulated ICs have the potential to reach at least 10 years of functional lifetime in the body.

At first it may seem unlikely that PDMS, a polymer with low moisture barrier properties, would be suitable for long-term protection of ICs. However, as we have demonstrated in this paper, the IC’s own passivation layers have excellent moisture barrier properties. Therefore, other material characteristics of PDMS are key in achieving the long lifetime: 1) Low stress/strain ratio. Moisture-induced expansion of the polymer packaging may introduce stress on the IC’s ceramic layers^36,54–56^. In fact, shear stress cracking of the IC passivation has been reported to be one of the main failure mechanisms for epoxy packaged ICs^35^. The soft and compliant properties of PDMS, on the other hand, make it an ideal polymer for packaging implantable ICs. 2) Long-term biostability. In addition to its optimal biocompatibility, due to the unique properties of its siloxane backbone, PDMS is one of the most biostable medically used polymers^57^. *In vivo* degradation of other medical grade polymers, e.g., parylene C, have been reported through processes such as peeling or cracking^58^, which would allow direct tissue exposure to the IC material. 3) Long-term underwater adhesion to the IC surfaces, especially to the nitride passivation. Other polymers like epoxy and polyimide that are typically used for IC packaging have been reported to have weak bonds to IC passivation in moist environments.

For emerging implantable applications incorporating an IC, selecting a CMOS process suitable for chronic implantation requires extensive and long-term testing often not within the scope of academic research groups. Therefore, we encourage researchers to use the technology nodes evaluated in this work when designing their ICs. In smaller CMOS technology nodes (90 nm and smaller), significant changes in the IC materials have been made. Most importantly, for reducing resistance and crosstalk, the aluminum metallization has been replaced by copper, and for the IMD layers, new porous dielectric materials such a silicon oxy carbide (SiOC) have been adopted^33,34^. Various investigations have reported the instability of these porous dielectrics in the presence of moisture^47,59,60^. Therefore, if smaller technology nodes are to be used for implantable applications, accelerated testing in combination with the methodology used in this work can be a useful tool in evaluating the stability of the IC material.

Accelerated testing has been extensively investigated as a model to estimate the longevity of novel implantable devices. In section 2.2.2.2, we show that in the regions protected by PDMS, elevated temperatures could be used as a model to accelerate moisture penetration and degradation. However, for the exposed bare die regions of the ICs, it was found that PBS solution is not an ideal model to mimic the complex environment *in vivo* as a ∼ 20 times higher nitride dissolution rate was observed in the body environment compared to the estimated dissolution rate in PBS solution at 37 °C (assuming an Arrhenius acceleration factor from 37 °C to 67 °C). Higher *in vivo* dissolution rates for silicon-based dielectrics have been previously reported^38,39,61^. However, for the first time in this work, we compared the chemistry of the first few nanometers (0 - 5 nm) of the dissolving SiNx layer using ToF-SIMS analysis. Results revealed that all aged ICs create an off-stochiometric oxidized (SiOx) layer on top of the SiNx, with the ICs exposed to PBS solution creating a thicker oxidized layer.

ICs exposed to the body, on the other hand, created a thinner oxide layer with lower [SiO_2-_] intensity, making the Si-N bonds more readily available for cleavage and dissolution. The theory of nitride dissolution rate relying on the off-stochiometric oxidation reactions has been proposed before^62^ and has been experimentally demonstrated in this investigation. The thinner oxidized layer on the SiNx passivation could be due to a similar mechanism observed for corrosion of titanium metal implants^63^, where the proteins and enzymes in the body inhibit the surface oxidation of the metal, allowing more bulk metal to be available for corrosion.

In this work, through the 1-year accelerated aging study, no change in the electrical performance of both metal shielded and unshielded test structures was found. For longer applications, however, we believe the use of a top shield and extra wall-of-vias (WoV) around the IC may increase the longevity of the chip by creating a metal barrier around the entire IC structure. Especially for implantable IC fabricated using smaller CMOS technology nodes incorporating porous dielectric layers, the use of a metal cage could be explored as a solution to extend the longevity of the chip. In addition, a simplified version of the dielectric sensor used in this study could be employed for implantable ICs fabricated in smaller CMOS technology nodes to detect early moisture ingress.

In the accelerated *in vitro* study, corrosion of interconnect wire-bonds was found to be the cause of failure for some of the devices. Corrosion is believed to be driven by moisture and galvanic reactions between the gold wire-bond and aluminum pads, causing the corrosion of the pad. Non-bonded aluminum pads showed no signs of corrosion (**Figure S9**), mainly due to the presence of a native oxide layer on Al which has been reported to create strong interface adhesion bonds with PDMS^54^. As a mitigation, for PDMS encapsulated wire-bonded ICs, we propose the use of aluminum wire-bonds instead of gold as it can prevent the galvanic corrosion and result in a stronger adhesion between the wire-bonds and PDMS.

As demonstrated in this work, the longevity of implantable ICs greatly relies on the top passivation layers. Any stress-induced cracking in the passivation during packaging and post processing could undermine the protection offered by these layers. Die thinning, for example, has been explored as a technique to enable die embedded flexible bioelectronics^64,65^. This technique, however, may introduce stress and cracks in the passivation dielectrics. Therefore, as a future investigation, one can investigate how thin the ICs can be made without compromising the barrier properties of the IC’s passivation layers.

The effect of continuous electric fields on the stability of IC materials was investigated in a worst-case condition by applying a DC bias voltage between the IDC metals and PBS solution. For the first time, using ToF-SIMS analysis we identified enhanced [OH^-^] incorporation in the SiNx and SiO_2_ passivation layers for ICs exposed to 15 V DC in comparison to unbiased ICs. For lower voltages typically used for implantable ICs (5 V), the applied bias was found to have no long-term effect on the stability of the passivation layers. Therefore, for implantable ICs requiring higher voltages (>15 V DC) for functionality, further investigations are recommended in determining the long-term stability of the materials and interfaces.

In our complementary investigation [Nick’s EIS paper], the usefulness of these guidelines is validated by demonstrating ultra-long lifetimes for PDMS-coated ICs.

## 4. Conclusions

Ensuring the longevity of the IC in body will be a key challenge in the development of next generation miniaturized neural implants. The work presented here outlines a comprehensive methodology for evaluating the longevity of silicon-ICs. Evaluation results for IC fabricated by two CMOS foundries demonstrated the excellent moisture and ionic barrier properties of the SiNx/SiO_2_ passivation layers on the ICs, maintaining the ICs electrical functionality for at least a year, even when exposed to PBS solution as bare die. Exposure of the ICs, however, resulted in the material degradation of the IC material, with the most prominent degradation being the dissolution of the nitride passivation. PDMS coating, on the other hand, was shown to prevent the observed degradation, making it a promising soft biocompatible encapsulant for multi-year implantations. The authors believe the collected insights in this work are key in making the next step in engineering long-lasting active neural implants.

## 5. Experimental Section

### Fabrication of silicon-IC test structures

Details of the fabricated test structures are given in the main text. Data of exact technology nodes are available upon request.

### Sample preparation and PDMS coating for the long-term accelerated *in vitro* aging study

For the long-term accelerated *in vitro* aging, detailed sample preparation can be found elsewhere^40^. In summary, all ICs were glued to ceramic (alumina) adaptors with thick-film Pt/Au tracks. The adaptors were soldered to Teflon-coated stainless-steel wires. Subsequently, the chips were mounted and wire-bonded to the adaptor using a 30 µm diameter gold wire. Before wire-bonding, samples were placed in a plasma chamber (RF power 400 W, 80% Argon, 20% Oxygen) for 180 seconds to surface activate the aluminum pads of the IC for optimal wire-bonding. To insulate the metal tracks on the ceramic substrates and the connecting wire-bonds to the IC, PDMS (DOW CORNING 3140) was applied locally using a dispenser. Prior to PDMS coating, samples were activate using UV-ozone for 15 minutes for optimal PDMS adhesion. PDMS curing was done according to the manufacturer’s guidelines. The ceramic adaptors were subsequently connected to bungs. All samples were fully immersed in phosphate buffered saline (PBS) solution and aged in a dedicated apparatus^66^ (See **Figure S5**).

### Sample preparation and PDMS coating for the long-term *in vivo* animal study

For animal studies, the chips were placed on 3 mm thick, soft PDMS substrates fabricated from medical grade silicone rubber (MED2-4213, NuSil). For fabricating the PDMS substrates, a custom-made Polytetrafluoroethylene (PTFE) mold was filled with uncured PDMS which was then partially cured in the oven at 100 °C for 30 minutes. At the same time, for optimal PDMS adhesion, the ICs were UV-ozone pre-treated for 15 minutes with their top passivation facing the ozone lamp. The ICs were placed in the center of the semi-cured PDMS substrate. After positioning, all aluminum pads and parts of the IC passivation were coated with the same medical grade PDMS (MED2-4213) using a dispenser. All samples were then fully cured in the oven at 100 °C for two hours.

### Animals, implantation, and explanation procedures

All experiments were performed according to the EC Council Directive of September 22, 2010 (2010/63/EU), and all procedures were reviewed and approved by the Animal Care Committee of the Research Centre for Natural Sciences and by the National Food Chain Safety Office of Hungary (license number: PE/EA/1253-8/2019). The implantation procedures were carried out as follows. Wistar rats (n=12; weight, 315.5 ± 59.6 g at the initiation of the treatment; 3 females, 9 males) were anesthetized by an intraperitoneal injection of a mixture of ketamine (75mg/kg of body weight; CP-Ketamin, Produlab Pharma B. V., Raamsdonksveer, The Netherlands) and xylazine (10mg/kg of body weight; CP-Xylazin, Produlab Pharma B. V., Raamsdonksveer, The Netherlands). A body temperature of 37 °C was sustained with the aid of an electric heating pad connected to a temperature controller (Supertech, Pécs, Hungary).

Prior to implantation, samples were sterilized by immersing it in isopropyl alcohol for at least 20 min, followed by washing it with a continuous stream of distilled water for 2 min. In order to reduce the number of animals used, each animal was subcutaneously implanted with two ICs: Chip-A, on the right, and Chip-B, on the left side of the back close to the neck. The incision was closed using interrupted sutures followed by standard combined postoperative analgesic regimen.

Samples were explanted at three time points: months 3, 7 or 12. Before explantations, rats were initially anesthetized in the same way as described above. After anesthesia induction, the animals were overdosed with isoflurane (5% in 100% oxygen) until breathing stopped. Before explantations, the skin around the implants was examined for any inflammation. After long-term implantation, a tissue pocket was formed around each sample. To analyze the tissue pocket, the samples were removed from the pocket. The pocket was immersed in 4% paraformaldehyde solution for 24 hours, then washed in 0.1M PBS and stored in PBS. The tissue samples were then sectioned and stained with hematoxylin and eosin stain, and photomicrographs are taken for tissue analysis. Explanted microchips were carefully rinsed with DI water, blow dried and stored in a dried condition for a week.

### Electrical characterization of IDC structures: Electrochemical impedance spectrometry (EIS)

EIS was conducted with a Solartron Analytical Modulab XM, Potentiostat, Frequency Response Analyzer and Femtoammeter (**Figure S5**). For low-current measurements, the samples were placed in a dedicated Faraday cage with the shield of the Solartron coaxial cable connected to the cage. All EIS measurements, unless stated otherwise, were performed between the interdigitated combs in a frequency range of 10 mHz to 100 KHz using a 100 mV (RMS) sinusoidal voltage drive.

### Electrical characterization of transistor structures

The transistor structures were characterized using a HP4145AB parameter analyzer, which monitored the transistor drain-source current (I_DS_) over a range of gate-source voltages (−0.5 V < V_GS_ < 1.8 V) while having the V_DS_ at 1.8 V DC.

### PDMS decapsulation

PDMS decapsulation was performed post-aging to analyze the IC materials protected by PDMS. Decapsulation was done in two steps: 1) gently removing the excess PDMS material using a scalpel without damaging the ICs, 2) dissolving the remaining PDMS material using a PDMS solvent (DOWSIL™ DS-2025). The first step of removing the excess PDMS on the samples would reduce the required exposure time to the solvent. The samples were, therefore, placed in the solvent for 30 minutes and then gently rinsed in acetone, IPA and DI water and finally blown dried. All samples used for surface analysis were prepared in a similar fashion.

### IC multilayer stack characterization: Scanning electron microscopy (SEM) and energy-dispersive X-ray (EDX)

For SEM, samples were coated with a thin (10 - 20 nm) evaporated carbon layer. The carbon coated samples were investigated in a Thermo Scientific SCIOS2 system equipped with a Pt deposition gun filled with Me3PtCpMe and a Sidewinder gallium liquid metal ion source (LMIS) for focused ion beam (FIB). Prior to cross sectioning, Pt was deposited in-situ in order to properly develop the structure of the cross-sectional images. EDX was done with an Oxford Xmax 20 SDD (silicon drift detector). The applied Ga milling parameters were 30 keV and 15-30 nA, depending on the trench dimensions for coarse milling. Later, the cross-section surface was polished with 30 keV, 5 nA.

### Surface characterization using AFM

The Bruker Icon was used in tapping mode and peak-force QNM mode. After PDMS decapsulation, two regions on each IC, each having an area of 20 µm x 20 µm, on the PDMS-edge was scanned.

### Chemical composition of IC passivation layers: X-ray photoelectron spectroscopy (XPS) depth profiling

XPS analysis was carried out in a Quantera Hybrid SXMtm from ULVAC-PHI. The measurements were performed using monochromatic AlKα-radiation (25 Watt) and a take-off angle of 45°. At this angle the information depth during surface measurement is approximately 7 nm. A spot size of 100 µm in diameter was used for the analyses. First, at the surface, a survey scan was recorded. Subsequently, in the depth profiles, accurate narrow-scans of Si, N, O, C, Al, Na, Cl and F have been measured for quantification. Standard sensitivity factors were used to convert peak areas to atomic concentrations. For Chip-A and Chip-B ICs, XPS analysis was done on the M3-IDC and the M5-IDC test structures, respectively, as they contained flat surfaces with no microtopography.

### Chemical composition and barrier property of IC passivation layers: Time-of-flight mass spectrometry (ToF-SIMS) depth profiling

ToF-SIMS analyses were performed using an Ion-ToF ToF-SIMS IV instrument. The instrument was operated in both positive and negative mode using 2 keV O^2+^ and 2 keV Cs^+^ sputtering ions, respectively. At negative mode, sputtering was carried out using the 2 keV Cs^+^ primary ion beam to enhance the detection level of the electronegative species [O^-^], [OH^-^] and [H^-^]. The sputtered area was 250 x 250 µm^2^ and the analyzed area was 50 x 50 µm^2^ centered in the sputtered area. The analysis was done with a beam of 25 keV Bi^+^ ions. To increase the sensitivity for lighter elements like hydrogen [H^-^], all samples were stored in the instrument under ultra-high vacuum (<10^-9^ mbar) for 64 hours prior to analysis. Similar to XPS, all ToF-SIMS measurements were performed on the M3-IDC and the M5-IDC test structures for Chip-A and Chip-B, respectively, where the surface of chip is flat. The depth scale has been calibrated using known SiNx reference samples and has a measurement error of ∼ 10% (including a possible systematic error due to differences in stoichiometry). Shallow depth profiles (down to ∼20 nm) have been measured with 500 eV Cs^+^ ions for better depth resolution. Given the high sensitivity of ToF-SIMS compared to XPS (ppm to ppb for some species vs. 0.1–1.0 at.% for XPS), certain elements could be identified but not quantified. For this reason, all data on aged ICs have been compared to the levels measured on reference (as is from foundry) samples.

## Acknowledgments

The authors would like to thank Jurgen van Berkum and Jeannette Smulders from Eurofins, Eindhoven, The Netherlands for their support in ToF-SIMS and XPS analysis. For SEM analysis we would like to thank Mr. Levente Illés.

## Bibliography

1. Wise, K. D., Anderson, D. J., Hetke, J. F., Kipke, D. R. & Najafi, K. Wireless implantable microsystems: High-density electronic interfaces to the nervous system. in Proceedings of the IEEE vol. 92 (2004).

2. Drake, K. L., Wise, K. D., Farraye, J., Anderson, D. J. & BeMent, S. L. Performance of Planar Multisite Microprobes in Recording Extracellular Single-Unit Intracortical Activity. IEEE Trans Biomed Eng 35, (1988).

3. Rapeaux, A. B. & Constandinou, T. G. Implantable brain machine interfaces: first-in-human studies, technology challenges and trends. Current Opinion in Biotechnology vol. 72 Preprint at 10.1016/j.copbio.2021.10.001 (2021).

4. Musk, E. An integrated brain-machine interface platform with thousands of channels. J Med Internet Res 21, (2019).

5. Kim, S. et al. Integrated wireless neural interface based on the Utah electrode array. Biomed Microdevices 11, (2009).

6. Verplancke, R. et al. Development of an active high-density transverse intrafascicular micro-electrode probe. Journal of Micromechanics and Microengineering 30, (2020).

7. Vásquez Quintero, A., Arai, R., Yamazaki, Y., Sato, T. & De Smet, H. Near-Field Communication Powered Hydrogel-Based Smart Contact Lens. Adv Mater Technol 5, (2020).

8. Khan, Y., et al. A New Frontier of Printed Electronics: Flexible Hybrid Electronics. Advanced Materials vol. 32 Preprint at 10.1002/adma.201905279 (2020).

9. Eversmann, B. et al. A 128 × 128 CMOS Biosensor Array for Extracellular Recording of Neural Activity. in IEEE Journal of Solid-State Circuits vol. 38 (2003).

10. Hu, K., Arcadia, C. E. & Rosenstein, J. K. A large-scale multimodal CMOS biosensor array with 131,072 pixels and code-division multiplexed readout. IEEE Solid State Circuits Lett 4,(2021).

11. Khan, S. M., Gumus, A., Nassar, J. M. & Hussain, M. M. CMOS Enabled Microfluidic Systems for Healthcare Based Applications. Advanced Materials vol. 30 Preprint at 10.1002/adma.201705759 (2018).

12. Datta-Chaudhuri, T., Abshire, P. & Smela, E. Packaging commercial CMOS chips for lab on a chip integration. Lab Chip 14, 1753–1766 (2014).

13. Ramezani, R. et al. On-Probe Neural Interface ASIC for Combined Electrical Recording and Optogenetic Stimulation. IEEE Trans Biomed Circuits Syst 12, 576–588 (2018).

14. Steinmetz, N. A. et al. Neuropixels 2.0: A miniaturized high-density probe for stable, long-term brain recordings. Science (1979) 372, (2021).

15. Zaaimi, B. et al. Closed-loop optogenetic control of normal and pathological network dynamics. Nature Research (2020).

16. Wu, F. et al. Monolithically Integrated μLEDs on Silicon Neural Probes for High-Resolution Optogenetic Studies in Behaving Animals. Neuron 88, (2015).

17. Jeong, J. et al. Conformal Hermetic Sealing of Wireless Microelectronic Implantable Chiplets by Multilayered Atomic Layer Deposition (ALD). Adv Funct Mater 29, (2019).

18. Song, E. et al. Transferred, Ultrathin Oxide Bilayers as Biofluid Barriers for Flexible Electronic Implants. Adv Funct Mater 28, (2018).

19. Song, E. et al. Ultrathin Trilayer Assemblies as Long-Lived Barriers against Water and Ion Penetration in Flexible Bioelectronic Systems. ACS Nano 12, 10317–10326 (2018).

20. Song, E. et al. Thin, Transferred Layers of Silicon Dioxide and Silicon Nitride as Water and Ion Barriers for Implantable Flexible Electronic Systems. Adv Electron Mater 3, (2017).

21. Xie, X. et al. Long-term reliability of Al2O3 and Parylene C bilayer encapsulated Utah electrode array based neural interfaces for chronic implantation. J Neural Eng 11, (2014).

22. Pak, A. et al. Thin Film Encapsulation for LCP-Based Flexible Bioelectronic Implants: Comparison of Different Coating Materials Using Test Methodologies for Life-Time Estimation. Micromachines (Basel) 13, (2022).

23. Edell, D. J. CHAPTER 3.2 INSULATING BIOMATERIALS.

24. Jin, X. et al. Stability of MOSFET-Based Electronic Components in Wearable and Implantable Systems. IEEE Trans Electron Devices 64, 3443–3451 (2017).

25. Vanhoestenberghe, A. & Donaldson, N. Corrosion of silicon integrated circuits and lifetime predictions in implantable electronic devices. Journal of Neural Engineering vol. 10 Preprint at 10.1088/1741-2560/10/3/031002 (2013).

26. Stellari, F., Cabral, C., Song, P. & Laibowitz, R. Humidity Penetration Impact on Integrated Circuit Performance and Reliability. in Technical Digest - International Electron Devices Meeting, IEDM vols 2019-December (2019).

27. Lacour, S. P., Courtine, G. & Guck, J. Materials and technologies for soft implantable neuroprostheses. Nat. Rev. Mater. 1, (2016).

28. Giagka, V. & Serdijn, W. A. Realizing flexible bioelectronic medicines for accessing the peripheral nerves – technology considerations. Bioelectronic Medicine vol. 4 Preprint at 10.1186/s42234-018-0010-y (2018).

29. Liu, Y. et al. Bidirectional Bioelectronic Interfaces: System Design and Circuit Implications. IEEE Solid-State Circuits Magazine 12, (2020).

30. Schiavone, G. et al. Soft, Implantable Bioelectronic Interfaces for Translational Research. Advanced Materials 32, (2020).

31. Won, S. M. et al. Recent Advances in Materials, Devices, and Systems for Neural Interfaces. Advanced Materials vol. 30 Preprint at 10.1002/adma.201800534 (2018).

32. Obidin, N., Tasnim, F. & Dagdeviren, C. The Future of Neuroimplantable Devices: A Materials Science and Regulatory Perspective. Advanced Materials vol. 32 Preprint at 10.1002/adma.201901482 (2020).

33. Interlayer Dielectrics for Semiconductor Technologies.

34. King, S. W. Dielectric Barrier, Etch Stop, and Metal Capping Materials for State of the Art and beyond Metal Interconnects. ECS Journal of Solid State Science and Technology 4, N3029–N3047 (2015).

35. Moore, T. M. & McKenna, R. G. Characterization of Integrated Circuit Packaging Materials. (Butterworth-Heinemann, 1993).

36. Schmitt, G. et al. Passivation and Corrosion of Microelectrode Arrays.

37. Kang, S. K. et al. Dissolution behaviors and applications of silicon oxides and nitrides in transient electronics. Adv Funct Mater 24, 4427–4434 (2014).

38. Maloney, J. M., Lipka, S. A. & Baldwin, S. P. In Vivo Biostability of CVD Silicon Oxide and Silicon Nitride Films. (2005).

39. Ammerle, H. H., et al. Biostability of Micro-Photodiode Arrays for Subretinal Implantation. Biomaterials vol. 23 (2002).

40. Lamont, C. et al. Silicone encapsulation of thin-film SiOx, SiOxNyand SiC for modern electronic medical implants: A comparative long-term ageing study. J Neural Eng 18, (2021).

41. Donaldson_1977_silicone_rubber_adhesives.

42. Andrew, C. & Lamont, W. Non-Hermetic Protection of Implanted Thin Film and CMOS Electronic Medical Devices.

43. Nanbakhsh, K., Ritasalo, R., Serdijn, W. A. & Giagka, V. Long-term Encapsulation of Platinum Metallization Using a HfO2 ALD - PDMS Bilayer for Non-hermetic Active Implants. in Proceedings - Electronic Components and Technology Conference vols 2020-June (2020).

44. Nanbakhsh, K. et al. Effect of Signals on the Encapsulation Performance of Parylene Coated Platinum Tracks for Active Medical Implants. in Proceedings of the Annual International Conference of the IEEE Engineering in Medicine and Biology Society, EMBS (2019). doi:10.1109/EMBC.2019.8857702.

45. Akgun, O. C., Nanbakhsh, K., Giagka, V. & Serdijn, W. A. A Chip Integrity Monitor for Evaluating Moisture/Ion Ingress in mm-Sized Single-Chip Implants. IEEE Trans Biomed Circuits Syst 14, 658–670 (2020).

46. Lee, W. J. et al. Hygroscopic fluorine-doped silicon oxide thin-film-based large-area three-layered moisture barrier using a roll-to-roll microwave plasma-enhanced chemical vapor deposition system. Plasma Processes and Polymers 19, (2022).

47. Rubeck, S. et al. Effect of accelerated hydrothermal aging on the durability of Si-based dielectric thin films. Microelectron Eng 264, (2022).

48. Piras, F. M., Di Mundo, R., Fracassi, F. & Magnani, A. Silicon nitride and oxynitride films deposited from organosilicon plasmas: ToF-SIMS characterization with multivariate analysis. Surf Coat Technol 202, 1606–1614 (2008).

49. Visweswaran, B. et al. Diffusion of water into permeation barrier layers. Journal of Vacuum Science & Technology A: Vacuum, Surfaces, and Films 33, (2015).

50. Lee, H. I. et al. Degradation by water vapor of hydrogenated amorphous silicon oxynitride films grown at low temperature. Sci Rep 7, (2017).

51. Yang, H. L. et al. Silicon oxynitride thin films by plasma-enhanced atomic layer deposition using a hydrogen-free metal-organic silicon precursor and N2 plasma. Mater Sci Semicond Process 164, (2023).

52. Klerk, L. A. et al. TOF-secondary ion mass spectrometry imaging of polymeric scaffolds with surrounding tissue after in vivo implantation. Anal Chem 82, 4337–4343 (2010).

53. Donaldson, N., Baviskar, P., Cunningham, J. & Wilson, D. The permeability of silicone rubber to metal compounds: Relevance to implanted devices. J Biomed Mater Res A 100 A, (2012).

54. 02_Donaldson_1995_aspects_silicone_rubber_mixed_oxides.

55. Donaldson, P. E. K. The Essential Role Played by Adhesion in the Technology of Neurological Prostheses. Int. J. Adhesion and Adhesives vol. 16 (1996).

56. 1996 - Corrosion-induced degradation of microelectronic devices.

57. Hassler, C., Boretius, T. & Stieglitz, T. Polymers for neural implants. Journal of Polymer Science, Part B: Polymer Physics vol. 49 18–33 Preprint at 10.1002/polb.22169 (2011).

58. Caldwell, R. et al. Characterization of Parylene-C degradation mechanisms: In vitro reactive accelerated aging model compared to multiyear in vivo implantation. Biomaterials 232, (2020).

59. Guo, X. et al. The effect of water uptake on the mechanical properties of low-k organosilicate glass. J Appl Phys 114, (2013).

60. Li, Y. et al. Influence of absorbed water components on SiOCH low-k reliability. J Appl Phys 104, (2008).

61. Voskerician, G. et al. Biocompatibility and biofouling of MEMS drug delivery devices. Biomaterials 24, 1959–1967 (2003).

62. Pezzotti, G. et al. Off-stoichiometric reactions at the cell-substrate biomolecular interface of biomaterials: In situ and ex situ monitoring of cell proliferation, differentiation, and bone tissue formation. Int J Mol Sci 20, (2019).

63. Eliaz, N. Corrosion of metallic biomaterials: A review. Materials vol. 12 Preprint at 10.3390/ma12030407 (2019).

64. Gupta, S., Navaraj, W. T., Lorenzelli, L. & Dahiya, R. Ultra-thin chips for high-performance flexible electronics. npj Flexible Electronics vol. 2 Preprint at 10.1038/s41528-018-0021-5 (2018).

65. Stieglitz, T. et al. Highly conformable chip-in-foil implants for neural applications. Microsyst Nanoeng 9, (2023).

66. Donaldson, N., Lamont, C., Idil, A. S., Mentink, M. & Perkins, T. Apparatus to investigate the insulation impedance and accelerated life-testing of neural interfaces. J Neural Eng 15, (2018).

67. Donaldson, N. et. al., PDMS and Silicon-IC passivation for the protection of implantable electronics: part-1, accelerated life tests, under preparation.

