## Supporting Information for "Longevity of Implantable Silicon-ICs for Emerging Neural Applications: Evaluation of Bare Die and PDMS-Coated ICs After Accelerated Aging and Implantation Studies"

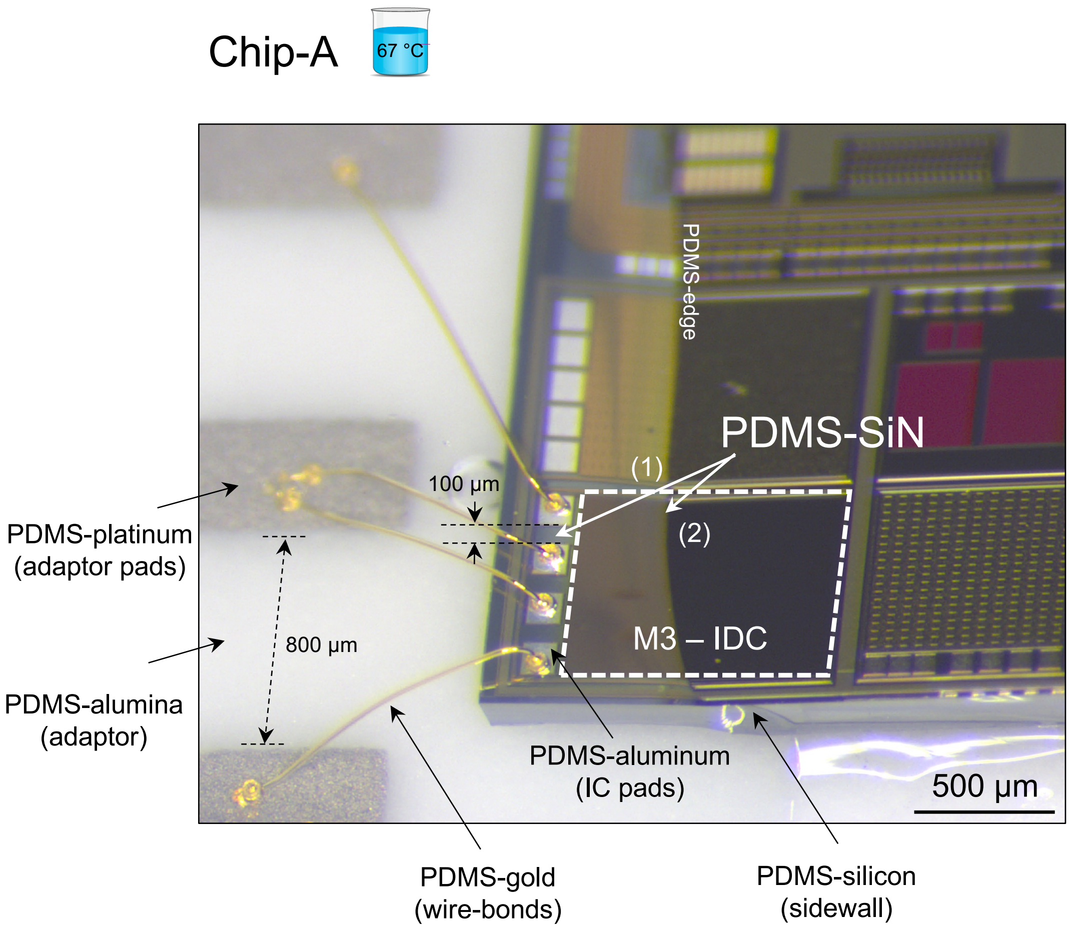


**Figure S1**. A tilted optical micrograph of a representative wire-bonded M3-IDC test structure (Chip-A) used for the accelerated *in vitro* aging study demonstrating the 6 critical PDMS interface bonds on the test structure.

De-bonding of PDMS could affect the electrical measurements during the *in vitro* accelerated aging study. Figure S1 presents the 6 PDMS interface bonds on the test structures. The most critical interface is the PDMS-SiN interface bond which is composed of two regions: (1) the region between the IC pads and (2) the region extending to the PDMS-edge. Both interface regions are crucial for maintaining a stable electrical performance. The interface at the PDMS-edge is directly exposed to phosphate buffered saline (PBS) solution where interfacial debonding between the PDMS and SiN will allow lateral ingress of ionic liquid. All other interfaces are protected from the ionic fluid by PDMS. Nevertheless, due to the moisture permeability of PDMS, these interfaces will be subjected to moisture (gaseous water). At the regions between the IC pads, debonding of the PDMS-SiN interface would allow the condensation of the diffused moisture, resulting in shunt water-leakage paths between the pads [1, 2]. Given the narrow ~100 µm gaps between the pads, PDMS de-bonding in this region will impact the electrical performance.

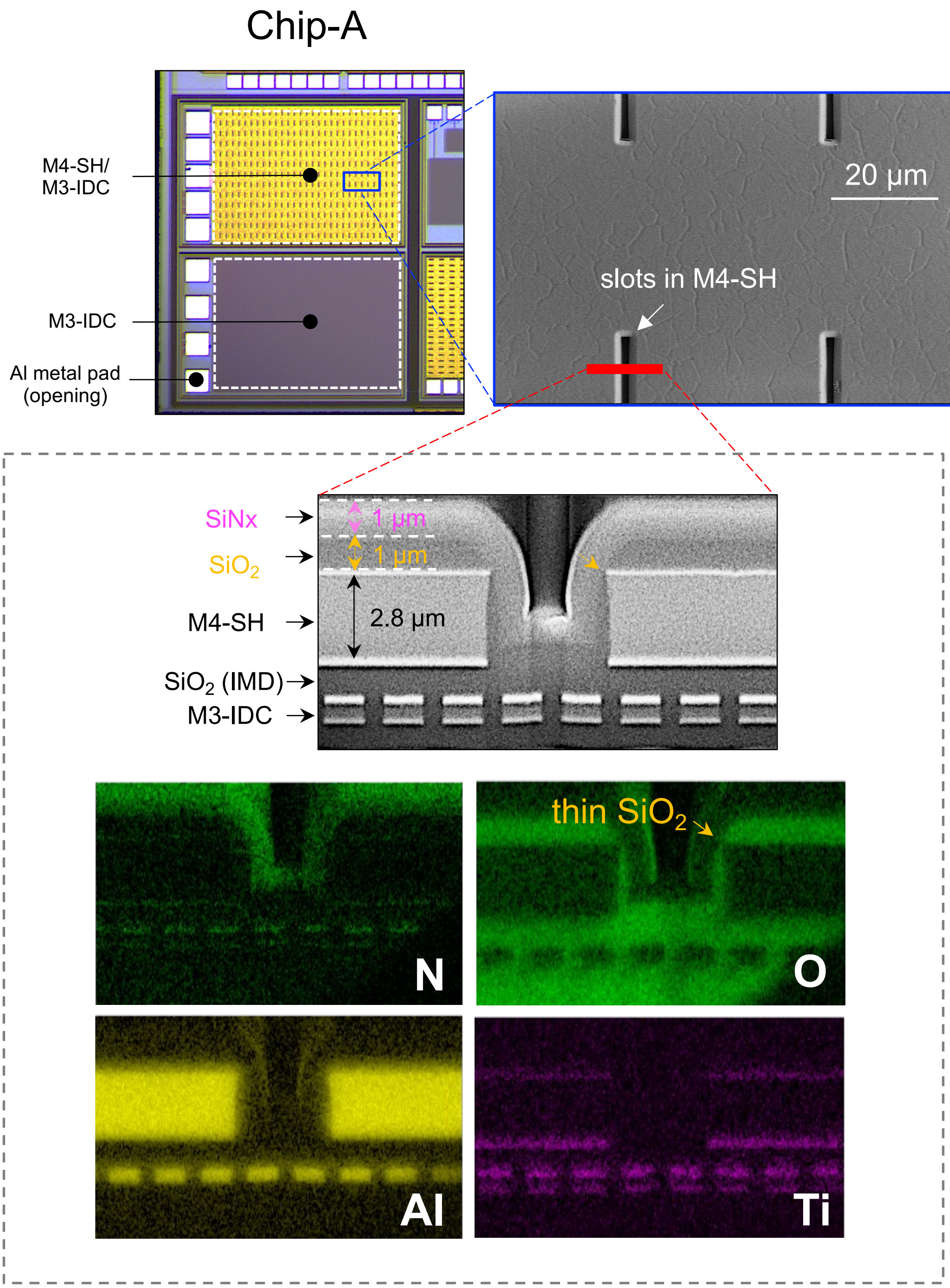


**Figure S2**. Optical and scanning electron microscopy (SEM) micrographs of the M4-SH/M3-IDC (shield layer in Metal-4 and IDC in Metal-3) test structure (Chip-A). Top right SEM depicts the surface microtopography created due to the slots in metal 4 (top metal). Slots are created to in the metal layer to obey metal density rules defined by the IC foundry. Red line indicates the focused ion beam (FIB) cut for creating the cross section. SEM micrograph and energy dispersive X-ray (EDX) elemental mapping of the cross section showing the top material stack of the IC: SiNx/$SiO_2$ passivation layers, Metal-4 shield layer and the M3-IDC structure. The presence of the top metal creates  poor conformality in the passivation layers, resulting in a much thinner $SiO_2$ passivation layer (<100 nm) on the edges of Metal-4.

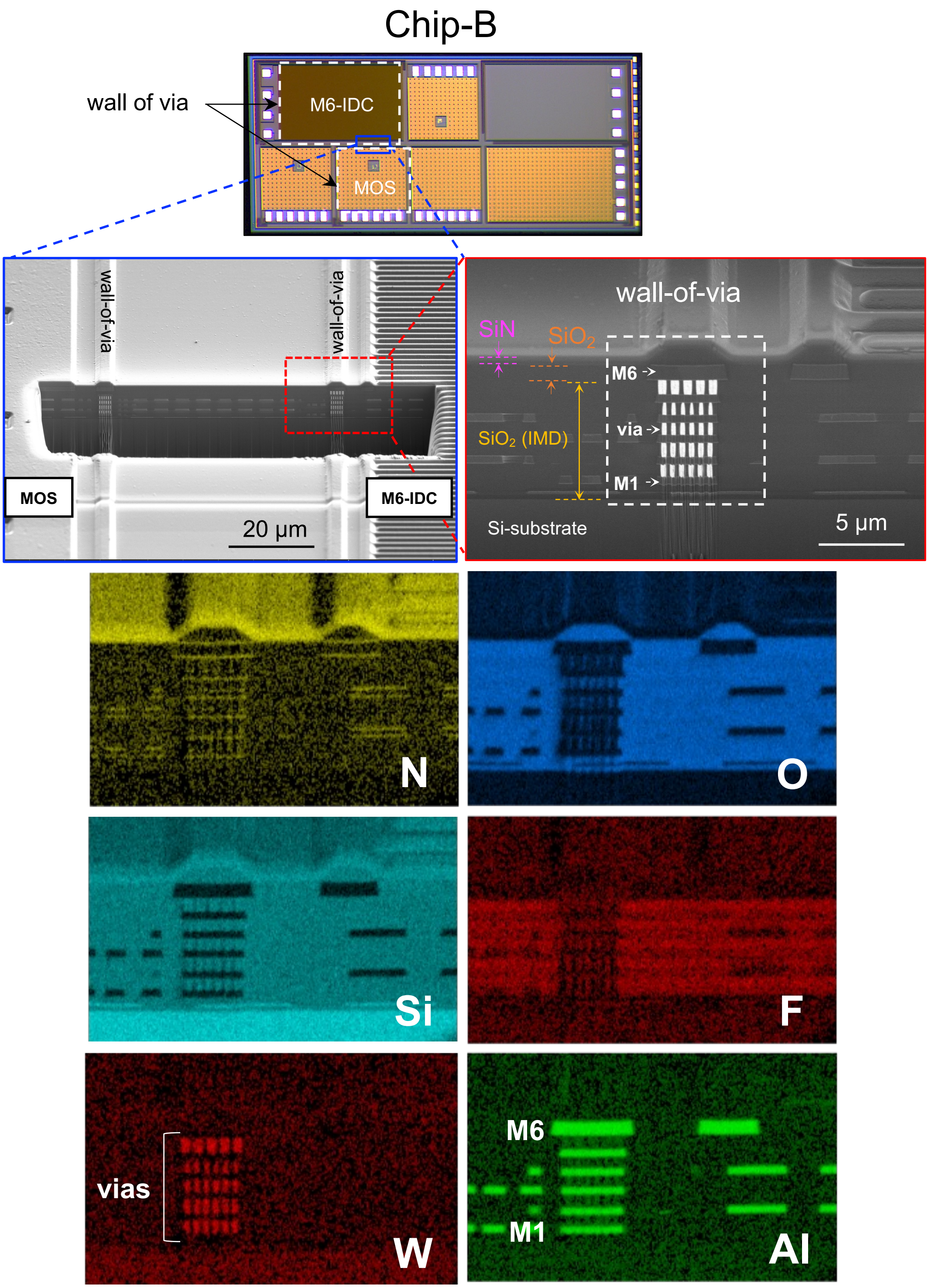


**Figure S3**. Optical and electron micrographs with EDX elemental mapping of wall of via (WoV) on a Chip-B IC. White dashed lines in optical micrograph show the wall-of-via (WoV) structure implemented around each test structure. Surface and cross-sectional SEM images show the WoV around the MOS and M6-IDC structures. Surface topography is seen as a result of using top metallization (M6). EDX elemental mapping shows the material stack of Chip-B. The WoV is implemented using M1 to M6 aluminum (Al) metallization and Tungsten (W) vias. EDX elemental mapping showing the materials used in the IC stack. The aluminum metallization is placed in between thin titanium nitride (TiN) layers, which are used as metal diffusion barriers. The fluorine (F) intensity in the intermetallic dielectric (IMD) layers shows that the IMD layers for Chip-B are Fluorine-doped silicon oxide (SiOF) to reduce the dielectric constant of the layers (*k* ~ 3.5).

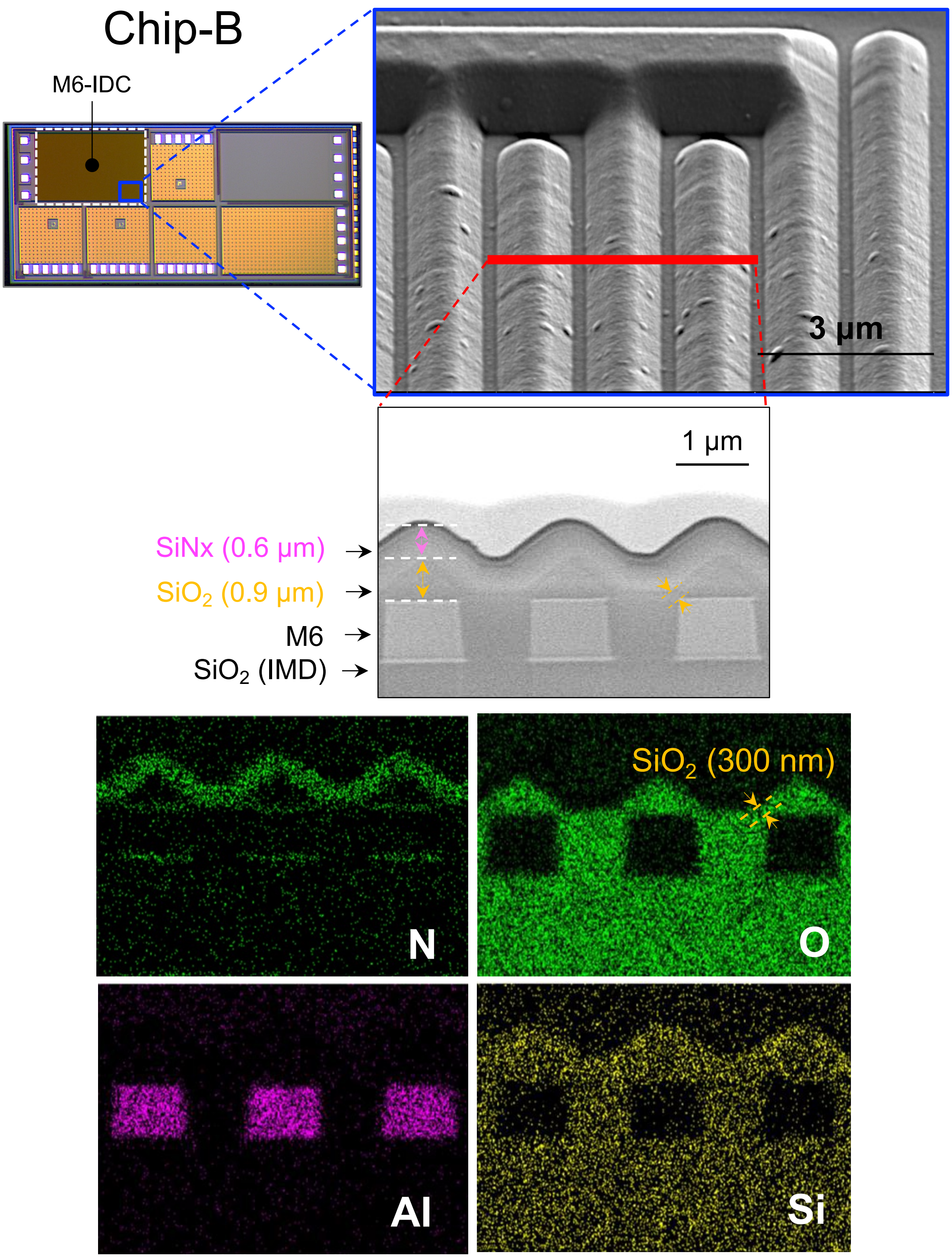


**Figure S4**. Optical and electron micrographs of a M6-IDC test structure on Chip-B showing surface microtopography. Tilted SEM images show surface microtopography created due to the use of the top metal layer (M6). Red line indicates the FIB cut for cross sectioning. SEM and EDX elemental mapping of the cross section depicts the top few layers showing SiNx and $SiO_2$ passivation. Due to the presence of the top metallization, poor conformality of the oxide passivation can be observed resulting in a thinner (~300 nm) $SiO_2$ passivation on the edges of the metal fingers.

**Table S1**: Test structures on Chip-A with specifications.

| Structure | Image | Design parameters |
|---|---|---|
| M3-IDC<br>(1.15 x 0.85 mm) | 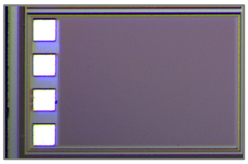 | All IDC metal in Metal 3<br>Pads in Metal 4<br>Bond pads size: 150x150 μm<br>Passivated everywhere except pads<br>Wall of via (WoV) around the whole structure<br>Finger width W = 1 μm<br>Finger gap G = 0.6 μm<br>Number of fingers N = 716 (358 each electrode)<br>Finger length = 844.4 μm |
| M4-SH/M3-IDC<br>(0.97 x 0.68 mm) | 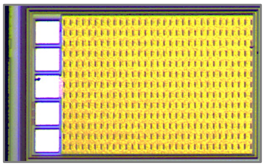 | All IDC metal in Metal 3<br>Pads and shield in Metal 4<br>Bond pads size: 150x150 μm<br>Passivated everywhere except pads<br>WoV around the whole structure<br>Finger width W = 1 μm<br>Finger spacing G = 0.6 μm<br>Number of fingers N = 606 (303 each electrode)<br>Finger length = 664.4 μm |

**Table S2**: Test structures on Chip-B with specifications.

| Structure | Circuit layout | Specifications |
|---|---|---|
| **M5-IDC**<br>(1.1 mm x 0.7 mm) | 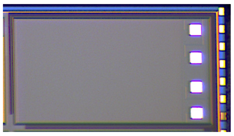 | All IDC metal in Metal 5<br>Pads in Metal 6<br>Bond pads size: 80 x 80 µm<br>Passivated everywhere except pads<br>Wall of via (WoV) around the whole structure<br>Finger width W = 1 µm<br>Finger gap G = 0.6 µm<br>Number of fingers N = 716 (358 each electrode)<br>Finger length = 700 µm |
| **M6-SH/M5-IDC**<br>(0.9 mm x 0.6 mm) | 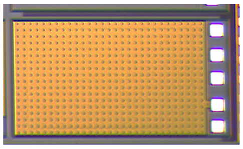 | All IDC metal in Metal 5<br>Pads and shield (SH) in Metal 6<br>Bond pads size: 80 x 80 µm<br>Passivated everywhere except pads<br>Wall of via (WoV) around the whole structure<br>Finger width W = 1 µm<br>Finger gap G = 0.6 µm<br>Number of fingers N = 606 (303 each electrode)<br>Finger length = 609 µm |
| **M6-IDC**<br>(1.1 mm x 0.7 mm) | 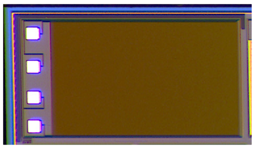 | All IDC metal in Metal 6<br>Pads in Metal 6<br>Bond pads size: 80 x 80 µm<br>Passivated everywhere except pads<br>Wall of via (WoV) around the whole structure<br>Finger width W = 1 µm<br>Finger gap G = 1 µm<br>Number of fingers N = 716 (358 each electrode)<br>Finger length = 704 µm |
| **NMOS** | **Shielded**<br>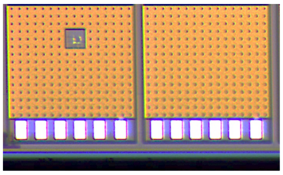 | NMOS transistors, with and without shield.<br>Pads in Metal 6.<br>Shield in Metal 5 and 6 (double shielded)<br>Passivated everywhere except pads<br>NMOS total W=400 µm (W=20 µm with N=20 fingers)<br>NMOS L = 0.36 µm<br>Electro static protection used for all pads |

**Table S3**: Overview of test structures used in different aging environments and the material analysis tools used for each structure.

| Chip | Test structure | Aging | DC bias | Aging duration (month) | Electrical[a] | SEM[b]/ AFM | ToF-SIMS/ XPS[c] |
|---|---|---|---|---|---|---|---|
| A (n=30) | M3-IDC | PBS @ 67 ºC | Unbiased (n=4) | 3, 6, 10, 16 | EIS | n=4 | n=1 |
| | | | 5 V (n=4) | 6, 12 | EIS | n=2 | n=2 |
| | | | 15 V (n=6) | 6, 12, 16 | EIS | n=3 | n=3 |
| | | DI @ 67 ºC | 15 V (n=2) | 16 | EIS | - | n=1 |
| | | rat[d] | Unbiased (n=6) | 3, 7, 12 | - | n=6 | n=3 |
| | M4-SH/ M3-IDC | PBS @ 67 ºC | 5 V (n=8) | 10, 12, 16 | EIS | n=2 | - |
| B (n=54) | M5-IDC | PBS @ 67 ºC | Unbiased (n=4) | 6, 12, 16 | EIS | n=2 | n=1 |
| | | | 5 V (n=4) | 6, 12 | EIS | n=2 | n=2 |
| | | | 15 V (n=6) | 12, 16 | EIS | n=2 | n=2 |
| | | DI @ 67 ºC | 15 V (n=2) | 16 | EIS | - | n=1 |
| | | rat[d] | Unbiased (n=6) | 3, 7, 12 | - | n=6 | n=3 |
| | M6-IDC | PBS @ 67 ºC | Unbiased (n=4) | 16 | EIS | n=1 | - |
| | | | 5 V (n=6) | 12 | EIS | n=2 | - |
| | | | 15 V (n=6) | 12, 16 | EIS | n=2 | - |
| | M6-SH/ M5-IDC | PBS @ 67 ºC | 15 V (n=4) | 12 | EIS | n=2 | - |
| | MOS | PBS @ 67 ºC | Unbiased (n=4) | 12 | $V_{GS}$ - $I_{DS}$ | n=2 | - |
| | M6/M5-SH/ MOS | PBS @ 67 ºC | Unbiased (n=1) | 12 | $V_{GS}$ - $I_{DS}$ | - | - |
| | Dielectric | PBS @ 67 ºC | Unbiased (n=1) | 5 | array measurements | - | - |

Optical microscopy (up to 200x magnification) was done on all devices.

[a]All electrical measurements were done monthly until the end of the *in vitro* accelerated aging.
[b]For samples which were analyzed using both SEM and ToF-SIMS/XPS, the SEM analyses was performed after the ToF-SIMS/XPS analysis.
[c]ICs were first analyzed using ToF-SIMS, both in positive and negative modes. For quantification, Ics were further analyzed using XPS. During the ToF-SIMS analysis of aged ICs, reference ICs (pristine, as is from foundry) was used for comparison.
[d]In vivo samples were first microscopically inspected, decapsulated and again inspected using microscopy. AFM at the PDMS-edge boundary was used to determine the thickness of the SiNx/SiO2 passivation. Later, samples were analyzed using ToF-SIMS/XPS. Finally, the ICs were inspected with SEM.

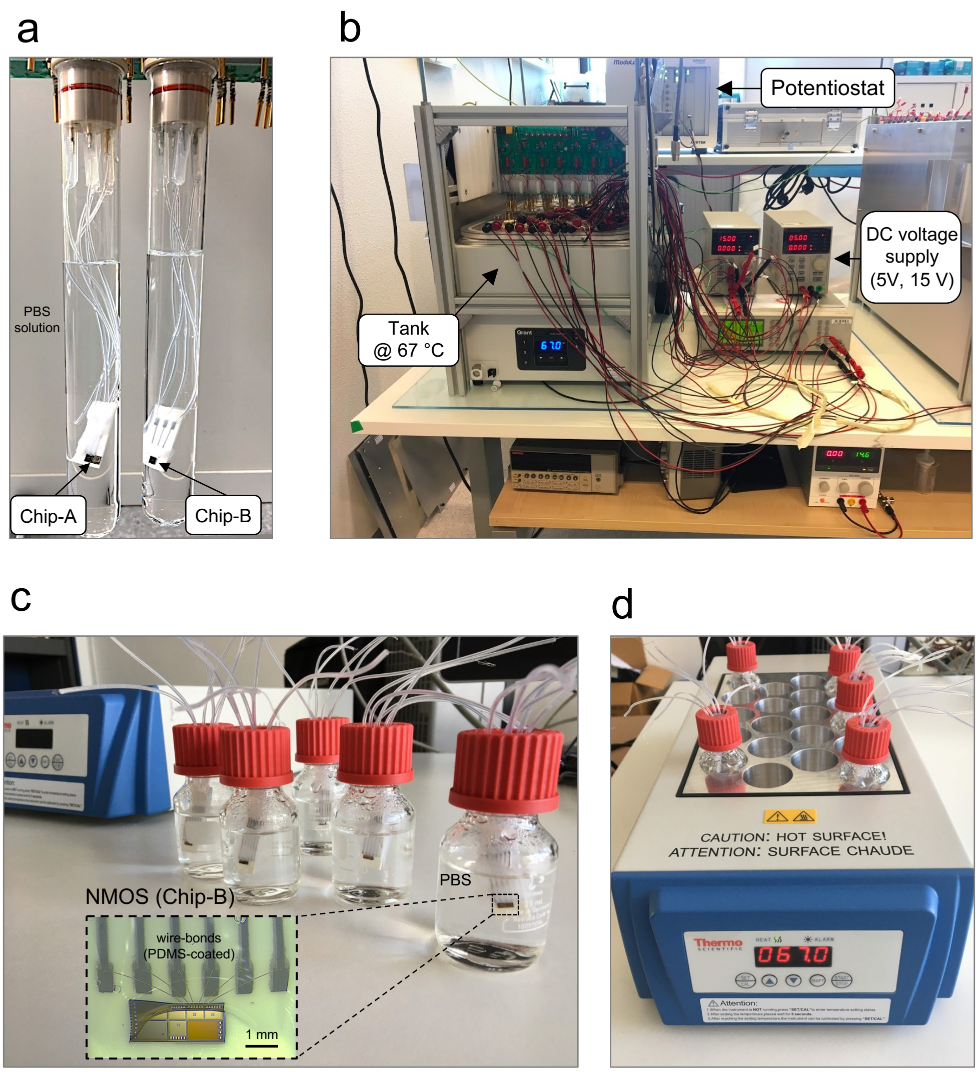


**Figure S5**. **a)** Chip-A and B ICs connected to 3-contact ceramic (alumina) adaptors immersed in vials filled with 50 ml of phosphate buffered saline (PBS) solution for accelerated *in vitro* aging of interdigitated capacitor (IDC) structures. **b)** Heat-regulated water-bath tank and voltage supplies (5 V and 15 V DC) used for accelerated aging and electrical stressing of IDC test structures. Potentiostat used for electrochemical impedance spectroscopy (EIS). **c)** N-channel metal oxide semiconductor (NMOS) transistors (Chip-B) wire-bonded to 6-contact ceramic substrates, partially PDMS-coated and immersed in PBS solution for *in vitro* accelerated aging, **d)** heater used for maintain a constant temperature at 67 °C.

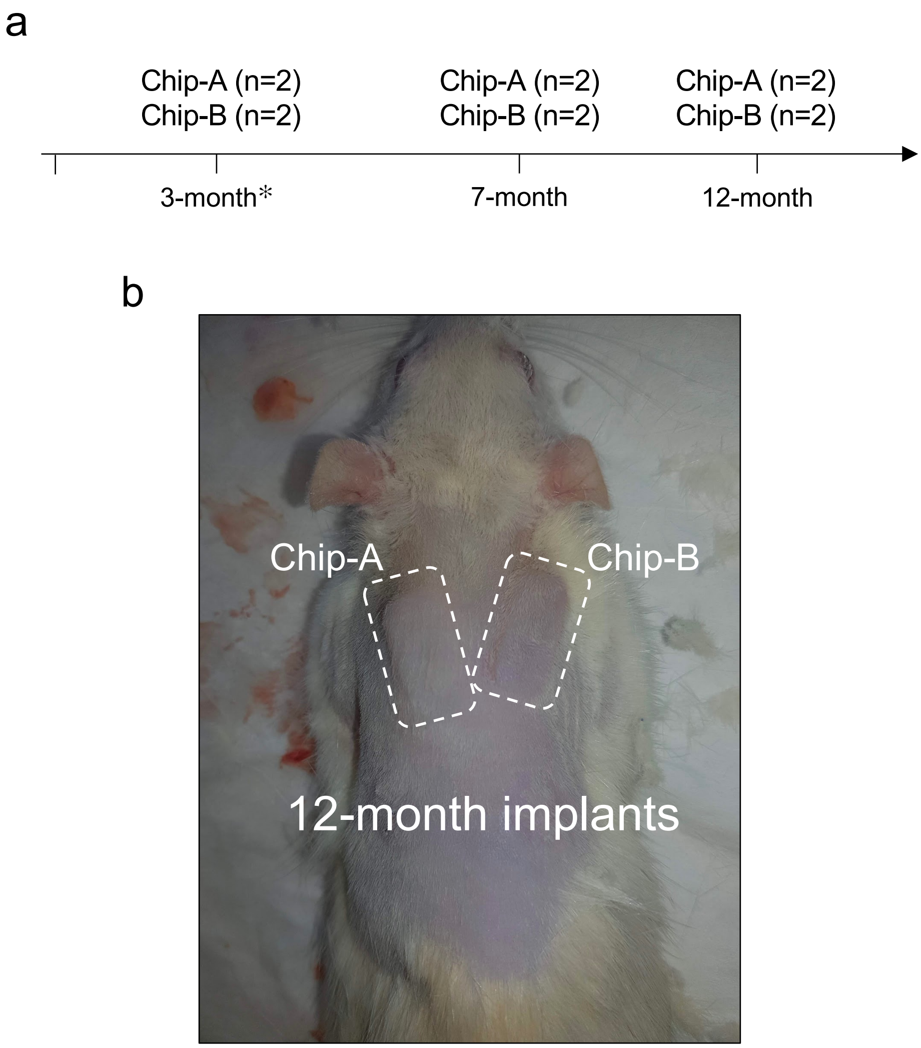


**Figure S6**. **a)** Explantations for the implanted Chip-A and B ICs in rats. **b)** image of a rat animal model after 12 months of implantation with complete wound healing and no observable inflammation around the two implanted chips. (* One animal died during the study due and was incinerated with the device still implanted.)

### Chemical composition of IC passivation layers (as received from foundry)

**Table S4**: Chemical composition of the SiNx and $SiO_2$ passivation layers for Chip-A and B ICs (as is from foundry), determined using XPS surface and depth profiling analysis. The measured carbon (C) is from surface contamination and is generally found at the surface of samples exposed to ambient air. After 1 sputter cycle the carbon contamination is removed.

| Chip A | Measured Depth (nm) | Si (at%) | N (at%) | O (at%) | C (at%) |
|---|---|---|---|---|---|
| SiNx | 0[a] | 29.9 | 7.9 | 39.8 | 21.8[b] |
| | 5 | 49.7 | 45.5 | 4.8 | 0 |
| | 10 | 51.5 | 48.5 | 0 | 0 |
| | 14 | 50.6 | 49.4 | 0 | 0 |
| $SiO_2$ | 1050 | 33 | 0 | 67 | 0 |
| **Chip B** | **Measured Depth (nm)** | **Si (at%)** | **N (at%)** | **O (at%)** | **C (at%)** |
| SiNx | 0[a] | 26.7 | 10.8 | 36.1 | 26.3 |
| | 5 | 49.6 | 47 | 3.5 | 0.1 |
| | 10 | 49.4 | 49.3 | 1.3 | 0 |
| | 14 | 49.0 | 51 | 0 | 0 |
| $SiO_2$ | 1100 | 32.7 | 0 | 67.3 | 0 |

[a] For XPS surface measurement, the information depth is approximately 7 nm.
[b] The carbon detected on the surface is from ambient environment (adventitious carbon).

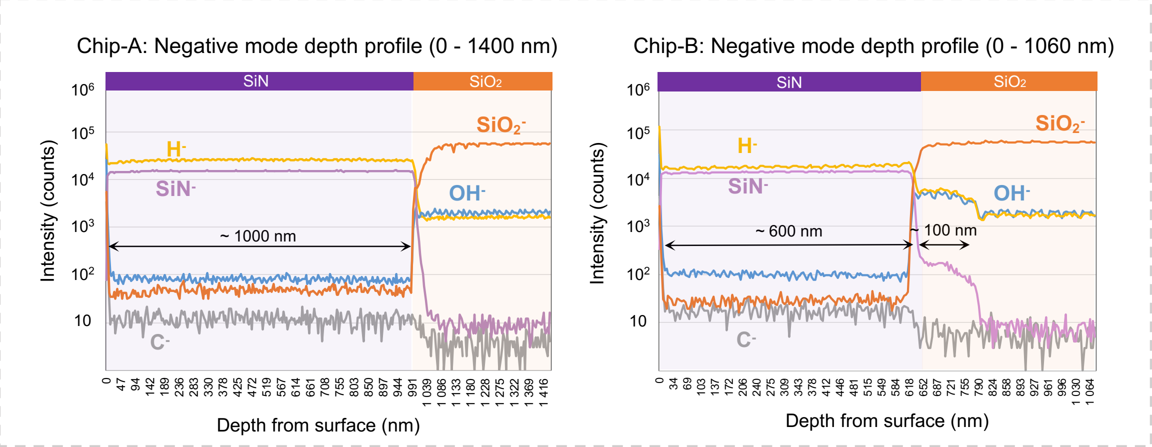


**Figure S7.** Negative mode ToF-SIMS depth profiles of Chip-A (right) and Chip-B (left) ICs (as is from foundry).

The quantification of the hydrogen content (H) in the SiNx passivation was done in ToF-SIMS software using a relative sensitivity factor (RSF). The RSF was derived from measurement results performed on known SiN:H reference layers. The hydrogen content on these SiN:H reference layers has been previously calibrated using elastic recoil detection (ERD).

### Interdigitated capacitor (IDC)

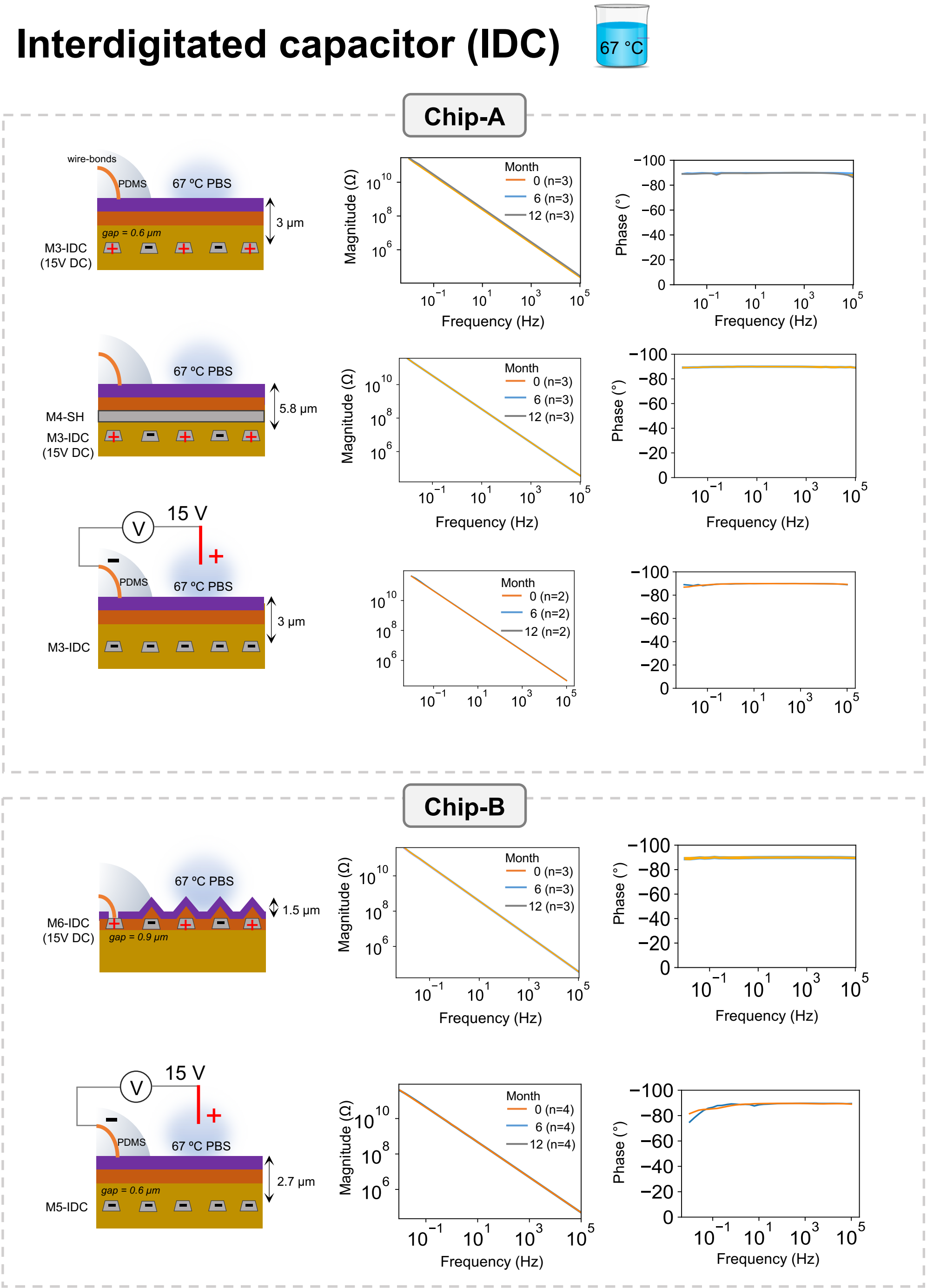


**Figure S8**. Electrochemical impedance spectroscopy (EIS) results of interdigitated capacitor (IDC) test structures on Chip-A and Chip-B ICs over 12 month accelerated *in vitro* aging in PBS solution at 67 °C.

### Electrical failures 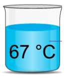


**Table S5**: Overview of test structures and the failures observed during the in vitro accelerated aging.

| Chip | Test structure | Aging | DC bias | Duration in test (months) | Time to failure | Failure |
|---|---|---|---|---|---|---|
| A | M3-IDC | PBS @ 67 °C | unbiased | 6, 10, 12, 16 (n=4) | 7-month (n=1) | Wire-bond corrosion |
| | | | 5 V | 12 (n=4) | | |
| | | | 15 V | 12, 16 (n=6) | 3-month (n=1) | Wire-bond corrosion |
| | | DI @ 67 °C | 15 V | 16 (n=2) | 5-month (n=1) | Wire-bond corrosion |
| | | rat | Unbiased | 3,7,12 (n=6) | - | - |
| | M4-SH/ M3-IDC | PBS @ 67 °C | 5 V | 10, 12 (n=8) | 7-month (n=3)<br>5-month (n=1) | Passivation<br>Wire-bond corrosion |
| B | M5-IDC | PBS @ 67 °C | unbiased | 6, 12, 16 (n=4) | | |
| | | | 5 V | 12 (n=4) | | |
| | | | 15 V | 12, 16 (n=6) | | |
| | | DI @ 67 °C | 15 V | 16 (n=2) | | |
| | | rat | unbiased | 3,7,12 (n=6) | - | - |
| | M6-IDC | PBS @ 67 °C | unbiased | 16 (n=4) | 3-month (n=2) | Passivation crack |
| | | | 5 V | 12 (n=6) | 1-month (n=1)<br>3-month (n=1) | Passivation crack |
| | | | 15 V | 12, 16 (n=6) | 1-month (n=2) | Passivation crack |
| | M6-SH/ M5-IDC | PBS @ 67 °C | 15 V | 12 (n=4) | 10-month (n=1) | Wire-bond corrosion |
| | MOS | PBS @ 67 °C | unbiased | 12 (n=4) | - | - |
| | M6-M5-SH/ MOS | PBS @ 67 °C | unbiased | 12 (n=1) | - | - |
| | Dielectric | PBS @ 67 °C | unbiased | 5 (n=1) | 5-month (n=1) | Wire-bond corrosion |

During the *in vitro* accelerated aging study, a group of samples showed irregularities in the electrical results. Table S5 gives an overview of these test structures. These samples were taken out from the remainder of the accelerated study and were analyzed using optical and electron microscopy. Results showed failures to be either due to wire-bond corrosion, stress-induce cracks in the passivation or poor conformality of the passivation layer.

### Wire-bond corrosion

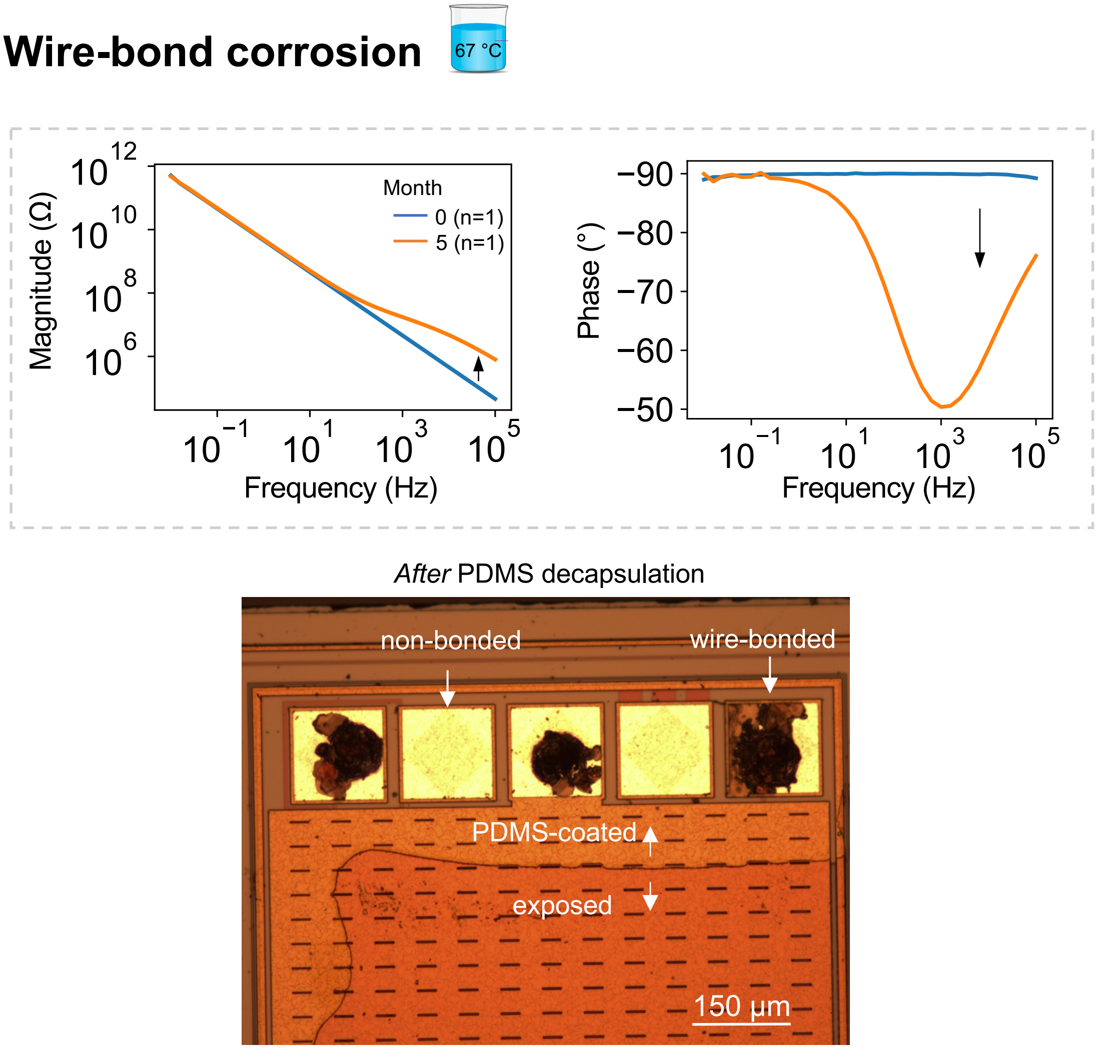


**Figure S9**. EIS results and optical micrographs of sample with wire-bond corrosion in the PDMS-coated region after 4-months of soaking in PBS solution at 67°C. EIS results presented as Bode plots for a M4-SH/M3-IDC test structure (Chip-A) at 5-month show change in higher frequency ranges. Optical micrograph of the structure after PDMS decapsulation show aluminum corrosion on the wire-bonded pads. No corrosion is seen for the non-bonded pads.

In this study, a few number of IDC samples (n=5 out of in total n=56 ) showed change in EIS results in the frequency range of 10 Hz to 100 KHz. Optical microscopy revealed that these samples experienced wire-bond corrosion which resulted in a high ohmic wire-bond connection to the IDC structure. Closer examination revealed the corrosion to mainly only occur on the wire-bonded pads (Figure S9). For PDMS-coated wire-bonds, this type of corrosion has been reported before [3] and is most likely due to the galvanic corrosion at the Al-Au interface which is triggered by moisture. The galvanic corrosion results in the degradation of the less noble metal, in this case, aluminum. PDMS-coated aluminum pads without Au wire bonds, on the other hand, did not exhibit any corrosion on all tested samples during the accelerated study in PBS solution at 67 °C.

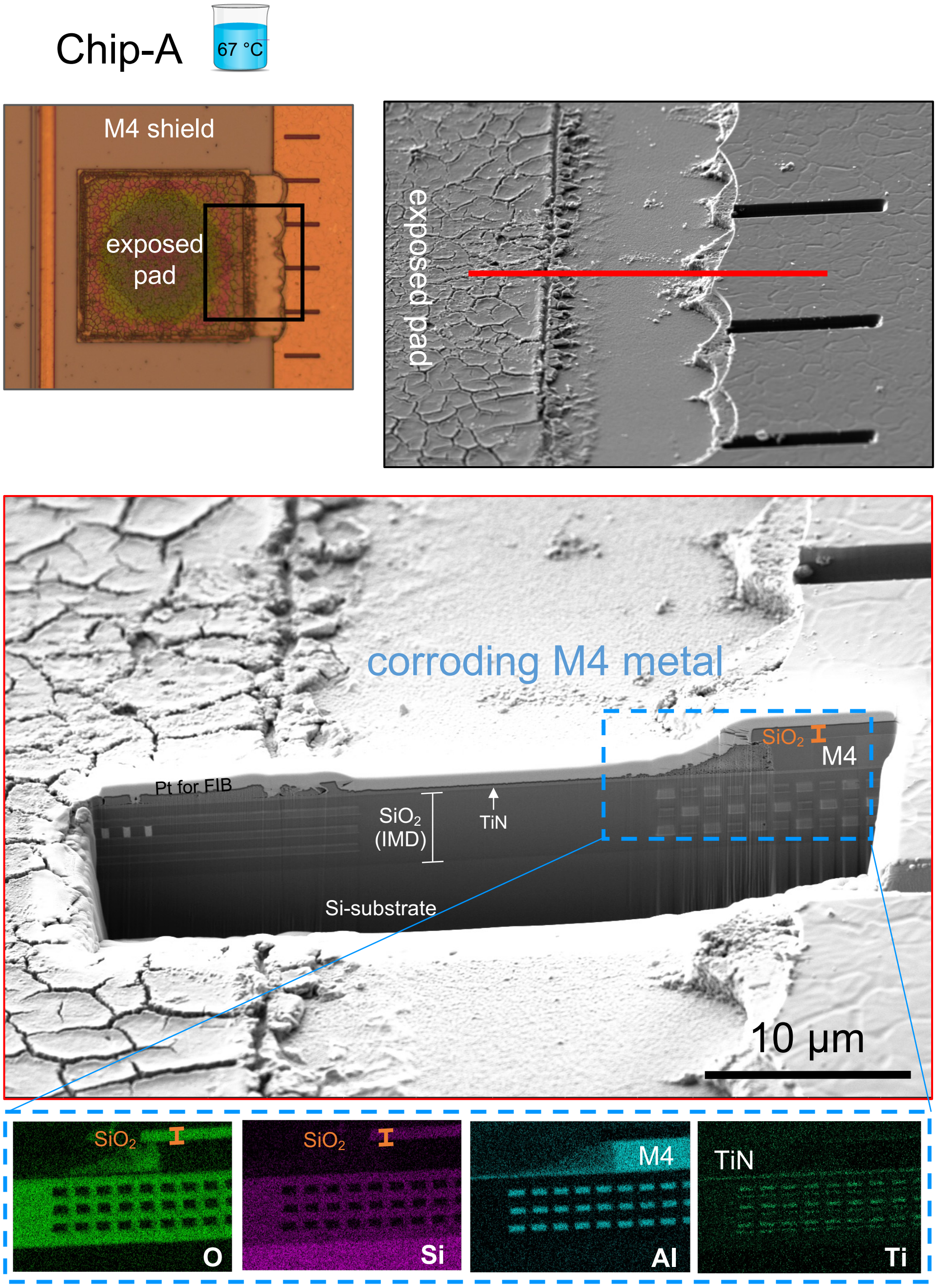


**Figure S10**. Optical and electron micrographs and EDX elemental mapping of an exposed aluminum pad connected to top metal shield (M4-SH) on a Chip-A sample after 10 months immersion in PBS solution at 67°C. Red line indicates the FIB cut for cross section analysis.

Metal pads are openings in the passivation to allow for electrical connections to the chip. Due to the opening, we were concerned if the corrosion of the aluminum pads would allow any liquid ingress, specially from the edges of the pads (the boundary between metal and the passivation). For this purpose, to evaluate the worst-case scenario, during the *in vitro* accelerated aging, some pads were left uncoated and exposed to PBS solution. Figure S10 shows an exposed pad connected to the M4-SH layer. After 10 months severe corrosion. Despite the severe corrosion, cross section SEM imagining and EDX elemental mapping revealed intact buried Al metallization. The thin titanium nitride (TiN) layer, used as a metal diffusion barrier, is also visible with no signs of corrosion, demonstrating its high stability in PBS solution at 67°C. Results indicate no ingress of corrosive liquid into the chip from the pad openings.

### Passivation planarity and stress-induced cracks

In the fabrication process of a silicon IC, each metal layer is covered by an insulating layer. For the deep metal layers, i.e. M1-M3 or M1-M5, for Chip-A and Chip-B, respectively, this insulating layer is the $SiO_2$ interlayer dielectric, and is always followed by a planarization step. When a topmost metal is used, i.e. M4 or M6, for Chip-A and Chip-B, respectively, the IC passivation (PECVD layer of $SiO_2$ followed by SiNx) is directly deposited instead with no final planarization step. In this case, no planarization step is performed, and the microtopography of that topmost metal layer will define the planarity of the IC passivation (Figures S2 and S3). A non-planar passivation layer may experience stress at high-aspect ratio features that can result in cracks [1]. Such stress-induced cracking would greatly compromise the barrier properties of the passivation and allow water/ion ingress points within the IC.

In our study, such stress was shown to be a source of failure in test structures where the topmost metal was included in the IDC design. In fact, Figures S11 and S12 depict the observed passivation cracking and subsequent metal corrosion which was introduced or accelerated while biasing the topmost metal, both for Chip-A and B. All the IDC structures that did not use top metal did not show similar failures, as was demonstrated.

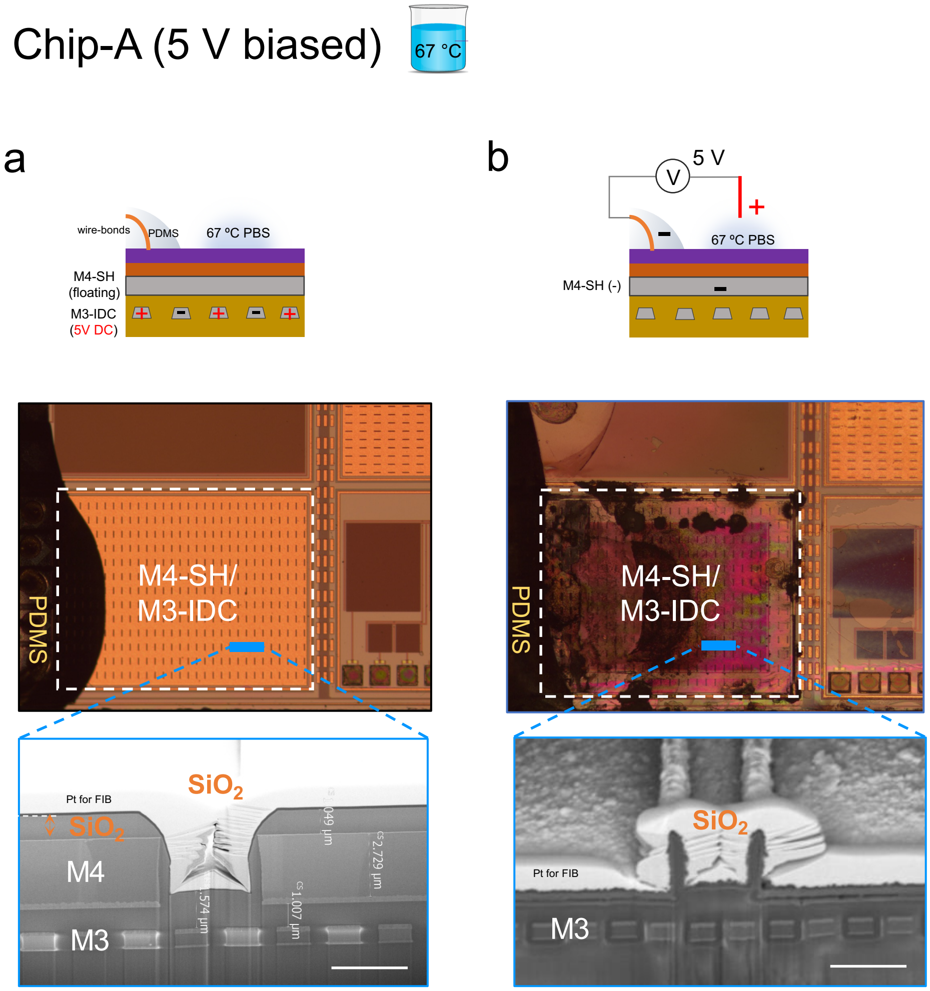


**Figure S11**. Optical micrograph and SEM cross-section images of two M4-SH/M3-IDC test structures (Chip-A) after accelerated *in vitro* testing with biasing. **a**) Optical micrograph of a sample after 10 months of accelerated aging at 67 °C with a continuous 5 V bias applied between the combs of the M3-IDC. M4-SH was not biased and was left floating. Blue line indicates FIB cut for cross section analysis. Cross-sectional SEM image showing total dissolution of the SiNx passivation after 10 months of soaking with no visible signs of corrosion. **b**) Optical micrograph of a different sample with a similar test structure but after 9 months applying the 5 V DC bias between the M4-SH and PBS solution showing severe degradation. Cross-sectional SEM image of the damaged area shows complete loss of both passivation layers (SiNx and $SiO_2$) and the M4-SH (2.8 μm) layer, leaving only the $SiO_2$ side walls. The M3-IDC and other buried material stacks remained intact despite the aggressive electrolysis (scale bar is 3 μm).

Utilizing the top-most metal can results in microtopography on the IC. Such microtopography may lead to poor conformality and stress in the IC passivation layers. Figure S11 shows the M4-SH/M3-IDC test structure on two different Chip-A samples after accelerated *in vitro* testing. On these structures, the presence of slots in the top metal shield layer creates microtopography on the IC.

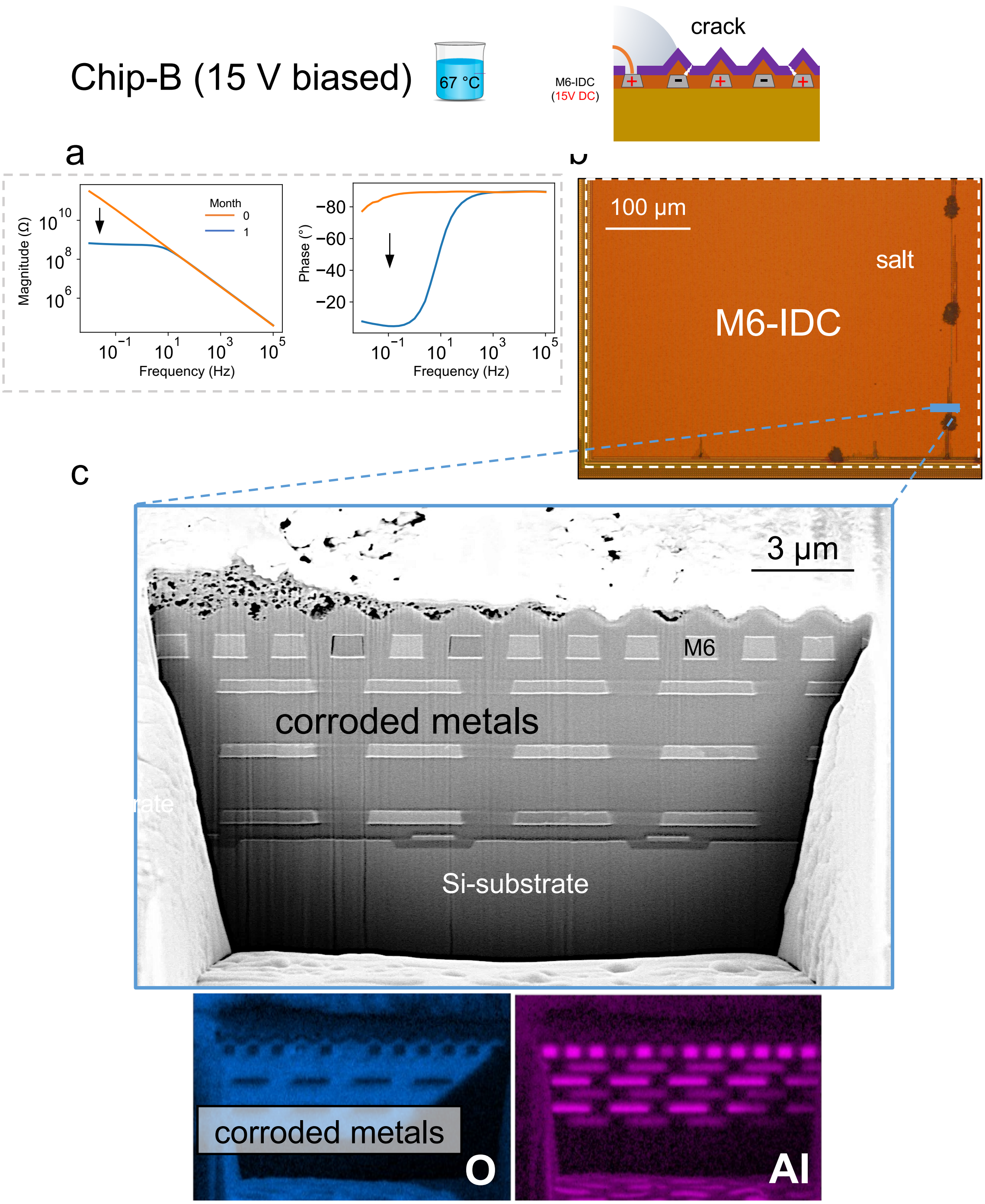


**Figure S12**. **a**) EIS results given as Bode plots for a representative M6-IDC (Chip-B) structure showing EIS irregularities after being applied to a 15 V DC bias voltage for 1-month. At month-1 EIS results show a significant drop in magnitude with more resistive behavior (phase ~ -20) at frequencies below 10 Hz. **b)** Optical micrograph of the representative M6-IDC structure with metal corrosion and salt residue on the corroded sites. Blue line indicates the FIB cut used for cross section analysis. **c)** Cross-sectional SEM image and EDX elemental mapping show the corroded aluminum metal fingers having higher oxygen content.

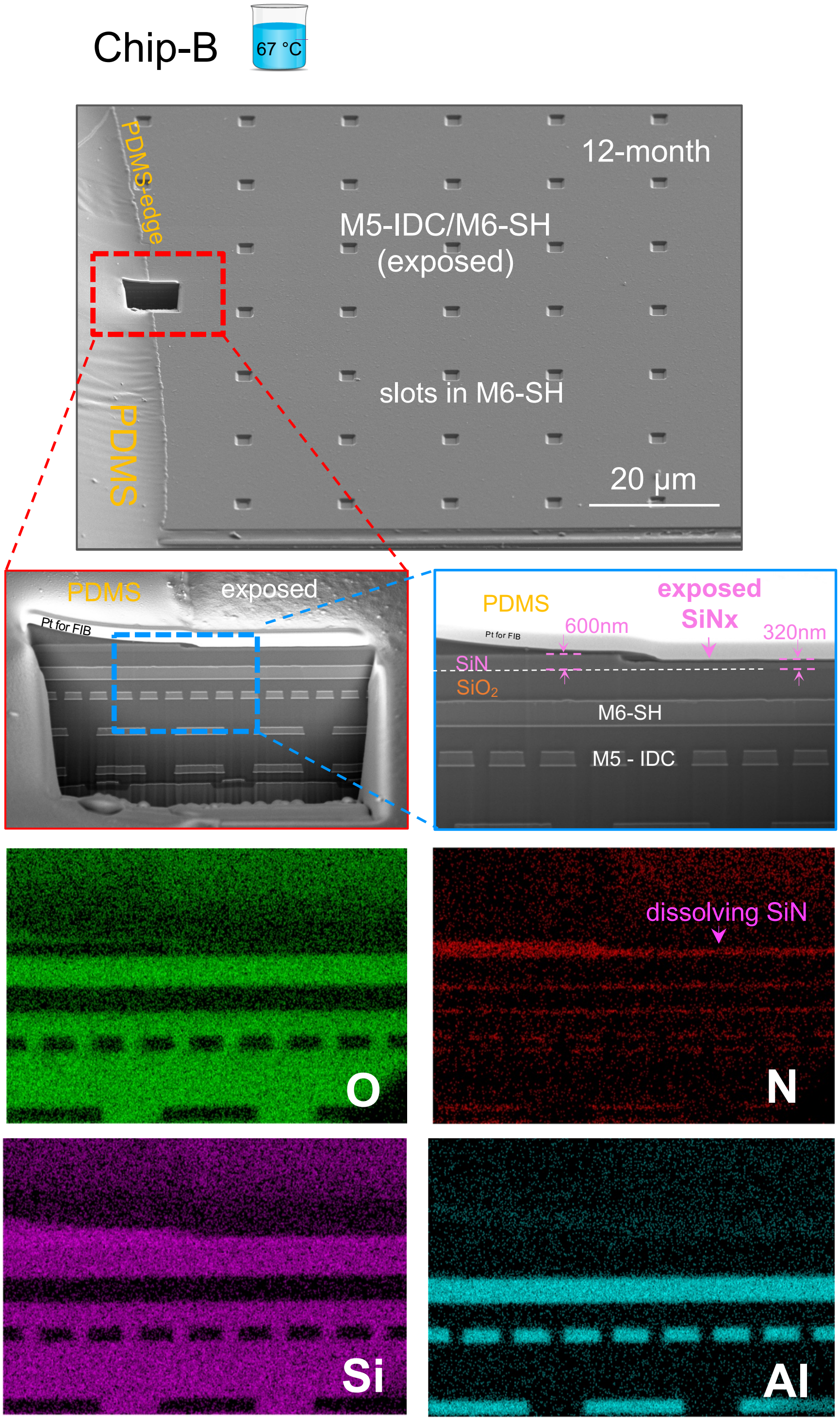


**Figure S13**. Tilted SEM surface image of a M6-SH/M5-IDC structure (Chip-B) after 12 months accelerated aging in PBS solution at 67 °C. Slots in M6 barrier result in microtopography on the IC surface. Slots are created to obey the metal density rules specified by the IC foundry. FIB cut (red square) on the PDMS-edge is used to evaluate the IC stack stability. Magnified SEM images and EDX elemental mapping of cross-section reveal a dissolution of the SiNx passivation in the exposed area, leaving 320 nm of SiNx in the exposed region of the IC. This results in a dissolution rate of ~ 22 nm/month for the SiNx on Chip-B in PBS solution at 67 °C. The PDMS-coated area appears intact (~600 nm).

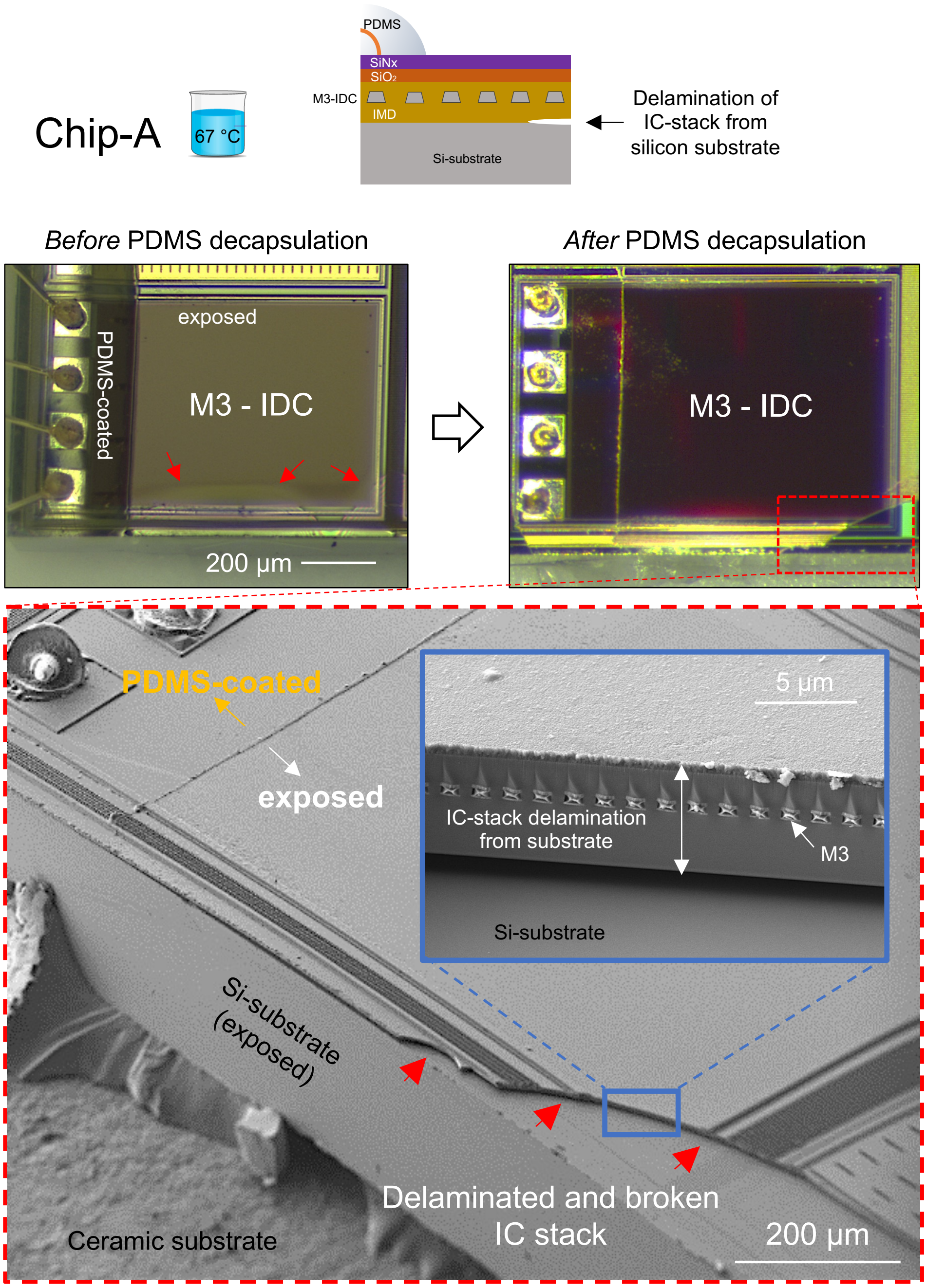


**Figure S14**. Tilted optical and electron micrographs of a M3-IDC test structure (Chip-A) showing delamination of IC-stack from silicon substrate after 16 months aging in PBS solution at 67 °C . In optical micrographs, delamination is seen as color fringes on the IC edge near the sidewall in the uncoated region, shown with red arrows (left). After PDMS decapsulation, a section of the delaminated area broken off due to handling, shown in red dashed square (right). Tilted SEM image of the chip sidewall after PDMS-decapsulation (bottom). Inset: magnified SEM image from the broken area where delamination of the entire IC stack from the Si-substrate is visible. The metallization used for the M3-IDC test structure is also visible.

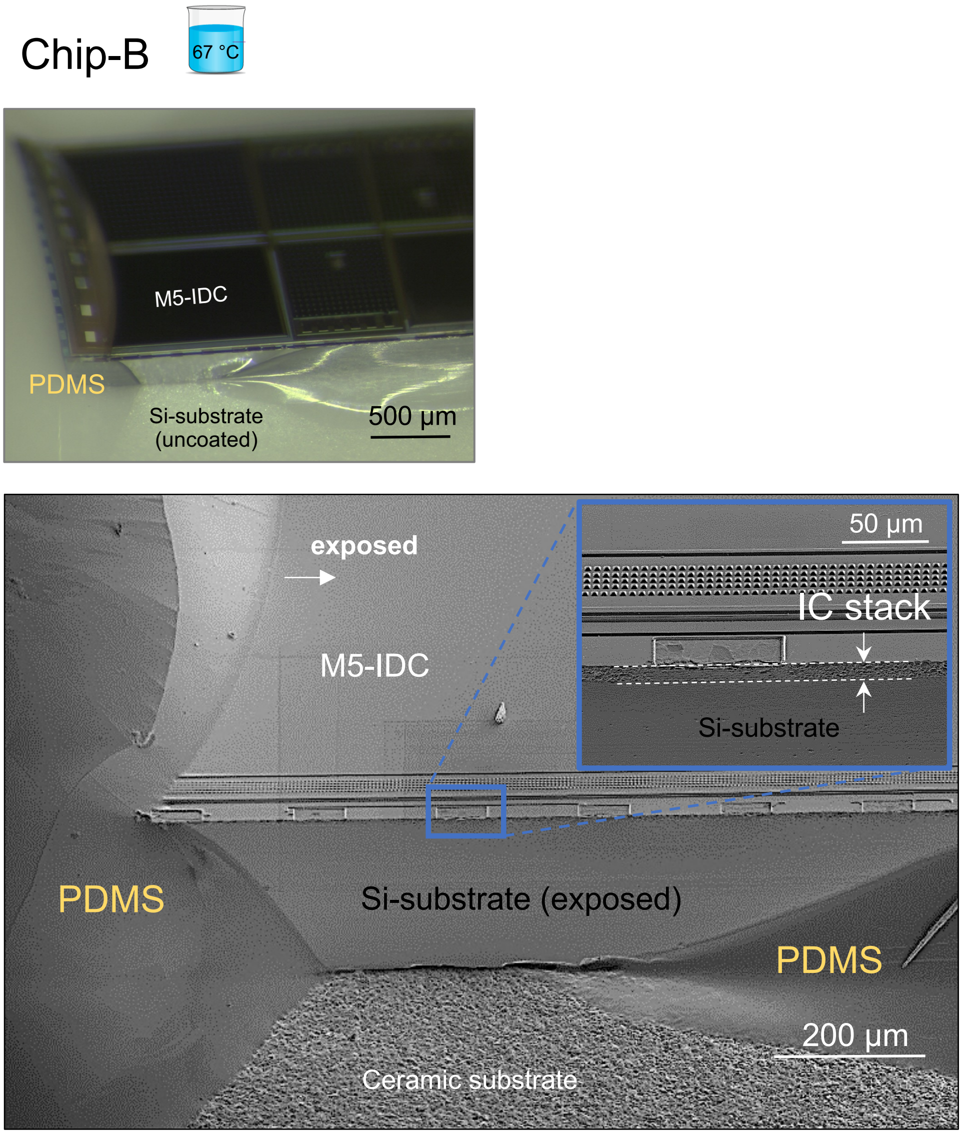


**Figure S15**. Tilted optical and electron micrograph of a M5-IDC test structure (Chip-B) after 16 months of accelerated aging in 67 °C PBS solution showing no delamination on the chip side wall in the uncoated region. Inset: magnified electron image where no delamination is observed between the IC stack (~ 9 µm) and the Si-substrate.

Chip-A (explanted after 3 months)

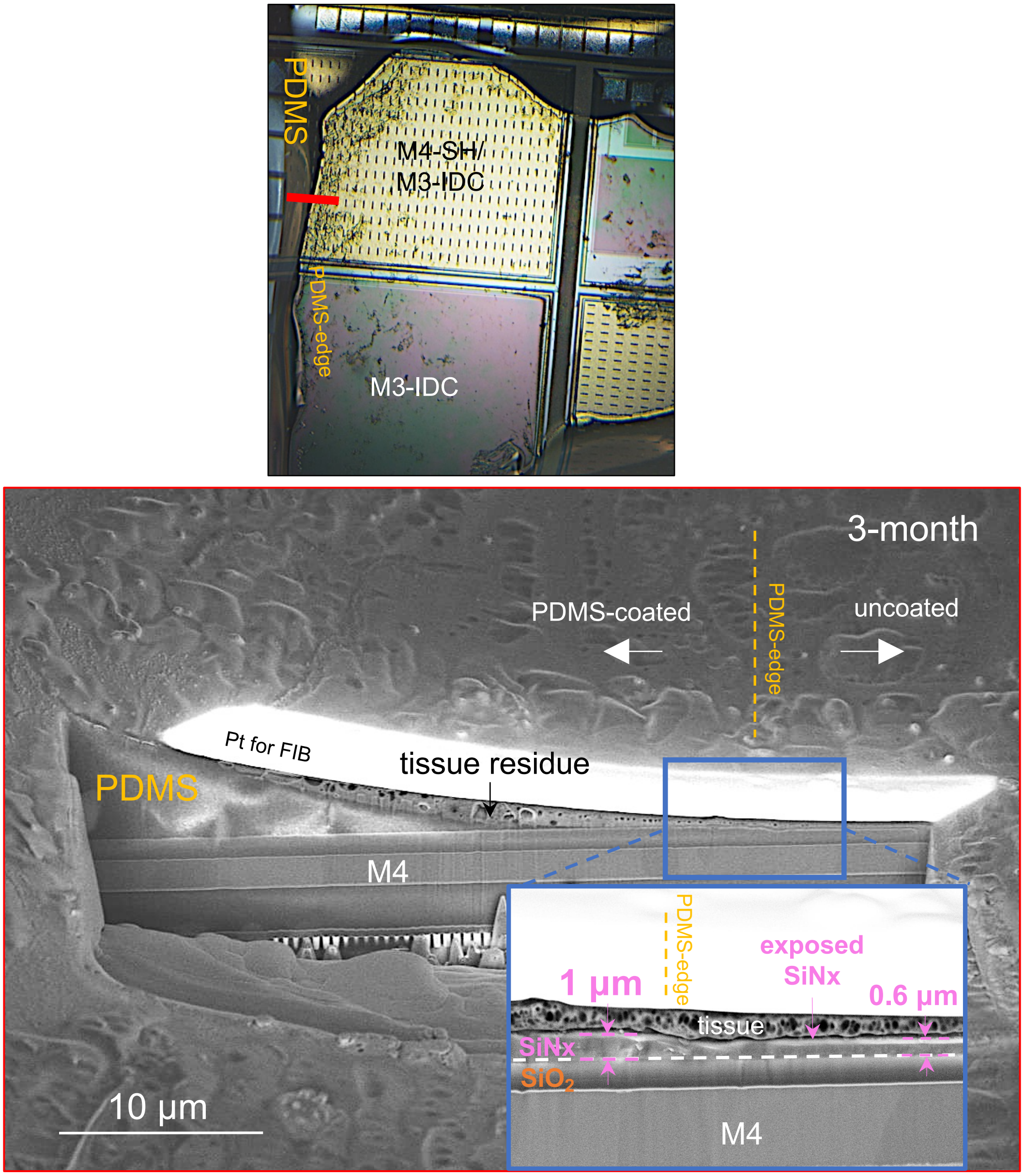


**Figure S16**. Optical micrograph of a Chip-A sample explanted after three months in rat, before tissue removal (top). Red line indicates FIB cut used for cross section analysis. Cross sectional SEM image near the PDMS-edge of the M4-SH/M3-IDC test structure showing the IC material stack (bottom). Inset: magnified SEM image of the cross-section near the PDMS-edge, comparing the top SiNx and $SiO_2$ passivation layers in the PDMS-coated and uncoated regions. Dissolution of the SiNx passivation in the uncoated region is visible, while no dissolution is observed in the PDMS-coated region. The thickness of the PDMS coating near the PDMS-edge is less than 1 µm. The remaining IC material stack (IMD and metallization) show no signs of delamination or degradation and remain intact.

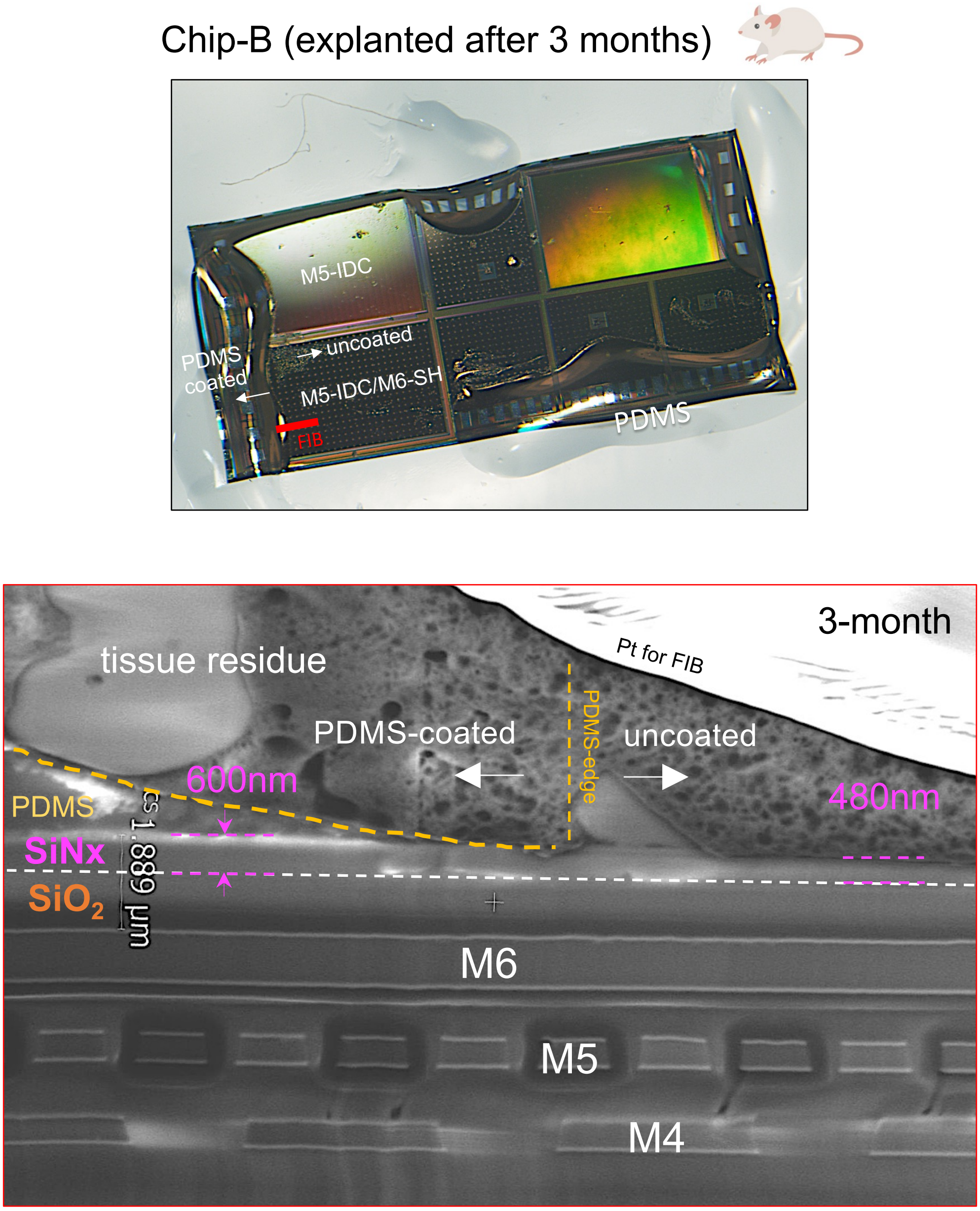


**Figure S17**. Optical micrograph of an explanted Chip-B sample after 3 months exposure to body environment, (before tissue removal). SEM cross-sectional image on the PDMS-edge of M5-IDC/M6-SH test structure show visible dissolution of SiNx passivation in the uncoated region. The remaining IC material stack (IMD and metallization) show no signs of delamination or degradation and remain intact.

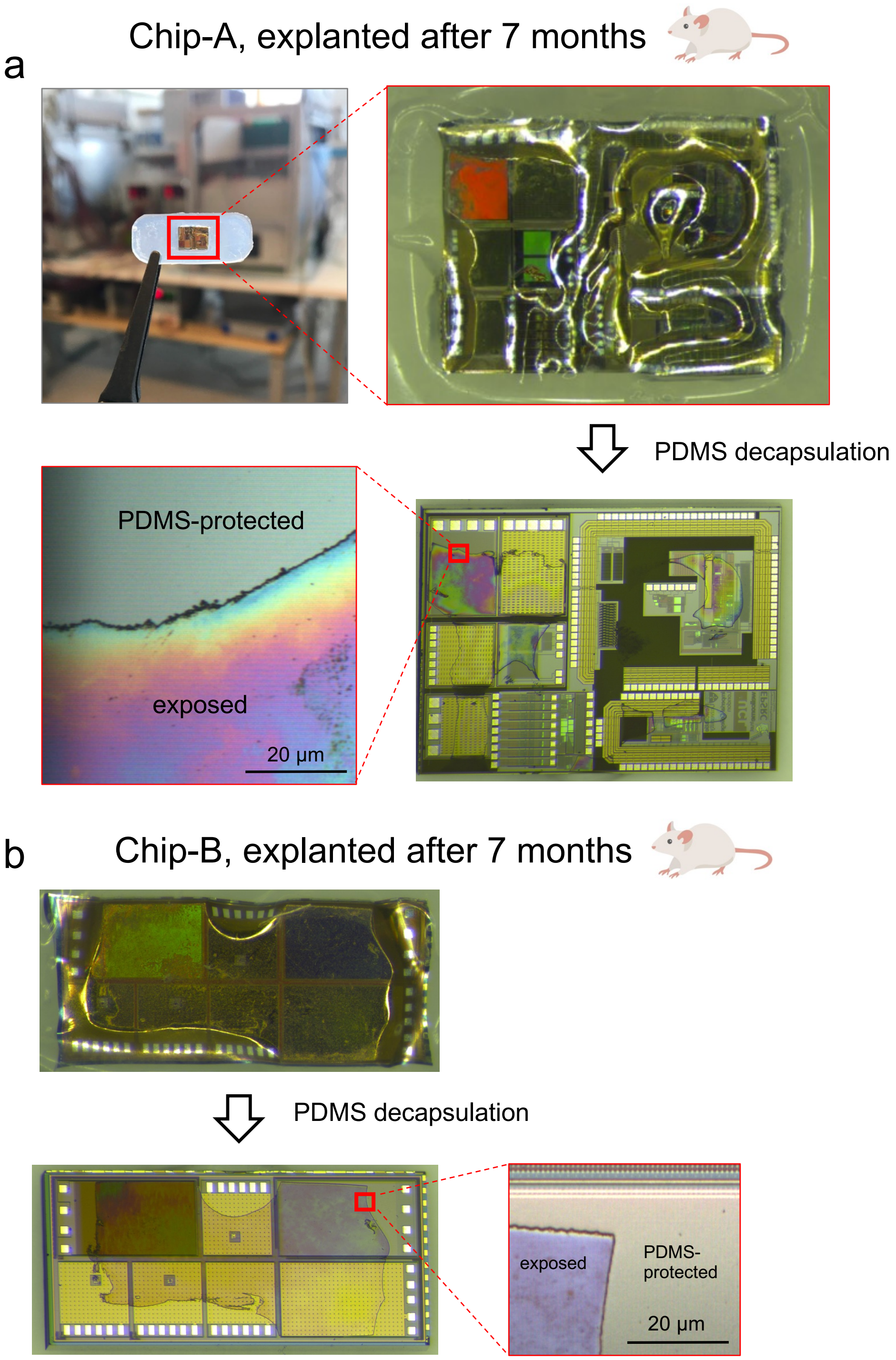


**Figure S18**. Representative optical micrographs of 7-month explanted chips. **a)** optical microscopic image of a Chip-A sample explanted after 7 months of implantation in rat, before and after PDMS decapsulation. Difference in color is noticeable between the PDMS protected and exposed regions which is due to the *in vivo* degradation of the SiNx passivation. **b)** Micrograph of a Chip-B sample explanted after 7 months, before and after PDMS decapsulation. Note that in both chips, the PDMS-coated aluminum pads remain intact.

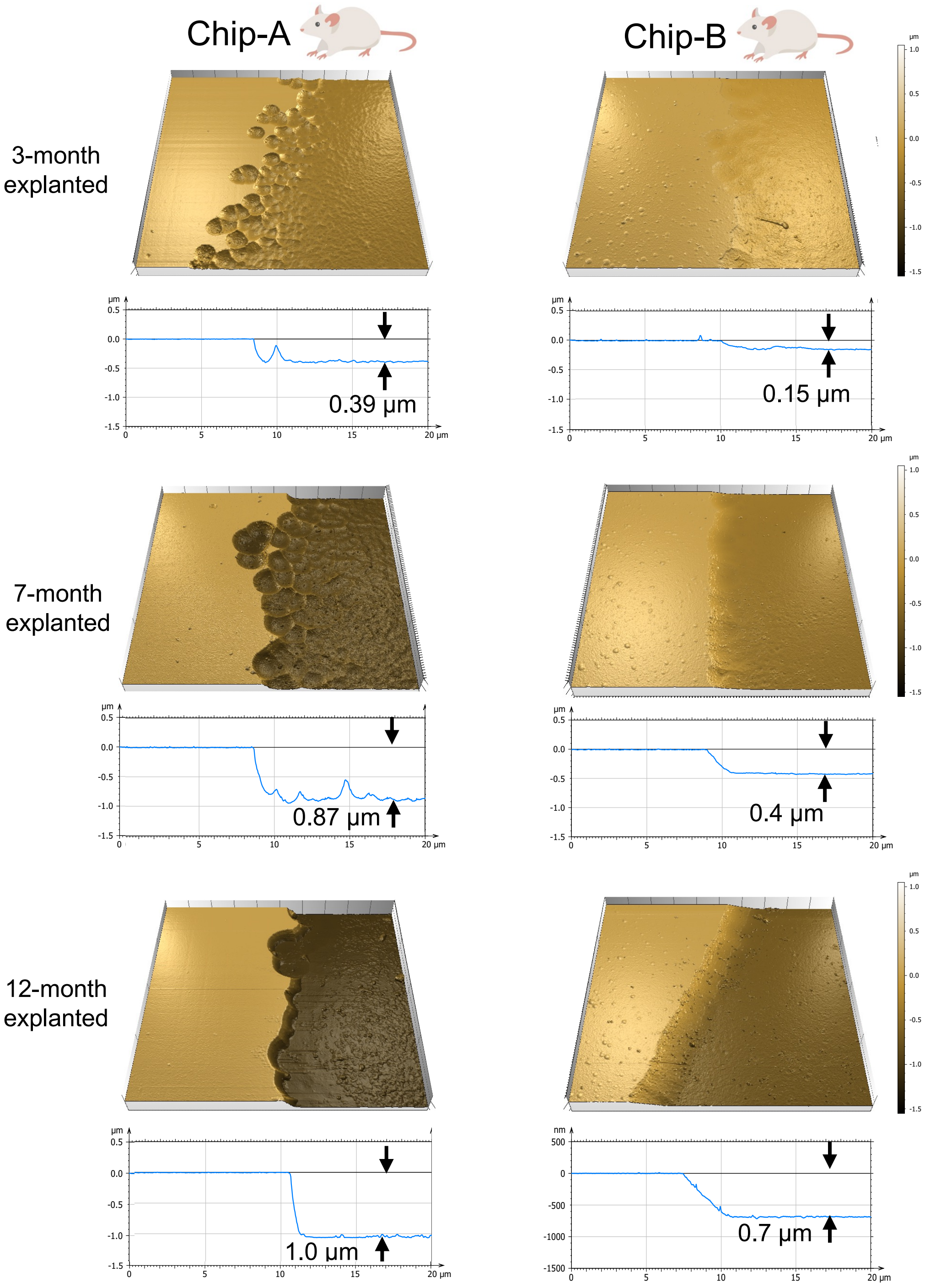


**Figure S19**. AFM surface topography analyzed on a 20 µm × 20 µm area at the PDMS-edge of decapsulated Chip-A and Chip-B ICs explanted at different time points showing gradual dissolution of the SiNx layer *in vivo*.

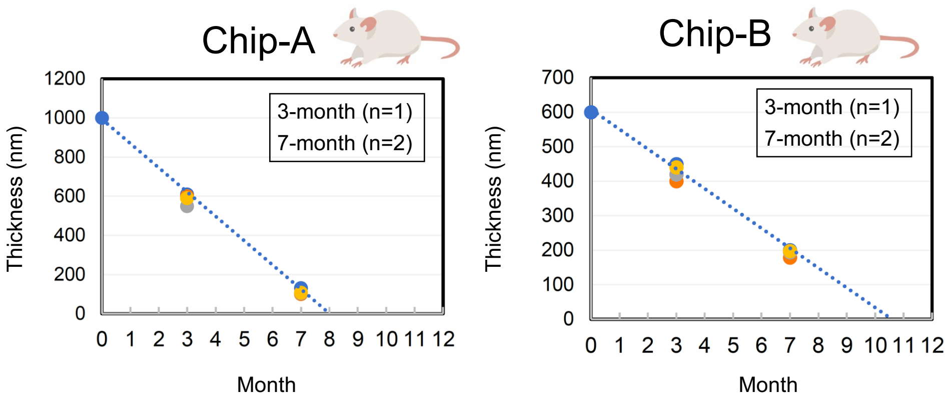


**Figure S20**. *In vivo* SiNx passivation loss over time for Chip-A and B ICs. Measurements were done on two explanted ICs at 3 and 7-month time points. Each IC was measured on two different locations. Extrapolated line (blue dashed line) shows at month 8 and 10.5, the entire SiNx passivation will have dissolved for Chip-A and B, respectively.

#### Biocompatibility of Silicon-IC passivation layers: SiNx and $SiO_2$ 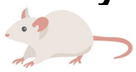

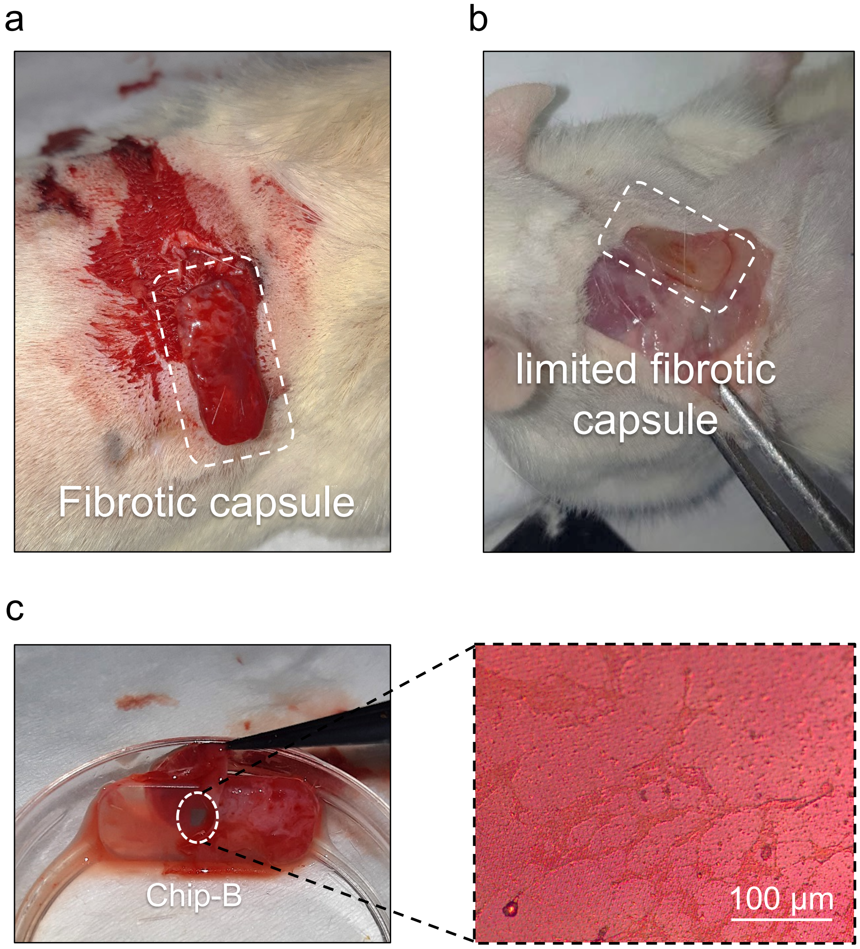


**Figure S21**. Images of explanted ICs. **a)** Tissue pocket formation around an implanted sample (Chip-A) after three months of implantation. **b)** Explanted sample with limited tissue pocket formation around the samples after 7 months of implantation where the PDMS substrate is clearly visible. **c)** Easy tissue pocket removal from a 7-month explanted Chip-B IC showing no tissue adhesion to the IC passivation surface or the surrounding PDMS. Optical micrograph of the passivation surface showing fibroblasts covering the SiN (left).

Before explanation, the skin around the implants was examined for any inflammation.

#### Biocompatibility of Silicon-IC passivation layers: SiNx and $SiO_2$

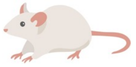

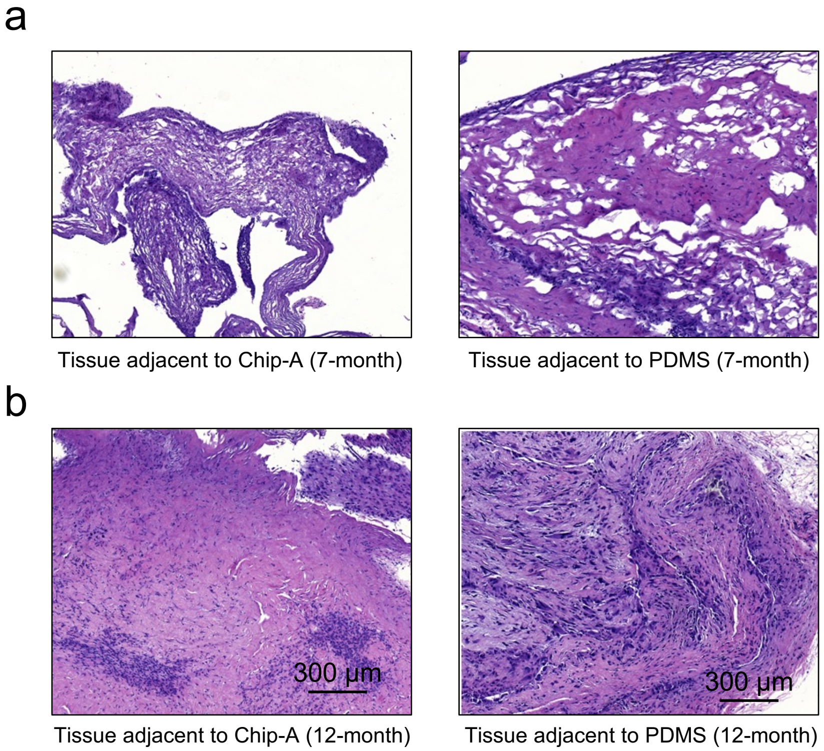


**Figure S22.** Histology comparing the tissue adjacent to the silicon-IC (Chip-A) with tissue adjacent to the PDMS substate (used as control). **a**) Histology of a 7-month explanted Chip-A. At 7 months, the surface of the IC is SiNx. **b**) Histology of a 12-month explanted Chip-A. For Chip-A, at month 12, the $SiO_2$ passivation is exposed to tissue for approximately 4 months.

All tissue samples were stained with hematoxylin-eosin stain for visualizing the cell nuclei and the cytoplasm. Qualitative analysis of the stained samples showed mature fibrotic tissue with mast cell infiltration in 12-month implants, whereas less mature, more cellular tissue was seen in the case of 7-month-old implants for both PDMS and chip implants.

Hämmerle et al. [4], have previously studied the long-term (6-12 months) biocompatibility of SiNx/$SiO_2$ layers in the rabbit eye using histology analysis. Other works have followed a similar approach [Voskerician Biomaterials 2003]. Here, our results also indicated no inflammation or tissue damage after 1-year of subdermal implantation of two CMOS foundry ICs in rats. These results are despite the observed SiN dissolution for ICs from foundries.

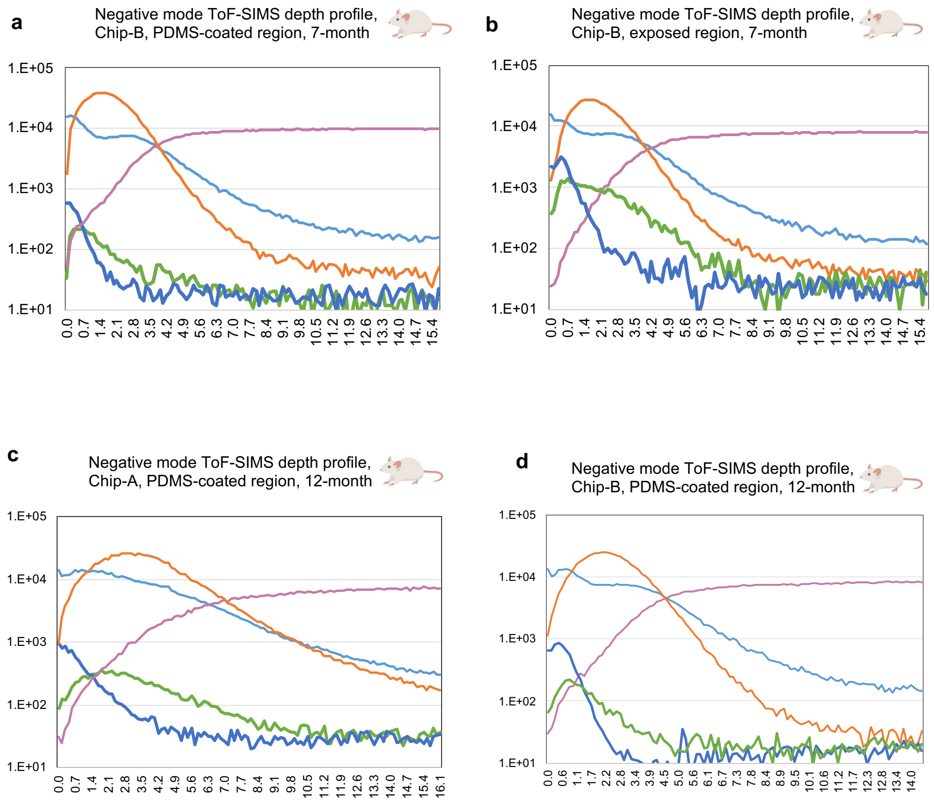


**Figure S23**. Negative mode shallow ToF-SIMS depth profiles acquired with sub-nanometer step-size from 0 - 15 nm of PDMS-coated and exposed (uncoated) regions of 7-month and 12-month explanted ICs.

a Chip-A (15V biased)

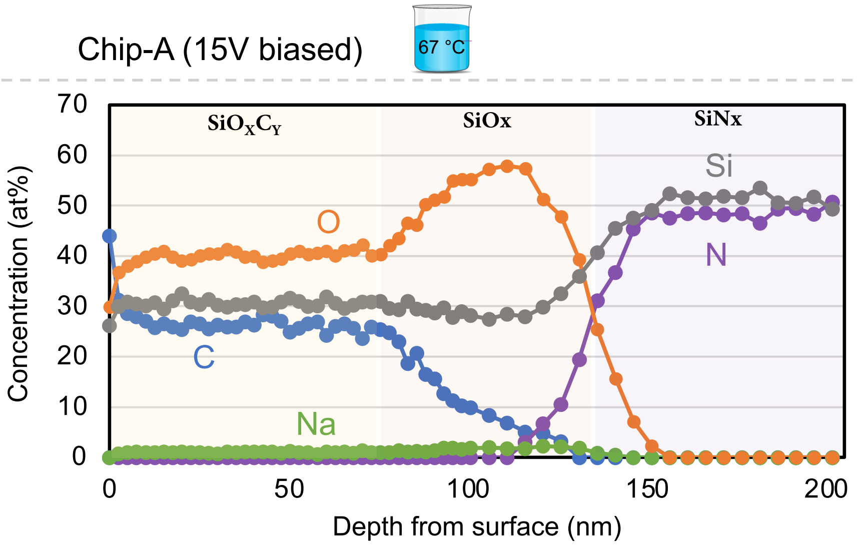


| | | Si | N | O | C | Na | Cl | K |
|---|---|---|---|---|---|---|---|---|
| $SiO_XC_Y$ | ~80 nm | 31.4 | 0 | 40.5 | 25.6 | 0.8 | 0 | 0 |
| SiOx | ~60 nm | 28 | 0 | 58 | 6.8 | 1.8 | 0 | 0.6 |
| SiNx | | 50 | 50 | 0 | 0 | 0 | 0 | 0 |

b Chip-B (15 V biased)

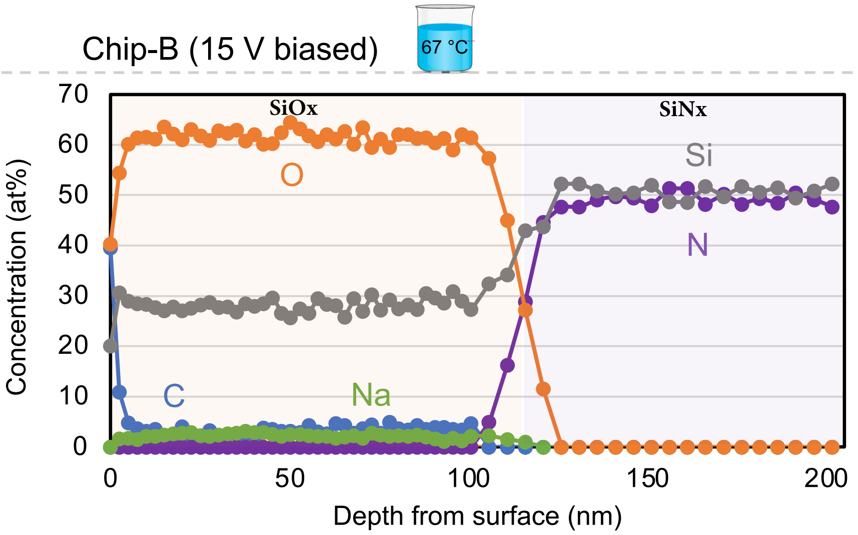


| | | Si | N | O | C | Na | Cl | K |
|---|---|---|---|---|---|---|---|---|
| $SiO_X$ | ~116 nm | 27.9 | 0 | 62.1 | 1.6 | 2.9 | 0 | 0.5 |
| SiN | | 50 | 50 | 0 | 0 | 0 | 0 | 0 |

**Figure S24**. XPS depth profiling giving chemical composition of the first 200 nm of the SiNx passivation layer after 12 months of continuous electrical biasing with 15 V DC in PBS solution at 67 °C. **a)** Chemical composition of Chip-A when biasing between M3-IDC (negative) and PBS (positive). **b)** Chemical composition of Chip-B when between biasing M5-IDC (negative) and PBS (positive).

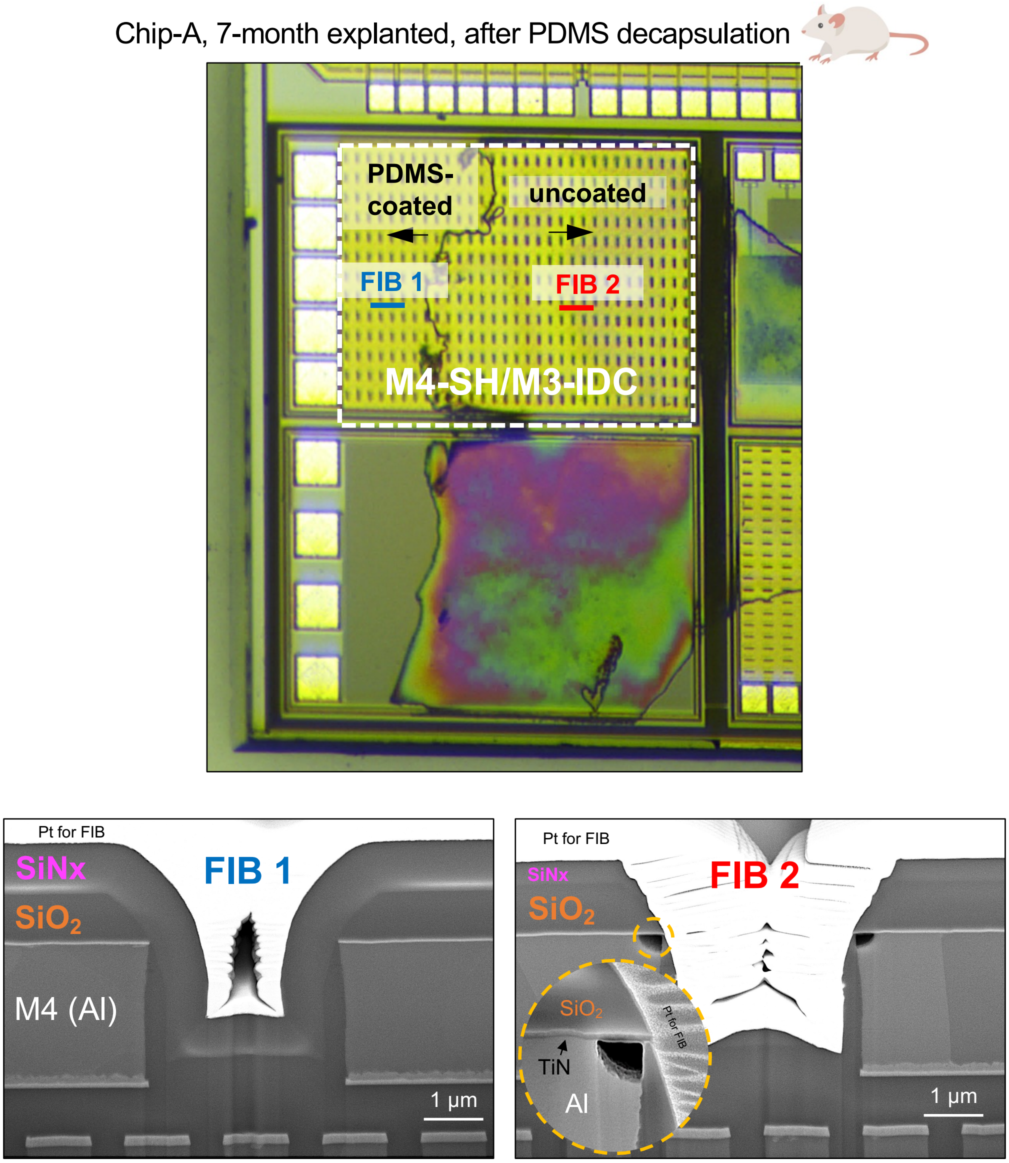


**Figure S25**. Optical micrograph and SEM cross section images of a Chip-A sample explanted at month 7 and PDMS decapsulated. Blue and red lines in the M4-SH/M3-IDC structure indicate FIB cuts in the PDMS-coated and exposed (uncoated) regions, respectively, that are used for cross sectioning. SEM cross-sectional images of the FIB cuts compare the PDMS-coated (FIB 1: bottom left) and exposed (FIB 2: bottom right) regions of the SH-M4/M3-IDC structure. In the exposed (uncoated) region, corrosion of the Metal-4 (top metal) layer is visible due to the dissolution of the SiNx passivation and the poor conformality of the $SiO_2$ passivation layer. Note on top of the IC, in the flat area, a thinned SiNx is still present. The sides, however, due to poor conformality, have a thinner SiNx. Inset shows a high magnification of the corroded aluminum metal while the TiN layer remain intact. The poor conformality of the $SiO_2$ is a result of using the top metallization (in this case M4).
